# The role of BST4 in the pyrenoid of *Chlamydomonas reinhardtii*

**DOI:** 10.1101/2023.06.15.545204

**Authors:** Liat Adler, Chun Sing Lau, Kashif M. Shaikh, Kim A. van Maldegem, Alex L. Payne-Dwyer, Cecile Lefoulon, Philipp Girr, Nicky Atkinson, James Barrett, Tom Z. Emrich-Mills, Emilija Dukic, Michael R. Blatt, Mark C. Leake, Gilles Peltier, Cornelia Spetea, Adrien Burlacot, Alistair J. McCormick, Luke C. M. Mackinder, Charlotte E. Walker

## Abstract

In many eukaryotic algae, CO2 fixation by Rubisco is enhanced by a CO2- concentrating mechanism, which utilizes a Rubisco-rich organelle called the pyrenoid. The pyrenoid is traversed by a network of thylakoid-membranes called pyrenoid tubules, proposed to deliver CO2. In the model alga *Chlamydomonas reinhardtii* (**Chlamydomonas**), the pyrenoid tubules have been proposed to be tethered to the Rubisco matrix by a bestrophin-like transmembrane protein, BST4. Here, we show that BST4 forms a complex that localizes to the pyrenoid tubules. A Chlamydomonas mutant impaired in the accumulation of BST4 (***bst4***) formed normal pyrenoid tubules and heterologous expression of BST4 in *Arabidopsis thaliana* did not lead to the incorporation of thylakoids into a reconstituted Rubisco condensate. Chlamydomonas *bst4* mutant did not show impaired growth at air level CO2. By quantifying the non-photochemical quenching (**NPQ**) of chlorophyll fluorescence, we show that *bst4* displays a transiently lower thylakoid lumenal pH during dark to light transition compared to control strains. When acclimated to high light, *bst4* had sustained higher NPQ and elevated levels of light-induced H2O2 production. We conclude that BST4 is not a tethering protein, but rather is an ion channel involved in lumenal pH regulation possibly by mediating bicarbonate transport across the pyrenoid tubules.

**One-sentence summary:** In Chlamydomonas, the pyrenoid-localized bestrophin-like protein BST4 is a putative ion channel involved in pH regulation of the thylakoid lumen, possibly by mediating bicarbonate transport.

## INTRODUCTION

Maintaining improvement in crop yields to keep pace with the rising demands for food is becoming increasingly challenging (Horton et al., 2021). Current models predict that an increase in food supply between 35 and 56% from 2010 to 2050 is required (van Dijk et al., 2021). A possible solution to overcome this challenge is engineering a biophysical CO2-concentrating mechanism (**CCM**) into C3 crop plants, which has been proposed to improve crop yields by between 8 and 60%, as well as water-use and nitrogen-use efficiency (Price et al., 2013; McGrath and Long, 2014; Long et al., 2019; Fei et al., 2022; Wu et al., 2023). The biophysical CCMs in algae typically function by concentrating CO2 into a liquid-liquid phase separated microcompartment called a pyrenoid, which is predominantly made up of a Ribulose-1,5-bisphosphate carboxylase/oxygenase (**Rubisco)**-rich matrix. This raises the [CO2]:[O2] ratio around the primary CO2-fixing enzyme Rubisco, which brings Rubisco closer to its maximal carboxylation rate and minimizes the competing oxygenation reaction. *Chlamydomonas reinhardtii* (hereafter **Chlamydomonas**) has the most well understood pyrenoid-based CCM and has become the blueprint for engineering such a CCM into C3 plants (Hennacy and Jonikas, 2020; Adler et al., 2022).

An important yet little understood aspect of the Chlamydomonas CCM is the function and biogenesis of the thylakoid tubule network that traverses the pyrenoid, known as the pyrenoid tubules. The pyrenoid tubules are continuous with the thylakoid membrane (which harbors the photosynthetic electron transport chain) (Engel et al., 2015) and are thought to function as a delivery system for inorganic carbon (**Ci**) to the Rubisco-rich pyrenoid matrix (Mitra et al., 2005; Raven 1997). In the current model, bicarbonate (**HCO3^-^**) is channelled into the thylakoid lumen by bestrophin-like proteins 1-3 (**BST1-3**) (Mukherjee et al., 2019) and diffuses to the pyrenoid tubules where it is converted to CO2 by carbonic anhydrase 3 (**CAH3**) (Karlsson et al., 1998; Mitra et al., 2005) thanks to a low lumenal pH generated by the photosynthetic electron transport chain (Burlacot et al., 2022). Traversion of the pyrenoid Rubisco-matrix by tubules is predicted to be essential for an efficient CCM (Fei et al., 2022). Therefore, understanding the mechanisms of pyrenoid tubule formation and function will be crucial for future plant pyrenoid engineering efforts.

The protein **BST4** (bestrophin-like protein 4, also known as Rubisco binding membrane protein 1, RBMP1; Cre06.g261750) localizes exclusively to the pyrenoid tubules and has been proposed to function as a tether protein, linking the Rubisco matrix to the tubules (Meyer et al., 2020). BST4 is a predicted transmembrane protein and has two Rubisco binding motifs (**RBMs**) on its long, disordered C-terminus (**Fig. 1A**) (He et al., 2020; Meyer et al., 2020). RBMs facilitate the targeting of proteins to the pyrenoid and are also hypothesized to underpin the assembly of the pyrenoid. Meyer et al. (2020) proposed that BST4, together with other tether proteins, may recruit Rubisco to the tubule network.

**Figure 1.**
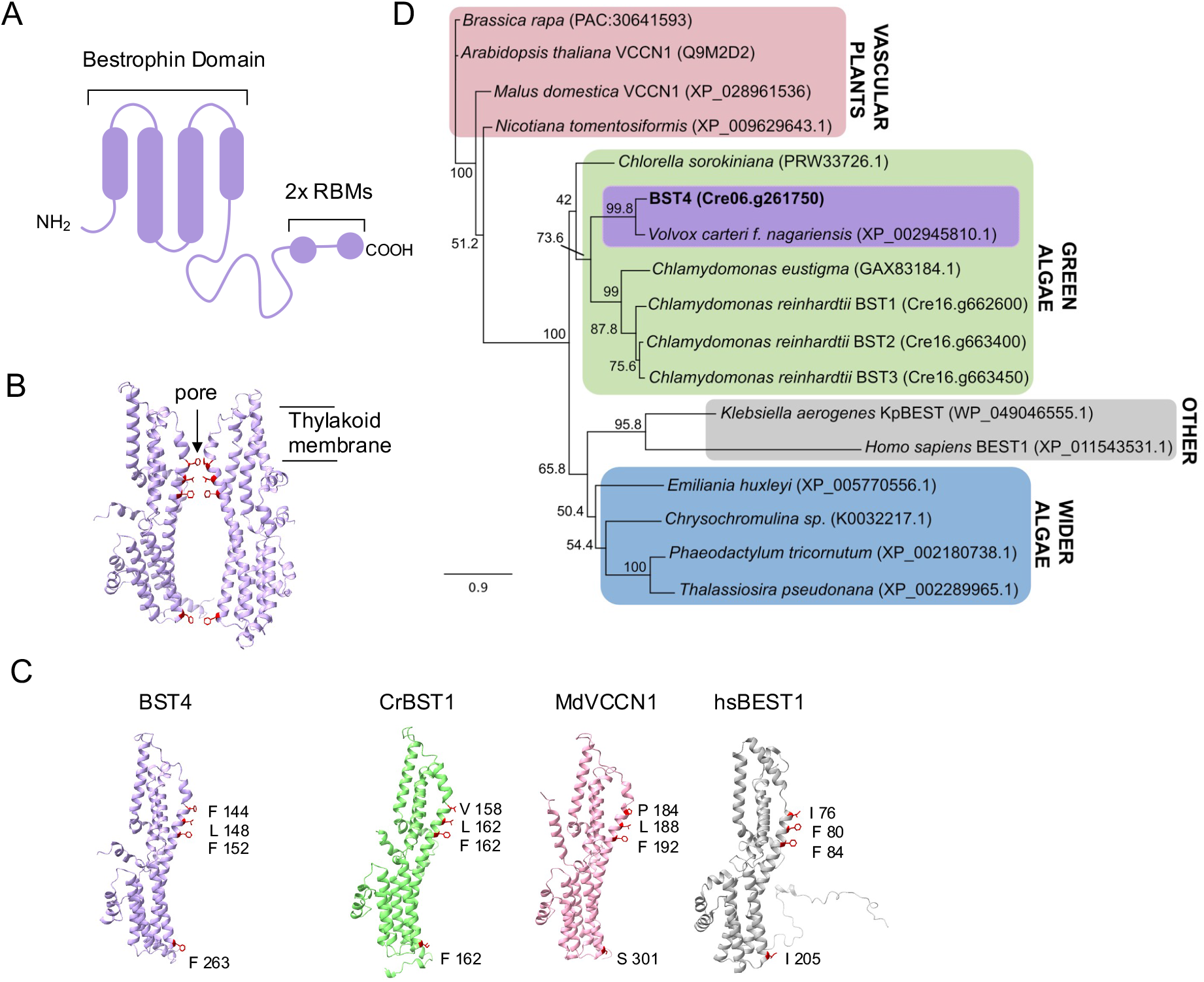
BST4 is a bestrophin protein that is distinct from BST1-3. **A.**Schematic of the topology of BST4. BST4 is predicted to have four transmembrane domains and a disordered C-terminus containing two Rubisco Binding Motifs (RBMs) **B.** AlphaFold v2 models of two BST4 bestrophin domains (amino acid residues 53-386 shown for clarity) to show a typical bestrophin channel pore. Pore lining residues are highlighted in red. **C.** AlphaFold v2 structure of the BST4 bestrophin domains alongside the predicted structure of another bestrophin-like protein from *Chlamydomonas reinhardtii* (*Cr*BST1; Alphafold amino acids 51-end shown for clarity), and experimentally determined structures of bestrophins *Malus domestica* voltage-dependent Cl^−^ channel 1 (*Md*VCCN1;7EK1) and *Homo sapiens* Bestrophin 1 (*Hs*BEST1; 8D1I). Residues known to line the channel pore are highlighted. **D.** Phylogenetic analysis of the BST4 bestrophin domain (bold) with the disordered C-terminal removed. The alignment used was trimmed at residue 369. The evolutionary history of BST4 was inferred by using the maximum likelihood method based on the Le and Gascuel substitution model with discrete Gamma distribution (5 categories) and 500 bootstrap replicates. The tree is drawn to scale, with branch lengths measured in the number of substitutions per site.

BST4 also has a well-conserved bestrophin domain similar to those present in the thylakoid-localized BST1-3 proteins (Mukherjee et al. 2019). Bestrophins primarily act as anion channels and are found in a wide diversity of organisms, including animals, plants and fungi. They are best known to have permeability to chloride and HCO3^-^ (Qu and Hartzell, 2008), although some are reportedly permeable to cations (Yang et al., 2014) and larger organic anions (Roberts et al., 2011). It is possible that BST4 functions as an HCO3^-^ channel, like that proposed for BST1-3 (Mukherjee et al. 2019), but there are currently no data to support this hypothesis.

In this study we aimed to elucidate the role of BST4 in the Chlamydomonas CCM. We tested BST4 as a thylakoid-Rubisco tethering protein as well as its suitability in promoting a thylakoid-Rubisco matrix interface in the model land plant Arabidopsis. We also explored an alternative hypothesis that BST4 functions as an ion channel in the pyrenoid tubules and discussed the implications this has for our current understanding of CO2 fixation by the pyrenoid.

## RESULTS AND DISCUSSION

### BST4 is a bestrophin-like protein that is localized in the intra-pyrenoid thylakoid tubules in Chlamydomonas

The amino acid sequence of BST4 has two key unique features compared to the previously characterized BST1-3 (**Fig. 1**) (Mukherjee et al. 2019). First, BST4 has an extended disordered C-terminus that contains two RBMs (**Fig. 1A**) (He et al., 2020; Meyer et al., 2020). Second, BST4 has a phenylalanine residue in the first position of the putative selection pore, as opposed to valine which is conserved throughout BST1- 3 (Mukherjee et al. 2019), although the residue at this position is variable in other well characterized bestrophin proteins (*Malus domestica* voltage-dependent Cl^−^ channel 1 (***Md*VCCN1**) and *Homo sapiens Bestrophin 1* (***Hs*BEST1**)) (**Fig.1B, C**). In corroboration, we found the evolutionary history of BST4 diverges from BST1-3 when investigated using maximum likelihood phylogenetic analysis. We analysed the full-length sequence (**Fig. S1**) and a truncated version without the disordered C-terminal, leaving the bestrophin domain and N-terminal (**Fig. 1D**), both of which found BST4 to resolve in a distinct clade from BST1-3 but within the wider green algae group.

As well as being distinct at a sequence level, BST4 also localizes differently from BST1-3 (**Fig. 2**). While BST1-3 localize throughout the thylakoid membrane and are enriched at the pyrenoid periphery (Mukherjee et al., 2019), BST4 localizes to the centre of the pyrenoid in a pattern that resembles the pyrenoid thylakoid tubule system (**Fig. 2A, B**; Meyer et al., 2020). To confirm tubule localization, we generated a dual-tagged line that expressed BST4-mScarlet-I and the Chlamydomonas Rubisco small subunit 1 (***Cr*RBCS1**) fused to Venus. BST4-mScarlet-I was enriched where *Cr*RBCS1-Venus was depleted (**Fig. 2C, D**), indicating that BST4 is located in the tubules and not the Rubisco-enriched pyrenoid matrix. Previous work suggests that the C-terminal RBMs of BST4 enable the protein to interact with *Cr*RBCS1 (Meyer et al. 2020). We confirmed using a yeast-2-hybrid approach that the C-terminus of BST4 interacts with *Cr*RBCS1 (**Fig. S2A**). We also measured the efficiency of Förster resonance energy transfer (**FRET**) from Venus to mScarlet-I and found that the FRET efficiency was ∼35%, supporting the proximity of BST4 to *Cr*RBCS1 (**Fig. S2B**).

**Figure 2.**
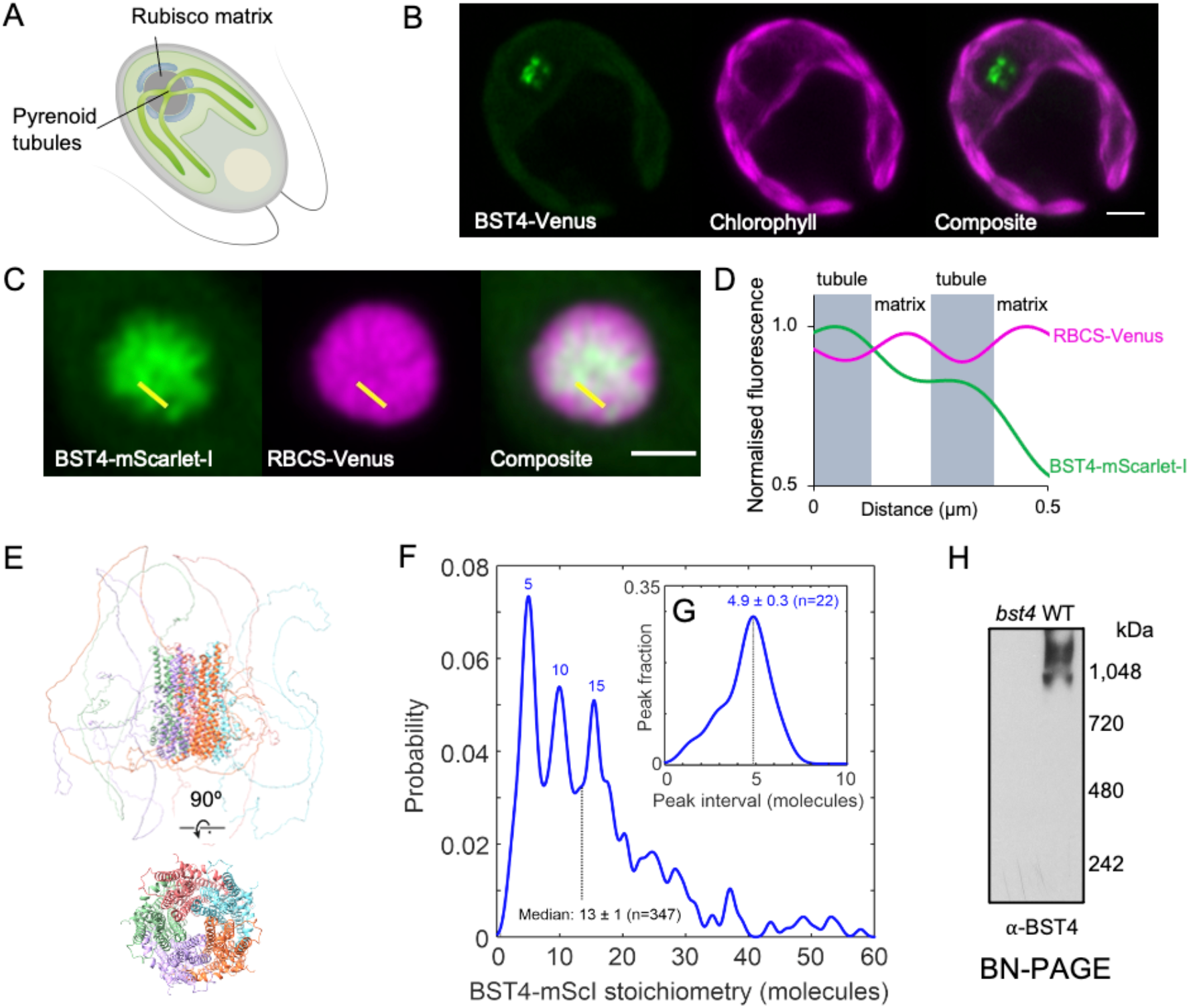
BST4 forms higher order assemblies in the pyrenoid tubules in Chlamydomonas. **A**. Diagram of a Chlamydomonas cell with the pyrenoid Rubisco matrix and pyrenoid tubules indicated. **B.** Confocal image of a Chlamydomonas cell expressing BST4-Venus. Scale bar is 2 µm. **C.** Pyrenoid in dual-tagged Chlamydomonas. BST4-mScarlet-I and RBCS-Venus are shown in green and magenta, respectively. Overlap appears white. The yellow line shows the 1D cross-section used for generating the line plot in (D). Scale bar is 1 µm. **D**. Plot of normalized fluorescence intensity values from a 1D cross-section from (C). mScarlet-I and RBCS-Venus are shown in green and magenta, respectively. **E**. AlphaFold Multimer v2 prediction of pentameric BST4 structure. Top structure includes the disordered C-terminus, bottom structure displays amino acid residues 53-386 only for clarity. **F**. BST4-mScarlet-I stoichiometry probability distribution based on single particle tracking and molecular counting in live Chlamydomonas (n=347 tracks) using Slimfield microscopy. **G**. Averaging the intervals between peaks in BST4-mScarlet-I stoichiometry (n=22 intervals) indicates a consistent pentameric unit. **H**. Immunoblot of proteins from *bst4* mutant and wild type (WT) Chlamydomonas thylakoids separated by Blue Native-PAGE.

Bestrophins typically form pentameric assemblies (Bharill et al., 2014; Hagino et al., 2022). When five chains of BST4 were inputted to AlphaFold a typical bestrophin pentameric structure was predicted (**Fig. 2E**). To test the complex assembly of BST4 *in vivo*, we utilized a Slimfield microscopy molecular tracking method (Plank et al., 2009). For this method, it is important to have only fluorescently tagged BST4 molecules and no native (untagged) BST4 molecules. To find a strain that was impaired in the accumulation of BST4 protein, we screened *bst4* insertional mutants from the CLiP library (Zhang et al., 2014; Li et al., 2019) and confirmed a mutant strain (***bst4***) (**Fig. S3** and methods). We then expressed BST4-mScarlet-I in the *bst4* mutant background (**Fig. S3**). We used Slimfield microscopy to image BST4 in the pyrenoid tubules and subsequent image analysis (detailed in methods) to track individual fluorescent molecules of BST4-mScarlet-I and quantified the number of BST4 monomers per complex. The resulting probability distribution revealed that the most common BST4 complex is made up of five molecules (**Fig. 2F**). Other peaks showed complexes with numbers of molecules divisible by five, which may be multiple pentameric channels grouping together. The interval between the probability peaks was also five (**Fig. 2G**). To further support higher order complex assembly of BST4 we ran purified Chlamydomonas thylakoid membranes on a Blue Native-PAGE gel (**BN-PAGE**) and immunodetected BST4 (**Fig. 2H**). BST4 formed a smear at ∼1000 kDa, which is considerably larger than a pentamer (∼330kDa). This could be due to higher order assemblies of BST4, pentameric BST4 in a complex with other proteins and/or aberrant migration during BN-PAGE due to influences by complex shape. Collectively, our data support *in vivo* higher order assembly of BST4 potentially as a pentamer.

### BST4 localizes to the stroma lamellae thylakoid membrane in Arabidopsis

We used *Arabidopsis thaliana* (**Arabidopsis** hereafter) as a heterologous system to examine if BST4 was able to tether the Rubisco matrix to thylakoid membranes. In previous work, an Arabidopsis Rubisco small subunit (*At*RBCS) double mutant (*1a3b*) was complemented with *Cr*RBCS2, resulting in a line with hybrid Rubisco representing ∼50% of the Rubisco pool (**S2Cr**) (Izumi et al., 2012; Atkinson et al., 2017). Subsequent expression of the pyrenoid linker protein Essential PYrenoid Component 1 (**EPYC1**), in S2Cr resulted in the formation of an EPYC1-hybrid Rubisco condensate or ‘proto-pyrenoid’ (Atkinson et al., 2020). The S2Cr line was therefore used as a platform to test whether BST4 acts as a tether protein.

We initially localized a BST4-mNeon fusion protein following stable expression in S2Cr (**Fig. 3**). BST4-mNeon was observed in chloroplasts, demonstrating that the native chloroplast signal peptide was compatible with the chloroplast targeting mechanism in land plants (**Fig. 3A**), as seen previously for other Chlamydomonas proteins (Atkinson et al., 2016). The fluorescence signal from BST4-mNeon had a sponge-like pattern that was inversely correlated with the punctate chlorophyll autofluorescence signal that represents the grana stacks (**Fig. 3B, C**). The sponge-like pattern was similar to previous observations of autofluorescence originating from Photosystem (**PS**) I (Hasegawa et al., 2010), which is enriched in the stroma lamellae of thylakoids. We subsequently generated a stable Arabidopsis transgenic line expressing untagged BST4 to confirm its location by biochemical fractionation. BST4 was detected in the thylakoid fraction and not in the stromal fraction (**Fig. 3D**). The thylakoids were then further fractionated into grana stacks and stroma lamellae sub-fractions. BST4 was found in the stroma lamellae fraction (**Fig. 3E**), which is consistent with the observed sponge-like fluorescence pattern.

**Figure 3.**
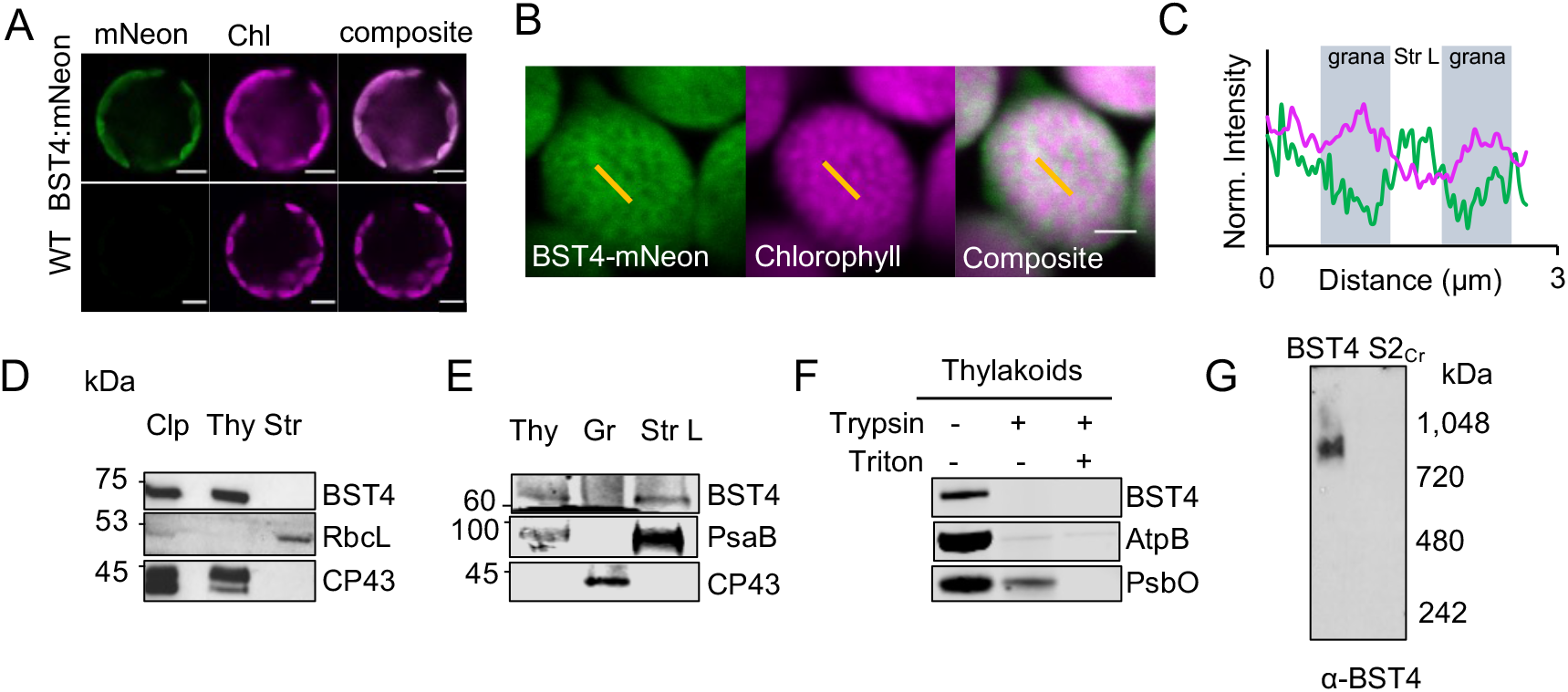
BST4 assembles as a complex in the stroma lamellae of thylakoids in Arabidopsis. **A.**Confocal image of WT protoplasts expressing BST4-mNeon. Scale bar is 5 µm. **B.** Mesophyll chloroplast from S2Cr Arabidopsis expressing BST4-mNeon. BST4mNeon and chlorophyll autofluorescence are shown in green and magenta, respectively. Overlap appears white. Scale bar is 2 µm. Yellow lines indicate selections for profile plot in (C). **C.** Plot of normalized fluorescence intensity values from a 1D cross-section (yellow line) through two grana stacks from (B). mNeon and chlorophyll autofluorescence are shown in green and magenta, respectively. **D.** Immunoblots of sub-chloroplast fractions isolated from Arabidopsis line S2Cr expressing BST4. RbcL and CP43 were probed for as stromal and thylakoid controls, respectively. **E.** Immunoblots of fractionation thylakoids from Arabidopsis line S2Cr expressing BST4. CP43 and PsaB were used for grana stack and stroma lamellae controls, respectively **F.** Trypsin protease protection assay. Intact thylakoids containing BST4 subjected to 0 or 100 μg/ml trypsin with or without the addition of 1% (v/v) Triton. AtpB and PsbO used as controls for stromal facing (exposed) and lumen facing (protected), respectively. **G.** Immunoblot of proteins from thylakoids separated by Blue Native-PAGE from either BST4 stable line or S2Cr background. Abbreviations: Clp, whole chloroplast; CP43, CP43-like chlorophyll binding protein; RbcL, Rubisco large subunit; PsbO, photosystem II manganese-stabilizing polypeptide;AtpB, ATP synthase subunit beta; PsaB, photosystem I P700 chlorophyll a apoprotein A2; Str, stromal fraction; Thy, thylakoid fraction.

As BST4 has two RBMs on its C-terminus, we hypothesized that BST4 should be orientated with the C-terminus facing the stroma so that the RBMs are available to interact with *Cr*RBCS. To determine the orientation of BST4, we performed a protease protection assay on thylakoids isolated from the untagged BST4 transgenic plants. Our antibody was raised against the C-terminal end of BST4, so it could be used to assess whether the C-terminus was exposed for degradation in the stroma or if it was protected in the lumen. We found that the BST4 C-terminus was fully degraded after a 60 min treatment of trypsin, indicating that it faced the stroma (**Fig. 3F**). There was some degradation of the lumenal control, the PSII subunit PsbO, which we attribute to a portion of the thylakoid membrane preparation not being fully intact. However, PsbO was fully degraded when the membranes were solubilized, indicating that they were sufficiently intact to differentiate between lumenal- and stromal-facing peptides. Therefore, BST4 was observed in the expected location and orientation in plant thylakoid membranes.

Finally, we also tested whether BST4 forms a complex in the Arabidopsis thylakoid membrane. We subjected thylakoids from the untagged BST4 transgenic plants to BN-PAGE and detected a single band of ∼850 kDa (**Fig. 3G**). Thus, BST4 forms a similar high order complex in Arabidopsis to that in Chlamydomonas but may be lacking additional interaction partners present in Chlamydomonas.

### BST4 is not sufficient for integration of thylakoids into the Arabidopsis proto-pyrenoid

To test whether BST4 could facilitate incorporation of thylakoids into hybrid Rubisco condensates (i.e., proto-pyrenoids) in plants, BST4-mCherry and EPYC1-tGFP were co-expressed in the S2Cr background (**Fig. 4**). When we expressed EPYC1 alone, Rubisco condensates formed in chloroplasts as previously described (Atkinson et al., 2020), and were visible as a ∼2 µm wide puncta in the tGFP channel (**Fig. 4A**). When BST4-mCherry and EPYC1-tGFP were co-expressed, approximately 60% of the BST4-mCherry fluorescence signal was observed in the condensate region (**Fig. 4B, D**), with the remaining signal exhibiting the same sponge-like pattern as seen in **Fig. 3A**.

**Figure 4.**
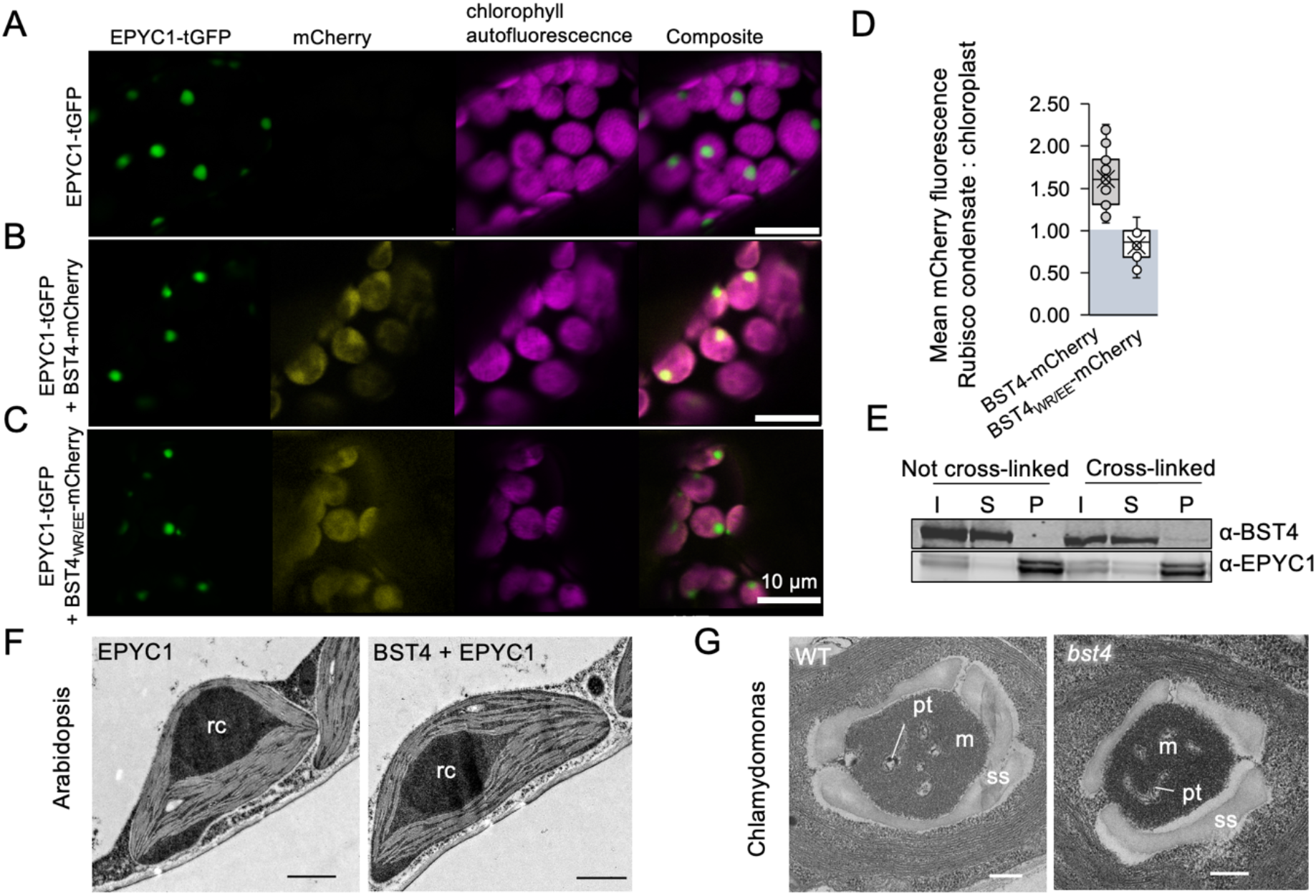
BST4 is not sufficient to enable the inclusion of thylakoid membranes in a Rubisco condensate in Arabidopsis and is not required for pyrenoid tubule inclusion in the Chlamydomonas pyrenoid. **A.**Confocal images showing EPYC1-tGFP in S2Cr Arabidopsis background. Scale bar is 10 µm. **B.** Confocal images showing BST4-mCherry co-expressed with EPYC1-tGFP in S2Cr Arabidopsis background. Scale bar is 10 µm. **C.** Confocal images showing BST4 with mutated RBMs (BST4(WR/EE)) fused to mCherry co-expressed with EPYC1- tGFP in the S2Cr Arabidopsis background. **D.** Ratio of mean mCherry fluorescence associated with the Rubisco condensate compares do the rest of chloroplast when mCherry was fused to either BST4 or BST4(WR/EE) (n=22- 23). **E.** Immunoblot of sedimented Rubisco condensates from S2Cr Arabidopsis expressing EPYC1-tGFP and BST4-mCherry. Abbreviations: I, input; S, supernatant; P, pellet. Rubisco condensates are enriched in the pelleted fraction. Cross-linked samples were prepared by vacuum infiltrating intact leaves with 1% (v/v) formaldehyde prior to sedimentation. **F.** Transmission electron micrograph of a chloroplast from S2Cr Arabidopsis expressing either EPYC1 alone or EPYC1-tGFP and BST4-mCherry. Rc is the Rubisco condensate. Scale bar is 1 µm. **G.** Transmission electron micrograph of pyrenoid from WT Chlamydomonas or *bst4* mutant. Abbreviations: pt, pyrenoid tubules; m, matrix; s, starch sheath. Scale bar is 250 nm.

To confirm that the observed co-localization was due to an interaction between BST4 and *Cr*RBCS2, we mutated the first two residues of each core RBM motif of BST4 to glutamic acid (WR to EE, the new version noted BST4WR/EE). Previously, these substitutions have been reported to disrupt the binding interface between EPYC1 and *Cr*RBCS2 (He et al., 2020). Using yeast-2-hybrid we confirmed that the interaction between the C-terminus of BST4 and *Cr*RBCS2 was disrupted by these mutations (**Fig. S3**). When BST4WR/EE-mCherry was expressed with EPYC1-tGFP in S2Cr, no enrichment of the mCherry signal was observed in the Rubisco condensate (**Fig. 4C, D**). Thus, the RBMs of BST4 were responsible for the enrichment of BST4 in the vicinity of the Rubisco condensate in Arabidopsis.

However, when condensates were sedimented and analysed by immunoblotting, BST4-mCherry was not detected in the condensate fraction (**Fig. 4E**). When leaf samples were subjected to formaldehyde cross-linking prior to sedimentation of the condensate, a small amount of BST4 was present in the condensate fraction. We concluded that BST4 was not present in the condensate itself but preferentially occupied the thylakoid membranes surrounding the *Cr*RBCS2-enriched condensate (Atkinson et al., 2020), likely due to interactions with *Cr*RBCS2.

Although BST4 partially co-localized with the Rubisco condensate, we found no evidence to suggest that BST4 could facilitate the inclusion of thylakoid membranes. Confocal microscopy was not sufficient to determine if chlorophyll autofluorescence was in the proto-pyrenoids of S2Cr lines expressing BST4-mCherry and EPYC1-tGFP. However, transmission electron microscopy (**TEM**) revealed no visible indication of thylakoid membranes in the condensates, which were structurally similar to condensates in S2Cr-EPYC1 lines lacking BST4 (**Fig. 4F**). Thus, BST4 appeared insufficient to enable thylakoid inclusion in the Rubisco condensate in Arabidopsis.

### BST4 is not necessary for tubule formation in Chlamydomonas

To investigate whether BST4 was necessary for normal formation of the thylakoid tubule-structure in Chlamydomonas we compared the structure of pyrenoids of *bst4* compared to the WT control strain (CMJ030) (Zhang et al., 2014; Li et al., 2019). TEM images showed that pyrenoids from *bst4* were structurally comparable to the WT control, including the presence of pyrenoid tubules (**Fig. 4G**). There were also no differences in pyrenoid size or shape between the two lines when comparing pyrenoids from 40-50 cells from each genotype (**Fig. S4**). As a result, we conclude that BST4 is not necessary for the pyrenoid tubule-Rubisco matrix interface in Chlamydomonas.

### The C-terminus of BST4 is required for localization to the pyrenoid tubules

Multiple copies of the RBM are sufficient to target proteins to the pyrenoid (Meyer et al., 2020). Furthermore, we have shown that the BST4 RBMs are required for RBCS interaction via yeast-2-hybrid (**Fig. S2A**) and that they are required for enrichment with the proto-pyrenoid in Arabidopsis (**Fig. 4B-D**). To investigate the role of the RBMs in BST4 localization in Chlamydomonas, we generated a truncated version of BST4 (residues 1 to 386) that lacked the C-terminus containing the two RBMs (BST4ΔC-term) and compared localization with full-length BST4 expressed in WT and the *bst4* mutant (**Fig. 5 and Fig. S5**). BST4ΔC-term-mScarlet expressed in the *bst4* mutant line did not localize to the pyrenoid tubules but was found throughout the thylakoid membrane (**Fig. 5**). This is consistent with our findings in Arabidopsis with BST4WR/EE (**Fig. 4C**), demonstrating that localization of BST4 is driven via RBM-Rubisco interaction. Thus, the C-terminus is necessary for BST4 localization to the pyrenoid tubules in the presence of the matrix. Collectively these data indicate that BST4 may require a pre-existing pyrenoid tubule network to be localized in the pyrenoid rather than driving the inclusion of thylakoid membranes into the Rubisco matrix or is redundant as a tether protein.

**Figure 5.**
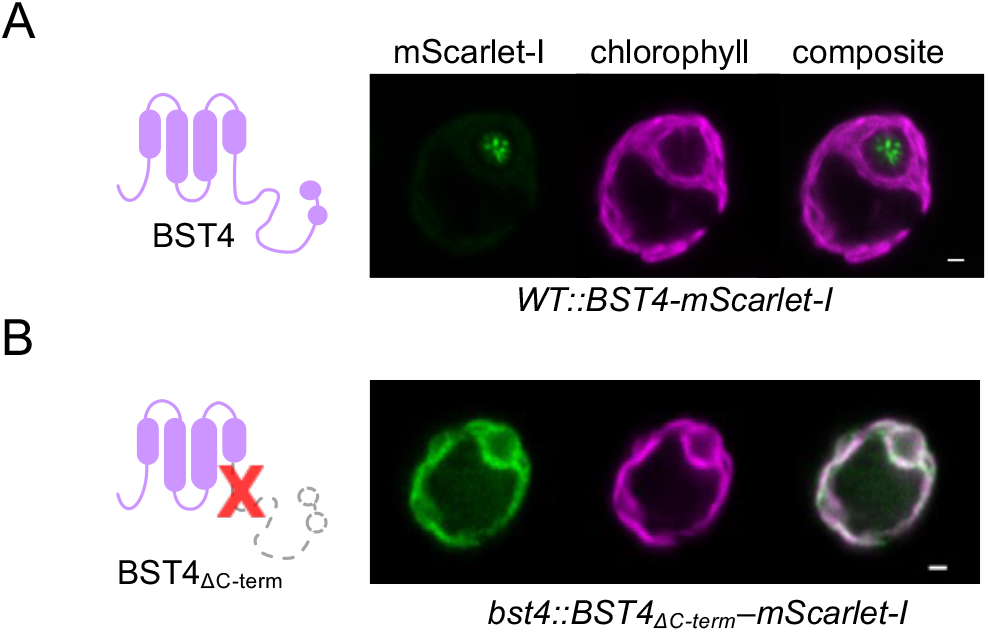
C-terminus of BST4 is required for pyrenoid localization. **A.**Confocal image of full length BST4- mScarlet-I in WT **B**. Confocal image of BST4ΔC-term-mScarlet-I in *bst4*. Scale bars are 1 µm.

When BST4ΔC-term-Venus was expressed in WT Chlamydomonas (i.e., that still produced non-truncated BST4), we observed fluorescence throughout the thylakoids, but with the majority of the signal still localized to the pyrenoid tubules (**Fig. S5**). We conclude that BST4ΔC-term-Venus is recruited to the pyrenoid through an interaction with the native full-length BST4, which is further evidence that BST4 oligomerizes.

We next investigated whether BST4 localizes to the pyrenoid tubules, or whether BST4 localizes to the tubules through an interaction with Rubisco. To do so, we utilized a Chlamydomonas mutant generated by Genkov et al. (2010) that expresses *At*RBCS but lacks both isoforms of *Cr*RBCS (*Crrbcs::AtRBCS*), and thus lacks a Rubisco matrix as EPYC1 does not interact with *At*RBCS (Atkinson et al., 2019). *Crrbcs::AtRBCS* retains reticulated thylakoid membranes at the canonical pyrenoid site, which are likely the nascent pyrenoid tubule network (Caspari et al., 2017) (**Fig. S6A**). Caspari et al. (2017) show large starch granules accumulating at the canonical pyrenoid site, which we confirmed by expressing STA2-Venus as a starch marker (Mackinder et al., 2017) in *Crrbcs::AtRBCS* (**Fig. S6B**). We expressed BST4-Venus in *Crrbcs::AtRBCS* and found that BST4 localized to a punctum adjacent to the canonical pyrenoid site, which we attribute to the nascent pyrenoid tubules (**Fig. S6C**). To confirm this, we also expressed a known pyrenoid tubule marker protein, PsaF (Emrich-Mills et al., 2021) in *Crrbcs::AtRBCS*, which showed a similar localization pattern to that of BST4 (Fig. S6D). This suggests there may be an additional Rubisco-binding independent mechanism for localization of BST4 as well as other proteins to the pyrenoid tubules. Further investigation of the role of the whole C-terminal domain will be required to understand the mode of BST4 pyrenoid tubule localization.

### Chlamydomonas *bst4* mutant is not defective in growth at air level CO2 but increased H2O2 production

To test whether BST4 has a role in the operation of the CCM, we measured the growth of *bst4* compared the WT control strain under various CO2 conditions (**Figs. 6, S7**). Spot assays did not reveal any reduction in growth under CO2-limiting conditions (**Fig. S7**). When grown in liquid medium, *bst4* even seemed to grow slightly better than WT when sparged with 0.04% CO2 (**Fig. 6A**). However, when comparing the calculated specific growth rates (μ h^-1^) for both the exponential growth phase (days 0-3, *bst4* 0.0402 ± 0.0003 μ h^-1^ and WT 0.0389 ± 0.0007 μ h^-1^) or the full growth assay (days 0- 5, *bst4* 0.0257 ± 0.0001 μ h^-1^ and WT 0.0241 ± 0.001 μ h^-1^), there was no statistically significant increase in specific growth rates between *bst4* and WT (two-tailed t-test, p = 0.17 and 0.20, respectively, n=3, full results **Table S1**). We conclude from these experiments that BST4 is not essential for growth at air levels of CO2 and might not be necessary for the functioning of the CCM. We also included the complemented *bst4::BST4* and *bst4::BST4ΔC-term-mScarlet-I* (hereafter *bst4::BST4ΔC-term*) lines in the spot and liquid growth assays. While all lines grew well in the spot assay, in the liquid growth *bst4::BST4ΔC-term* grew comparably to WT and *bst4* whereas *bst4::BST4* exhibited a slightly reduced growth than the other lines in both CO2 conditions.

**Figure 6.**
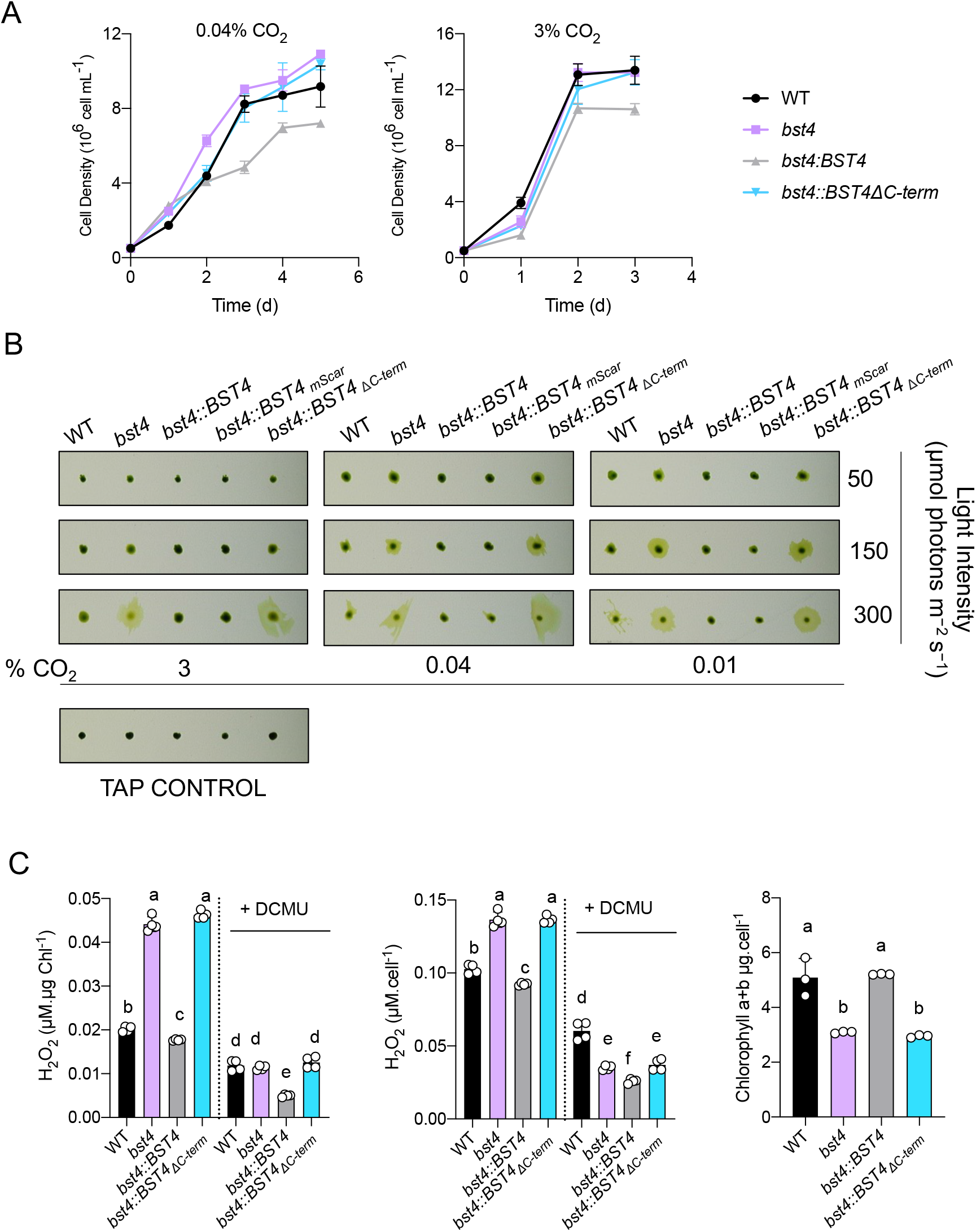
The *bst4* mutant does not have an impaired growth phenotype under CCM induced conditions but has increased H2O2 production. **A.***Chlamydomonas* strains were subjected to a liquid growth assay using pH 7.4 TP media that was bubbled with 0.04% CO2 or 3% CO2 (+/- 2 ppm). Error bars are ± SEM (n=3). **B.** Dot assay of WT, *bst4,* complemented *bst4::BST4* and *bst4* complemented with C-terminal truncation of BST4 (*bst4::BST4Δ*C-term) on minimal pH 7.4 TP agar at indicated light intensities and CO2 concentrations. **C.** H2O2 assay. Chlamydomonas cells were grown in pH 7.4 TP liquid media and exposed to 150 µmol photons m^-2^s^-1^ for 24 h with or without the photosynthetic inhibitor DCMU (10 µM). The concentration of H2O2 was subsequently quantified using Amplex Red (n=4), and is presented both proportionately to cell density and chlorophyll content. Chlorophyll content was quantified for all cell lines (n=3). Different letters indicate significance (p<0.05) as determined by a one-way Anova and Tukey’s post-hoc test.

One noticeable difference between *bst4* and WT lines from the growth assays conducted on solid media (**Fig. 6B and S7**) was that *bst4* cells had a distinct halo of diffuse cells on the periphery of the colony. We used a range of CO2 and light conditions to investigate the diffuse colony phenotype (**Fig. 6B**) and found it was most apparent under high light (300 μmol photons m^−2^ s^−1^) and low or very low CO2 conditions (0.04% CO2, 0.01% CO2, respectively) (**Fig. 6B**). WT, *bst4::BST4* and *b*s*t4::BST4-mScarlet-I* complemented lines had little or no diffusivity, although WT did display a slightly diffuse colony phenotype at 300 μmol photons m^−2^ s^−1^. Interestingly, *b*s*t4::*BST4ΔC-term was unable to rescue the diffuse colony phenotype suggesting that either the presence of the C-terminus or localization to the tubules is essential for the function of BST4.

The diffuse colony phenotype was most apparent under high light (300 μmol photons m^−2^ s^−1^), to a slightly lesser extent at medium light (150 μmol photons m^−2^ s^−1^), and exacerbated by low CO2 (0.01% and 0.04%). These are conditions where carbon fixation can be limiting and therefore, the energy production by photosynthesis could exceed the energetic demand required to fix CO2 by the CCM and the Calvin cycle. This imbalance can result in the release of Reactive Oxygen Species (**ROS**) (Erickson et al., 2015). Because cells exposed to ROS have altered phototactic responses to light (Wakabayashi et al., 2011), we thus thought to assess the phototactic capacity of *bst4* and its control WT strain by exposing them to directional light in liquid culture (**Fig. S8A**). In this assay, *bst4* cells displayed strong positive phototaxis, whereas WT and complemented *bst4::BST4* lines displayed negative phototaxis. To test whether the phototactic response of *bst4* was due to an increase in ROS production, we recorded the direction of phototaxis in cells exposed to either ROS or a ROS quencher. In the presence of the ROS quencher N,N’-dimethylthiourea (**DMTU**), *bst4* positive phototaxis was disrupted, resulting in a negative response to directional light (**Fig. S8A**). When the ROS H2O2 (75 μM) was added, WT, *bst4::BST4* and *bst4* displayed positive phototaxis (**Fig. S8A**). To directly quantify the difference in ROS generation, we analysed the H2O2 produced by cells exposed to 150 μmol photons m^−2^ s^−1^ (**Fig. 6C**). The *bst4* and *bst4::BST4ΔC-term* lines had significantly higher H2O2 production than WT and *bst4::BST4* when normalized to both chlorophyll content and cell density. We validated the assay by using the same ROS quencher DTMU from **Fig. S8A**, and saw a consistent reduction in H2O2 detected in all lines **Fig. S8B**. To test the involvement of photosynthetic activity in the increased ROS production, we also treated cells with the PS II plastoquinone binding site inhibitor 3-(3,4-dichlorophenyl)-1,1-dimethylurea (DCMU) whereby the concentration of H2O2 produced was reduced in all lines (**Fig. 6C**). The cellular chlorophyll content was also assessed, we found that *bst4* and *bst4::BST4ΔC-term* had significantly lower cellular chlorophyll content than WT and *bst4::BST4.* The reduction in cellular chlorophyll and the increase in ROS scavenging pigments is a known physiological response to high light exposure and subsequent elevated ROS production (Bonente et al., 2012) and is consistent with other studies (Ma et al 2020; Erikson et al 2015).

ROS generation occurs when light energy exceeds the capacity of electron transport, which can be caused by either high light conditions or limited CO2 fixation due to an impaired CCM or low CO2 availability (Santhanagopalan et al., 2021; Choi et al., 2022). As *bst4* can maintain similar growth to WT under low CO2 conditions, we conclude that the absence of BST4 might affect photosynthesis more broadly and not specifically influence the sustained delivery of CO2 to Rubisco.

### BST4 regulates the lumenal pH in Chlamydomonas during dark to light transition

To assess the impact of BST4 on photosynthesis, we used a pulsed amplitude modulation fluorimeter (**PAM**) to measure chlorophyll fluorescence upon a dark to light transition in all the strains (**Fig. 7**). We used the PAM to assess the quantum yield of PSII (**Y(II)**) and the amount of non-photochemical quenching of chlorophyll fluorescence (**NPQ**) (**Fig. S9**). NPQ is mediated by multiple mechanisms and harbors multiple components (Erickson et al., 2015), one of them, termed energy-dependent quenching, (**qE**) is quickly induced and relaxed and is mostly mediated by the activity of the proton-sensing Light Harvesting Complex Stress related 3 (**LCHSR3**) protein (Bonente et al., 2011; Peers et al., 2009; Steen et al., 2022; Tian et al., 2019). The magnitude of qE has recently been shown to be an indicator of the lumenal pH, the lower the pH, the higher the qE (Burlacot et al., 2022; Tian et al., 2019). While no differences in Y(II) were observed between the strains upon a dark to light transition (**Fig. S9C, D**), the NPQ of *bst4* mutant was transiently higher than WT (**Fig. 7A**), the NPQ becoming undistinguishable from WT after three minutes of illumination (**Fig. 7A**). The complementing *bst4::BST4* strain was undistinguishable from the WT control during the first three minutes of illumination and *bst4::BST4ΔC-term* had a NPQ similar to *bst4* (**Fig. 7A**). To establish the nature of the transiently increased NPQ in *bst4*, we used a shorter illumination time (**Fig. 7B**). The entirety of the 45% increase in NPQ for *bst4* mutant was quickly dissipated (**Fig. 7 B, C**) suggesting that it could be attributed to qE. Similar trends of NPQ kinetics were observed when cells were supplemented with HCO3^-^ before the measurement, although NPQ relaxation in the dark was different between lines (**Fig. S10**). Since *bst4* did not accumulate more LHCSR3 as compared to WT (**Fig. 7D, E**), we conclude that BST4 is involved in the lumenal pH regulation upon a dark to light transition.

**Figure 7.**
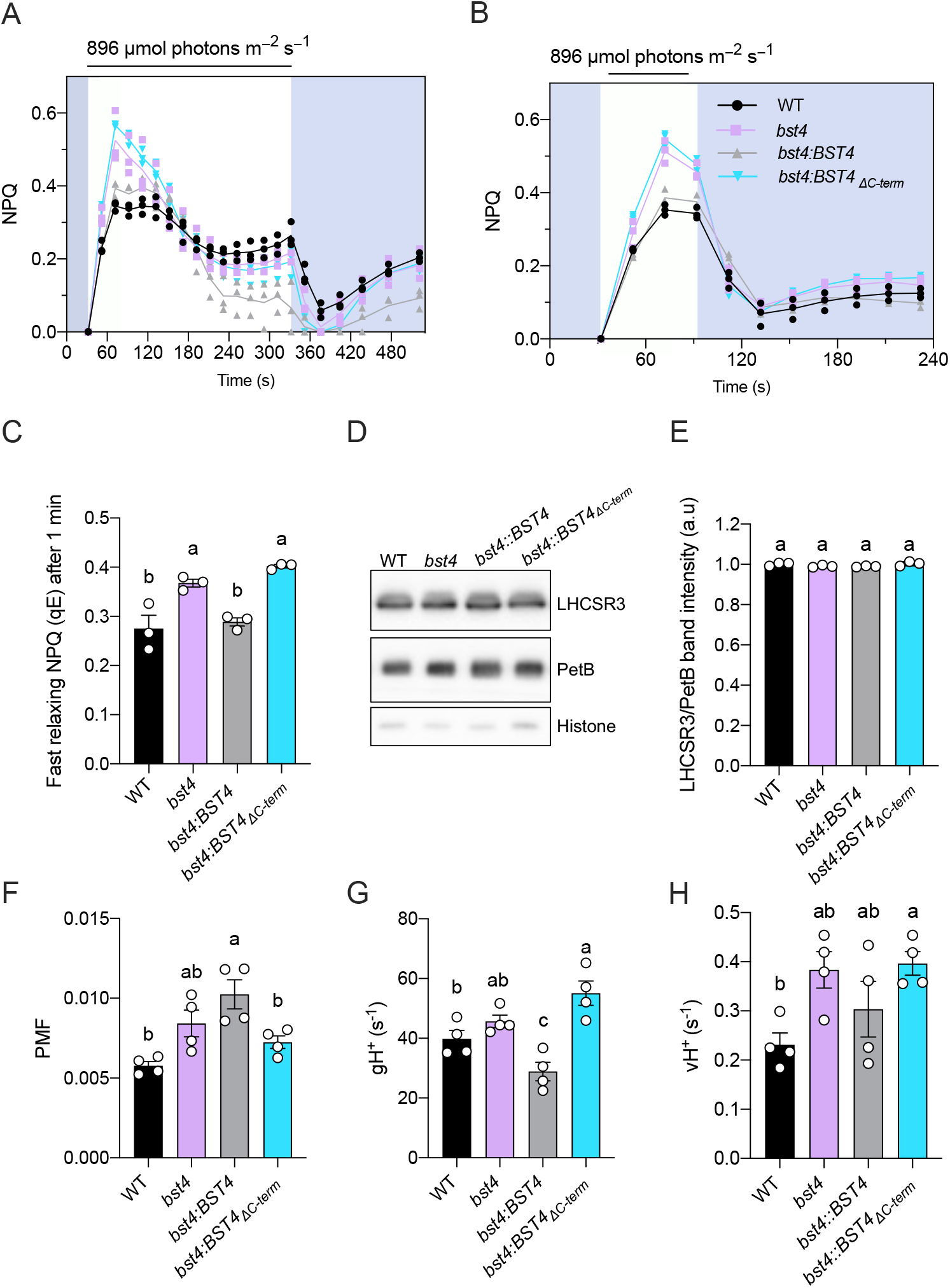
Chlamydomonas *bst4* mutant has an enhanced initial NPQ response under limiting Ci conditions. Wild type (WT) and mutants were grown in HS medium at 80 µmol photons m^−2^ s^−1^ and measured at 10 µg Chl ml^-1^ (prepared by dilution). The cells were dark adapted for 5 min before the measurements. **A.** Dynamics of Non-photochemical quenching (NPQ) on transition from dark to high light. Kinetics for induction of chlorophyll fluorescence were recorded during 5 min of illumination at 896 µmol photons m^−2^ s^−1^ followed by 5 min in darkness. **B.** Dynamics of NPQ during dark to light transition with one minute of light exposure. Shown are individual data points (dots) and their average (lines) **C.** Calculated fast relaxing NPQ after one minute of light exposure as determined by NPQ at the light to dark transition minus the minimum NPQ in the dark. **D.** Immunoblot of NPQ protein LHCSR3 in each strain plus PetB and Histone as loading controls.* **E.** Quantification of LHCSR3 band intensity normalized to PetB. Each point is the mean of two technical replicates from one biological replicate. **F.** Total proton motif force (PMF) as measured from ECS measurements. Shown are average of 3 technical replicates for each biological replicates (n=4 biological replicates) **G.** Proton conductance (g_H_^+^) and **H.** proton flux (v ^+^) were determined after 1 min illumination at 890 µmol photons m^−2^ s^−1^. (n=4 biological replicates). Bars show the mean ±SEM. Different letters indicate significance (p<0.05) as determined by a one-way ANOVA and Tukey’s post-hoc test.

The build-up of lumenal H^+^ concentration is usually accompanied with a build-up in the proton motive force (**PMF**) across the thylakoid membrane, which is used by the ATPase to generate ATP. We used Electrochromic Shift (**ECS**) measurements (Bailleul et al., 2010) to measure the total PMF size, as well as initial PMF dissipation rate (**gH^+^**) (**Fig. 7F, G & S11**). Interestingly, after one minute of illumination, which are conditions where the lumenal pH was higher in *bst4* mutant, neither the PMF, gH^+^ nor proton flux (**vH^+^**) differed between *bst4* and its WT strain. We conclude that the changes induced by BST4 on the lumenal pH are either a small contribution to the pmf formation or are compensated for in the *bst4* mutant.

To assess a longer timescale acclimation of the *bst4* mutant, we performed similar measurements but on cells pre-treated with three hours of high light (150 μmol photons m^−2^ s^−1^) and for a longer time-period (**Fig. S12**). We found that *bst4* was able to maintain a higher *F*v*/F*m than WT **(Fig. S12A),** had higher sustained NPQ compared to WT, which was concurrent with a higher LHCSR3 expression as compared to the WT strain **(Fig. S12B, G, H).** No significant differences in PMF or gH^+^ were observed although vH^+^ was higher in *bst4* compared to WT (**Fig. S12D, E, I, J**). The higher vH^+^ together with the higher expression of LHCSR3 suggest a build-up of protons in the tubule lumen. Since ROS induce photoprotective mechanisms (Roach & Na, 2017), we propose that in the absence of BST4, enhanced production of ROS **(Fig. 6)** might lead to enhanced NPQ and photoprotective mechanisms.

As a result of the NPQ difference seen between *bst4* and WT lines in response to illumination, we proposed that BST4 might be an anion channel involved in regulating the pH of the thylakoid lumen. Bestrophins are typically permeable to Cl^-^ and HCO3^-^. A plant thylakoid bestrophin, *At*VCCN1, is permeable to Cl^-^ and is also active in the first minutes of illumination to modulate the lumenal pH, although *vccn1* mutants have lower NPQ (Herdean et al., 2016). Alternatively, like BST1-3, BST4 may be permeable to HCO3^-^ (Mukherjee et al., 2019). To determine what BST4 may be permeable to, we expressed BST4 in Xenopus oocytes and measured currents in the presence of different anions (**Fig. S13A**). No currents for BST4 were detected in the presence of 100 mM KCl or 100 mM NaHCO3. As some bestrophins are autoinhibited by their C-terminus (Qu et al., 2006), we also tested two C-terminal truncations of BST4 (0-386 and 0-591), but no currents were detected. Another possibility is that BST4 is permeable to organic ions, similarly to *Hs*Best1, which has been shown to be permeable to **γ**-aminobutyric acid (GABA) (Lee et al., 2010) and glutamate (Woo et al., 2012), as well as Cl^-^ and HCO3^-^ anions. Fei et al. (2022) proposed a model whereby RuBP acts as proton carrier to increase H^+^ concentration in the pyrenoid tubules. It is possible that BST4 is the channel that facilitates RuBP translocation in the tubules in this model. To test this hypothesis, we used small molecule analogues K-PEP and K-Gluconate but no currents were detected for these either (**Fig. S13A**). Therefore, we were unable to draw conclusions as to what BST4 is permeable to. BST4 may require certain conditions to be open that are not met in the oocyte system, such as post-translational modifications, a specific pH, a specific voltage, or an interaction partner. BST4 was found to be phosphorylated and had an oxidized methionine residue in its C-terminus (Bergner et al., 2015). Methionine oxidation can serve as channel-regulating post translational modification (Ciorba et al., 1999), which would fit with the role that BST4 appears to have in preventing oxidative stress.

### BST4 has no impact on growth and photosynthesis in Arabidopsis S2Cr line

Although the permeability is undefined, if BST4 is a functional ion channel, it is possible that its presence in plant thylakoid membranes might have an effect. We generated three independent BST4 no tag lines in the S2Cr background and used them to assess the impact of BST4 on Arabidopsis physiology (**Fig. S14A**). We found there was no difference in growth between plant expressing BST4 and their azygous segregants, as determined by the rosette area (**Fig. S14B**). We also found that BST4 expressing plants tended to have slightly lower *Fv/Fm* as compared to azygous segregants and the parent line, although this was not significant (**Fig. S15A**). Further measurements were made for lines 2 and 3 on the kinetics of NPQ, Y(II) and for PMF size and partitioning but no consistent differences were observed (**Fig. S15**).

### Proposed model for the role of BST4 in the Chlamydomonas pyrenoid

Based on these findings, we propose that BST4 moderates the thylakoid lumen through its function as a bestrophin channel and is influential in this role during the transition from dark to light. Specifically, we propose that BST4 is a pentameric HCO3^-^ transmembrane channel found within the pyrenoid tubules but is not crucial for Rubisco matrix tethering. Rather, BST4 is targeted to the tubules by the C-terminal RBMs to facilitate its role in the pyrenoid. In its proposed primary role as a HCO3^-^ channel, BST4 may form part of a recovery system to take up HCO3^-^ from within the pyrenoid, allowing HCO3^-^ to be transported back into the tubules where it could be dehydrated to CO2 and thus available to be fixed by Rubisco **(Fig. 8).** The close proximity of BST4 to the luminal carbonic anhydrase CAH3 further supports this hypothesis. As indicated by our NPQ data, this role might be helpful during the dark to light transition where HCO3^-^ is needed immediately to both moderate lumenal H^+^ concentration and begin Ci uptake when Ci has not yet been transported into the thylakoid tubules. In this scenario, the absence of BST4 would lead to an initial build-up of protons as they are not being consumed by HCO3^-^ dehydration, thus the initial high NPQ. After a minute, the standard BST1-3 HCO3^-^ uptake system would take over, protons being used for HCO3^-^ dehydration and resulting in pH stabilization and NPQ relaxation. Over the longer term, a lack of BST4 may lead to a slight reduction in CCM operation efficiency, perhaps not sufficient to cause a growth penalty but enough to alter the homeostasis of photosynthesis and thus increase ROS production. Future work is needed to determine what species BST4 is permeable to and how it is regulated. In summary, BST4 might play a role in modulating the intricate dynamics between the CCM and photosynthetic ion management, two physiological processes that are key to algal primary productivity.

**Figure 8.**
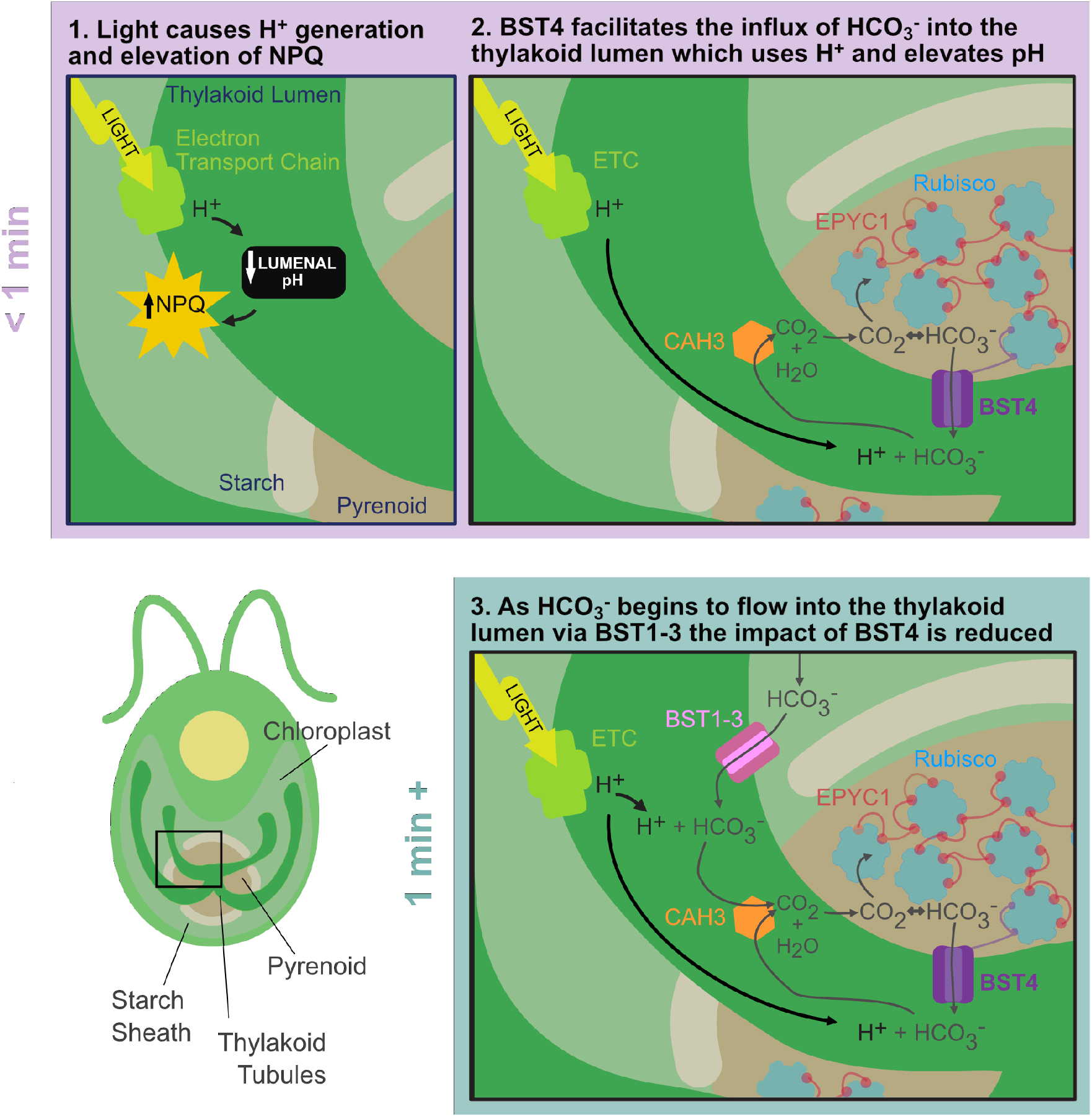
The role of BST4 in light and CO2 limiting conditions. The data presented here indicate that BST4 moderates the thylakoid lumen through its function as a bestrophin channel and is influential in this role during the transition from dark to light. 1) When the light is turned on, protons are generated by the electron transport chain (ETC) resulting in a luminal pH decrease and NPQ is initiation. 2) BST4 allows the passage of HCO3^-^ from the pyrenoid Rubisco-matrix into the lumen. The HCO3^-^ is then dehydrated by the carbonic anhydrase CAH3 to generate CO2, which can subsequently diffuse out of the lumen into the pyrenoid Rubisco-matrix for fixation. The conversion of HCO3^-^ to CO2 also acts as a sink for the protons generated by the ETC, increasing the luminal pH and reducing NPQ. This mechanism (shown in 1, 2) is most impactful in the first minute of illumination, after which **(3)** HCO3^-^ also begins to enter the thylakoid lumen through BST1-3 and the impact of BST4 uptake is reduced.

## MATERIALS AND METHODS

### Phylogenetic Analysis

Amino acid sequences for phylogenetic analysis were compiled by blasting BST4 (Cre06.g261750) in the NCBI database (Sayer et al 2022) and the manual addition of other well characterized bestrophin proteins including BEST1 *Homo sapiens* (XP_011543531.1), KpBEST *Klebsiella aerogenes* (WP_049046555.1), VCCN1 from *Arabidopsis thaliana* (Q9M2D2) and BST1-3 *Chlamydomonas reinhardtii* (Cre16.g662600, Cre16.g663400 and Cre16.g663450). Sequences were aligned using MAFFT (Katoh and Standley, 2013) visually inspected and manually trimmed (specified). Finalized alignments were run through IQTREE webserver (Minh et al., 2020) to identify the most appropriate substitution model. Maximum likelihood trees were then generated in Geneious v11 using the PhyML 3.0 (Guindon et al., 2010) plugin with a LG substitution model (Le and Gascuel, 2008) with Gamma distribution (4 categories) and 500 bootstrap iterations. Full alignments are found in **Fig. S16**.

### Alphafold structure prediction

Five chains of BST4 were submitted to Alphafold Multimer v2 with default settings. using the ColabFold server (Mirdita et al., 2022). All protein structure figures were generated using UCSF ChimeraX, developed by the Resource for Biocomputing, Visualization, and Informatics at the University of California, San Francisco, with support from National Institutes of Health R01-GM129325 and the Office of Cyber Infrastructure and Computational Biology, National Institute of Allergy and Infectious Diseases (Pettersen et al., 2021). The top ranked model of five was used for figure generation.

### Generation of plasmids

The plasmids for BST4 mutant complementation in *Chlamydomonas* were prepared using a recombineering method described previously (Emrich-Mills et al., 2021). BST4 is expressed under native promotor with either without a tag or with an mScarlet-I C-terminal tag, and a hygromycin AphVII selection marker. The same approach was used to generate STA2-Venus lines in the *Crrbcs::AtRBCS* background (Genkov et al., 2010).

Plasmids for plant expression were generated using the MoClo system (Engler et al., 2014). For visualisation, BST4 was combined with 35S promoter (pICH51277), mNeonGreen C-terminal tag (pICSL50015), HSP terminator and acceptor plasmid (pICH47732) and a pFAST-R section cassette used for selection. For all other experiments a no-tag BST4 construct was generated with the 35S protomer and HSP terminator parts and a Kanamycin resistance cassette was used for selection.

Plasmids for Xenopus expression were generated by using the Gateway system by cloning the coding sequence for BST4 into pGT vector (Grefen et al., 2010). *BST4* was amplified with Gateway adaptor sequences from a synthesized as g-block (IDT) with the PredAlgo (Tardif et al., 2012) predicted transit peptide removed from the N-terminal (sequence begins R35), any subsequent mutations were made via PCR.

### Arabidopsis transformation

Arabidopsis was transformed by floral dip as previously described in (Atkinson et al., 2016). BST4-mNeon primary transformants were screened for transgene insertion by seed fluorescence from pFAST-R and BST4 expression was confirmed by checking for mNeon fluorescence and by immunoblot. BST4 no tag primary transformants were screened using kanamycin resistance and immunoblot. Zygosity was checked via seed fluorescence from pFAST-R or kanamycin resistance.

### Chlamydomonas cell culture conditions and strain details

*Chlamydomonas reinhardtii* cultures were maintained as previously described (Ma et al., 2011). Tris-Acetate-Phosphate (TAP) and minimal (TP) media (acetate free) were prepared according to Sueoka (1960). TAP and TP agar plates for growth were made by adding 1.5% (w/v) agar. CMJ030 (CC-4533; *cw15, mt^-^)* and *bst4* (BST4 knock-out LMJ.RY0402.159478) were obtained from the CLiP collection at the *Chlamydomonas* culture collection (Zhang et al., 2014; Li et al., 2019). This CLiP mutant has two other mapped CIB1 cassette insertions at loci Cre04.g230046 and Cre08.g367750. The insertion of the CIB1 cassette in *BST4* locus was confirmed by PCR amplifying the insertion locus from genomic DNA (Fig. S3A) using loci specific primers (forward GAGCTTCGTGGATGGATGTT and reverse GTATGAAGGTCACCGCCTGT) in parallel with a control locus (forward ATGCTTCTCTGCATCCGTCT and reverse ATGTTTTACGTCCAGTCCGC). The additional two insertions were also confirmed using loci specific primers for Cre04.g230046 (forward TGTGCCTCTGTCAGTCTTGG and reverse TGCGTGGATGGGTAACAGTA), Cre08.g367750 (forward AATCAAGAAGCTTCCCAGCA and reverse CCTACCGCTATCTCAGCCAG) and STT7 locus as a control (forward GCACGAACCAAGACACACATAG and reverse GTAGACGATGTCACCGCACTT). Therefore, the *bst4* knock-out was complemented with *BST4* constructs described herein. All complemented lines were validated by western blotting of BST4 and specified epitope tags, described below (Fig. S3B-E).

### Chlamydomonas transformation

For each Chlamydomonas transformation, 28 ng kbp^−1^ of plasmid was linearized by restriction digest. Cells were grown to 2 - 4 x 10^6^ cell mL^-1^, harvested by centrifugation at 1000 xg for 10 min and resuspended in TAP with 40 mM sucrose at a concentration of 2 × 10^8^ cells mL^−1^. Linearized plasmid was mixed with 250 μL of cells at 15°C in a 0.4 cm gap electroporation cuvette and transformed immediately by electroporation using a Gene Pulser II (Bio-Rad) set to 800V and 25 μF. Cells were recovered overnight in TAP sucrose while shaking gently (140 rpm) in the dark. Transformed cell were subsequently subjected to selection by growth on TAP agar plates with paromomycin (20 μg mL^−1^) or hygromycin (25 μg mL^−1^) which were kept in low light (5–10 μmol photons m^−2^ s^−1^) until screening positive transformants.

### Chlamydomonas Growth Assays

Spot Tests: Cells were grown heterotrophically in TAP media. Once cultures reached 2 - 4 x 10^6^ cell mL^-1^, 1 x 10^6^ cells were harvested by centrifugation at 1000 xg for 10 min. Cell were washed and resuspended at a concentration of 1 x 10^6^ cell ml^-1^ in TP media. Liquid cultures were spotted onto TP agar (1.5%) in 1000, 100 and 10 cell spots at a range of pHs (specified). The plates were incubated in 3, 0.04 and 0.01% CO2 and illuminated under constant light at 400 µmol photons m^-2^ s^-1^. Growth was monitored for up to 10 days.

Liquid Growth: Cells were grown heterotrophically in TAP media. Once cultures reached 2-4 x 10^6^ cell mL^-1^, cells were harvested by centrifugation at 1000 xg and resuspended at a starting concentration of 1 x 10^5^ cell ml^-1^ in TP media pH 7.4. Cultures were incubated in a CellDEG HDC 22.10 culture platform (CellDeg GMBH, Berlin) bubbled with 0.04 and 3% CO2, illuminated at 150 µmol photons m^-2^ s^-1^ and consistently stirred at 180 rpm. Cell density and optical destiny (750 nm) measurements were taken daily for up to 10 days. Specific growth rates (SGR) per h were calculated using the following formula: *μ= ln(N2/N1)/t* whereby N = cell density Dot tests: Cultures were grown in a 96 format on agar plates and replicated by a Rotor+ (Singer Instruments) high throughput replication robot. The cultures were stamped onto pH 7.8 TP agar plates, incubated in 3, 0.04 and 0.01% CO2 and illuminated under constant light at a range of intensities (specified). Growth was monitored for up to 10 days.

### Phototaxis Assays

Chlamydomonas cells were grown heterotrophically in TAP media until they reached 2 - 4 x 10^6^ cell mL^-1^ and harvested by centrifugation at 1000 xg for 10 min. Pelleted cells were either re-suspended in TP media or, for ROS manipulation assays, a phototaxis buffer described previously by Ueki et al. (2016) (5 mM Hepes pH 7.4, 0.2 mM EGTA, 1 mM KCl, and 0.3 mM CaCl2). The assays took place in 12-well dishes with a thin layer of TP agar (0.8%) on the well bottom and approximately 1.5 x 10^7^ cells in 400 μL of homogenous suspension laid on top. The dishes were illuminated from one direction with 150 µmol photons m^-2^ s^-1^ illumination for up to 3 h. Plates were imaged using a Flatbed Scanner at specified intervals.

### Quantification of H2O2

Cells were grown heterotrophically in TAP media. Once cultures reached 2 - 4 x 10^6^ cell mL^-1^, cells were harvested by centrifugation at 1000 xg and resuspended at a concentration of 2 x 10^6^ cell ml-1 in TP media pH 7.4, illuminated at 150 µmol photons m^-2^ s^-1^ for 24 h and shaken at 140 rpm. For H2O2 quantification, 1 mL of culture was diluted at a 1:1 ratio with fresh TP media, containing 1U of horseradish peroxidase and 5 µM of Amplex Red (ThermoFisher) and incubated for 1 h (illuminated at 150 μmol photons m^-2^ s^-1^, shaking 140 rpm). Cells were removed by centrifugation. The H2O2 of the media was immediately quantified using a ClarioStar Plate Reader Excitation/Emission 520/570-600 and compared against a linear H2O2 standard curve up to 5 μM. Additional controls were included; some cells were treated with the ROS quencher N,N’ dimethylthiourea (DMTU) at a final concentration of 150 µM; and some with the PSII plastoquinone binding site blocker 3-(3,4-dichlorophenyl)-1,1- dimethylurea (DCMU) was dissolved in methanol at a final concertation of 10 μM prior to H2O2 quantification (specified). All measurements were conducted with a minimum of four technical replicates. The data shown represents one of a multiple of experimental repeats conducted on different days with fresh cultures. All H2O2 concentrations were normalized to cell density, calculated as described previously, and chlorophyll content, described below.

Total chlorophyll was calculated by resuspending 1 mL of harvested cells in 1 mL of methanol. All samples were protected from the light after menthol addition. After vortexing for 1 min to resuspend the pellet and incubating for 10 min, the cells were removed by centrifugation. The absorbance of the supernatant was analysed by spectrophotometer at 652 and 665 nm. Total chlorophyll was calculated using the formula below. All measurements are averaged from three technical replicates. Total chlorophyll (µg/mL) = 22.12 x Abs652 + 2.71 x Abs665.

### Growth of Arabidopsis

Arabidopsis seeds were sown on moist F2+S soil and stratified in the dark at 4℃ for 2 days. For growth experiments seeds were grown in a Percival SE-41AR3cLED chamber (CLF PlantClimatics GmbH, Wertingen, Germany) equipped with cool white LED lights under 12 h light (175-180 µmol photons m^-2^s^-1^)/12 h dark cycles at 21°C, respectively and 70% relative humidity.

### Chlamydomonas Confocal Microscopy

Transgenic fluorescent strains were initially grown heterotrophically in TAP media until reaching 2 – 4 ×10^6^ cells mL^-1^ and resuspended in TP media overnight prior to imaging. Cells were mounted on 8-well chamber slides and overlayed with 1.5% low melting point agarose made with TP-medium. Images were collected on a LSM880 (Zeiss) equipped with an Airyscan module using a 63× objective. Laser excitation and Emission setting for each channel used are set as below: Venus (Excitation: 514 nm; Emission 525 – 500 nm); mScarlet-I (Excitation: 561 nm; Emission 570 – 620 nm);

Chlorophyll (Excitation: 633 nm; Emission 670 – 700 nm).

### Yeast-2-Hybrid

Yeast two-hybrid to detect interactions between BST4 C-terminus and RbcS1 was carried out as described in He et al., (2020). BST4 C-terminus (amino acids 387-end) was cloned into the two-hybrid vector pGBKT7 to create a fusion with the GAL4 DNA-binding domain. Point mutations were introduced by PCR into BST4 RBMs, which was then cloned into the same vector. Mature CrRBCS1 was cloned into the vector pGADT7 to create a fusion with the GAL4 activation domain. Yeast cells were then co-transformed with binding and activation domain vectors. Successful transformants were cultured, diluted to an optical density at 600 nm (OD600) of 0.5 or 0.1 and plated onto SD-L-W (double drop out, DDO) and SD-L-W-H (triple drop out, TDO) media. The plates were imaged after three days and **Fig. S2** shows yeast spots from cultures diluted to an OD600 of 0.5.

### Slimfield microscopy

Chlamydomonas lines *bst4::BST4*-*mScarlet-I* and the unlabelled control line *bst4* were prepared overnight in TP media. Each was harvested and spotted onto a slide-mounted agar pad (GeneFrames, ThermoFisher), consisting of TP media with 1.5% low melting point agarose. Fluorescence imaging with single-molecule sensitivity was performed using a custom Slimfield microscope (Syeda et al., 2019). The setup used a high-magnification objective (NA 1.49 Apo TIRF 100× oil immersion, Nikon) and the detector was a Prime95B sCMOS camera (Teledyne Photometrics) operating in 12- bit ‘sensitivity’ gain at a high total magnification of 53 nm/pixel. The samples were illuminated either in brightfield, or for Slimfield fluorescence in camera-triggered frames by a collimated 561 nm wavelength, Gaussian mode OPSL laser (Coherent, Obis LS) at a peak intensity of 5 kW/cm^2^ at the sample plane. This beam was tilted into a HILO configuration (Payne-Dwyer and Leake, 2022) to reduce out-of-focus excitation while retaining quantitative molecular sensitivity. The fluorescence image was split into two parallel channels comprising emission bandpass filters (Semrock BrightLine^®^): one with a 585/15 emission filter (central wavelength/spectral bandwidth in nm) optimized to isolate the mScarlet-I signal, and a second with a 525/25 emission filter, used only to indicate autofluorescence background. The total length of each acquisition sequence was ∼5 s; sufficient to observe the full course of mScarlet-I photobleaching, from the initial unbleached state to single-molecule blinking, while also rapid enough (10 ms exposure/frame at 180 frames/s) to capture the motion of individual molecular assemblies.

### Single particle tracking and molecular counting

Slimfield image sequences were segmented manually in ImageJ to include only the pyrenoids in downstream analysis. The centroid positions of fluorescent tracks were identified from local intensity maxima in each frame using ADEMScode software in MATLAB (Wollman et al., 2022). The summed intensity of each candidate track was calculated in each frame by adding all pixel values within 5 pixels of the centroid, then subtracting the local background averaged between 5 - 8 pixels from the centroid. Candidates with a summed intensity <0.4× the standard deviation in the background region were discarded.

Fluorescent proteins are known to exhibit a characteristic integrated intensity per molecule under stable Slimfield imaging conditions and within the quasi-uniformly illuminated area within half the beam waist (Shepherd et al., 2021). After sufficient photobleaching of mScarlet-I in the Slimfield image sequences, only step-like blinking was observed at the end of each track. The modal integrated intensity of these steps was used to estimate this characteristic single molecule brightness, equivalent to 56 ± 9 photoelectrons per frame per molecule.

At the start of each track, we obtained an initial integrated intensity (independent of photobleaching) by linearly extrapolating the summed intensity backwards over the first 4 frames of the exposure. This initial intensity was then divided by the characteristic brightness of a single mScarlet-I to estimate the number of molecules, or stoichiometry, in that track. This estimate was precise enough to detect stoichiometry steps of up to 12 tagged molecules without ambiguity.

Stoichiometry distributions may exhibit peaks which are separated by a characteristic interval. The smallest consistent interval between peaks can be used to infer the size of a physical repeat unit or ‘periodicity’ within assemblies (Hunter et al., 2022; Payne-Dwyer et al., 2022). A kernel width of 0.7 molecules was chosen to generate the stoichiometry distribution (Fig. 2F), reflecting the background standard deviation. Peaks were pinpointed using MATLAB’s *findpeaks*.

The intervals between all peaks for each acquisition were aggregated across the pyrenoid population, weighted by inverse square-root distance (thereby accounting for shot-noise in broader intervals). A second distribution **(Fig. 2G)** was then generated from this weighted population of intervals. The kernel width in this estimate was 0.7 molecules multiplied by the square root of the mean stoichiometry divided by the root number of intervals (thereby accounting for shot-noise in intervals between peaks of higher stoichiometry). The periodicity was then reported as the mode of this distribution and its 95% confidence interval.

### Arabidopsis Confocal microscopy

Small sections of 3 – 4 week-old leaf tissue (∼5 - 10 mm^2^) were adhered to slides using double-sided tape with basal side up. A x40 water immersion objective lens was used. Samples were excited by 488 nm at 1% laser power, chlorophyll autofluorescence was collected at 680 – 750 nm and mNeonGreen fluorescence at 503–532 nm. For dual tagged lines, we used sequential acquisition to minimize bleed-through. mCherry was excited using the 542 nm laser and emission collected at 601-620 nm and mNeon as before. Images were acquired using the SP8 Confocal system and Leica LAS AF software (http://www.leica-microsystems.com/). Figures were prepared using ImageJ (http://fiji.sc/Fiji).

### Immunoblot detection

Two leaf disks (6 mm diameter) were harvested and immediately frozen in liquid nitrogen. Two steel balls (3 mm) were added and tissue was homogenized using a tissue-lyser twice for 30 Hz for 30 s. A four times volume of cold extraction buffer (20 mM Tris-HCl pH = 7.5, 5 mM MgCl2, 300 mM NaCl, 5 mM DTT, 1% (v/v) Triton X-100, 1 x Protease inhibitor (Roche)) was added and samples vortexed for 30 s. Samples were solubilized on ice for 5 mins and then centrifuged at 5000 xg for 5 min at 4℃. 17.5 µL of supernatant was used to make up 1x LDS and 100 uM DTT. 20 µL was loaded on a Novex™ 4 - 12% Bis-Tris Mini Gel, (Thermo Fisher, Catalogue number: NP0322BOX). The gel was run at 150 V for 60 minutes. Proteins were transferred to a nitrocellulose membrane using an iBlot 2, programme 0. The membrane was probed with primary antibody in 5% milk in 1 x TBST at the following dilutions: BST4 (1:1000; generated for this study, peptide from C-terminus: SDTELSEANRPRTRPDWRN) (YenZym, Antibodies LLC, USA), AtpB (1:2000; Agrisera:AS05085), RbcL (1:1000; kind gift from Griffiths lab), CP43 (1:3000; Agrisera: AS111787), PsaB (1:1000; Agrisera: AS10695), PsbO (1:2000; Agrisera:AS06142-33). Secondary antibody (goat ɑ-rabbit IR-800; Li-COR: 925-32211). Membrane was imaged using the Li-COR Odyssey CLx scanner.

In order to quantify BST4 protein in Chlamydomonas lines, cells were grown in TP media at ambient CO2 until reaching 2 - 4 x 10^6^ cells mL^-1^. Cells were harvested by centrifugation at 1000 x g for 10 mins, normalized to Chl content and resuspended in the extraction buffer described above. Samples were freeze-thawed three times and spun at 20, 000 x g for 20 min at 4℃. Protein extractions containing 5 µg of Chl with 1 x SDS loading buffer were boiled at 100℃ for 5 min and loaded onto a 4 - 20% polyacrylamide gel (Mini Protean TGX, Biorad Laboratories). Proteins were transferred to a PVDF-FL membrane on a Biorad semidry blotting system. BST4 primary antibody was used as described above alongside alpha-tubulin primary antibody raised in mouse (Agrisera), as a loading control. Anti-rabbit and anti-mouse fluorescent secondary antibodies, Invitrogen AlexaFluor 488 and 555 respectively, were used at a 1:20 000 dilution. Immunoblots were imaged using an Amersham typhoon 5 scanner with 488 and 535 excitation lasers and Cy2 and Cy3 emission filters. BST4 band fluorescent intensity was quantified using FIJI (Image J) (Schindelin et al., 2012) and normalized to alpha-tubulin loading control. All Chlamydomonas lines for quantification were extracted and analysed in triplicate. For LHCSR3 protein quantification, cells were seeded at 0.1 OD750 in TAP media for 4 days and then switched to TP media at a concentration of 30 µg Chl ml^−1^ and exposed to 150 µmol photons m^−2^ s^−1^ for 3 h. Protein was extracted according to Burlacot et al. 2022 and separated by SDS-PAGE as described for BST4 immunoblots.

Oocytes for expression of BST4 were collected after recording and prepared for western blot as described in Lefoulon et al., 2018. The BST4 primary antibody was used as described above and the secondary antibody used was horseradish peroxidase-coupled goat, anti-rabbit (dilution 1:10000 Abcam). Proteins were detected with ECL Advance kit (GE Healthcare,Poole UK).

### Blue-Native PAGE

A crude thylakoid enrichment was performed according to (Aro et al., 2004). Thylakoid membranes were solubilized in 0.5% DDM, 1x NativePAGE buffer (Thermo: BN2003), 1x cOmplete protease inhibitor tab (Roche;10x stock made by dissolving 1 tablet in 1 mL dH2O) for at a concentration of 0.8 µg Chl/µl for 15 min on ice. Unsolubilized material was removed by two rounds of centrifugation at 17,000 xg at 4°C for 15 min. 19.5 µL of supernatant was combined with 0.5 µL of Coomassie additive and loaded immediately onto a 4-16% Bis-tris gel (Thermo: BN1002BOX). Electrophoresis was performed at room temperature at 150 V for 90 min. Cathode buffer was swapped from dark to light when the dye front was a third way through the gel. Separated proteins were transferred to a nitrocellulose membrane by electrophoresis at 100 V for 90 min at 4 ℃.

Proteins were visualized using chemiluminescence. Secondary antibody (goat ɑ- rabbit, HRP;1:10,000; Abcam: ab6721). Chemiluminescence substrate SuperSignal West Pico PLUS (ThermoScientific, ref number: 34579) according to manufacturer’s instructions. Chemiluminescence was detected using clear blue X-Ray Film CL- Xposure^TM^ Film (ThermoScientific, ref number: 34090).

### Chlorophyll fluorescence measurements in Chlamydomonas for Fig. 7

To measure PSII activity in Chlamydomonas, cells were grown in HS media under 80 µmol photons m^−2^ s^−1^ for 3 days at 120 rpm (in a Multitron, Infors-ht) to reach logarithmic phase and then maintained at ∼10 µg Chl mL^-1^. Two millilitres of cells were added to a cuvette and bubbled with air (10 cc/min) and continuous stirring. Cells were incubated in the dark for five minutes before recording Chl fluorescence using DUAL- PAM-100 (Walz, Effeltrich, Germany). A saturating pulse of 8,000 µmol photons m^−2^ s^−1^ of 300 ms was applied to the samples for determination of the maximal fluorescence yield in the dark state (*Fm*) and maximal fluorescence yield during the period with actinic light (*Fm’*). The maximal quantum efficiency of PSII was calculated as (*Fm^’^-F)/Fm* where F is the stationary fluorescence. NPQ was calculated as (*Fm– Fm’*)/*Fm*’. Far-red light (4 µmol photons m^−2^ s^−1^) was used throughout the entire experiment to limit state transition from contributing to the NPQ. NPQ and Y(II) were calculated based on changes in Chl fluorescence as (*Fm–Fm’*)/*Fm*’ and (*F*m’–*F*)/ *Fm*’, respectively, according to Genty et al., (1989). When indicated, cells were supplemented with a final concentration of 500 µM HCO3^-^ at the beginning of the dark adaptation.

### Electrochromic shift (ECS) in Chlamydomonas for Fig. 7

Electrochromic shift (ECS) in Chlamydomonas was assessed by measuring the absorbance changes of cells at 520 and 545 nm using a JTS-100 spectrophotometer (BioLogic). Cells were grown and prepared as for Chl fluorescence experiments described previously except cells were resuspended to 150 µg Chl mL^-1^ before being loaded into a custom vertical light path cuvette. Cells were dark adapted for 1 min and then exposed to 890 μmol photons m^−2^ s^−1^ red light (630 nm) for 1 min. The light was switched off and decay kinetics were measured. ECS signal was calculated as the difference between absorbance changes measured at 520 and 545 nm. For each biological replicate, 3 technical replicates were taken and averaged. PMF size was calculated as the difference between the ECS signal in light and the minimum value of the ECS signal immediately after the light was turned off. The gH^+^ parameter was calculated as 1/τ (time constant for decay during the first 100 ms (Cruz et al., 2005).

### Chlorophyll fluorescence measurements in Chlamydomonas for Fig. S12

To measure PSII activity in Chlamydomonas, cells were grown in TAP in low light (20 µmol photons m^−2^ s^−1^) for 4 days at 50 rpm to reach logarithmic phase. Cells were then washed, resuspended to 30 µg Chl mL^-1^ in TP media and exposed to 150 µmol photons m^−2^ s^−1^ light for 3 h, followed by 1 h incubation in darkness at 50 rpm before recording Chl fluorescence using DUAL-PAM-100 (Walz, Effeltrich, Germany). A saturating pulse of 3,000 µmol photons m^−2^ s^−1^ of 800 ms was applied to the samples in a cuvette under continuous stirring for determination of the maximal fluorescence yield in the dark state (*Fm*) and maximal fluorescence yield during the period with actinic light (*Fm’*). The maximal photochemical efficiency of PSII (*Fv/Fm*) was calculated. NPQ was determined from slow kinetics during actinic illumination at 1,500 μmol m^−2^ s^−1^ for 17 min followed by 5 min of dark relaxation. NPQ and Y(II) were calculated based on changes in Chl fluorescence as (*Fm–Fm’*)/*Fm*’ and (*F*m’–*F*)/ *Fm*’, respectively, according to Genty et al., (1989).

### Electrochromic shift (ECS) in Chlamydomonas for Fig. S12

ECS measurements in Chlamydomonas were carried out using a the Dual-PAM-100 equipped with a P515/535 module (Walz). Cells grown and prepared as for Chl fluorescence experiments described previously were layered on a glass slide and exposed to actinic red light for the given period. The light was switched off and decay kinetics were measured. PMF size was calculated as the difference between the ECS signal in light and the minimum value of the ECS signal immediately after the light was turned off. Calculation of ΔpH and ΔΨ was performed using the steady-state time point of the ECS signal in darkness (Cruz et al., 2001). Before each ECS measurement, a 3 saturating 50-μs actinic red flashes of 200,000 μmol photons m^−2^ s^−1^ was applied to determine the ECSST; subsequently, the ECSST amplitude was used to normalize the ECS signal before the calculation of PMF size and partitioning values. To determine H^+^ conductivity (gH^+^), the light was switched off at specific time points to record the ECS signal decay during 620 ms dark intervals. The gH^+^ parameter was calculated as 1/τ (time constant for decay during the first 100 ms (Cruz et al., 2005). The total proton flux across the membrane was calculated as νH^+^ = PMF *× g*H^+^ (Cruz et al, 2001).

### Chlorophyll fluorescence measurements for Arabidopsis

Plants were grown for 8 weeks on S-Hasselfors soil in a Percival AR-82L chamber (CLF Plant Climatics, Wertingen, Germany) using 12 h light (180 µmol photons m^−2^ s^−1^)/ 12 h dark cycles at 21°C/19°C, respectively, and 70% relative humidity. Slow kinetics of Chl *a* fluorescence induction were recorded with a pulse-amplitude modulated fluorometer DUAL-PAM 100 equipped with DUAL-DB and DUAL-E emitter-detector module (Walz) on attached leaves of 30 min dark-adapted plants using actinic red light of 830 μmol photons m^−2^ s^−1^ for 10 min, followed by a 5 min dark period. The saturating pulse applied was 5,000 µmol photons m^−2^ s^−1^ and of 800 ms duration. NPQ and (Y(II)) were calculated based on changes in Chl fluorescence as (Fm–Fm’)/Fm’ and (Fm’–F)/ Fm’, respectively (Genty et al., 1989).

### Electrochromic shift (ECS) for Arabidopsis

ECS was recorded with a DUAL-PAM 100 system equipped with a P515/535 emitter/detector module (Walz). First, plants were dark adapted for 30 min, then illuminated with actinic red light at 830 µmol photons m^−2^ s^−1^ for 3 min followed by a 60 s dark period in which the ECS decay kinetics were recorded. Before each measurement, three pulses of 5 µs and 200,000 µmol photons m^-2^ s^-1^ were applied to determine ECSST, which was used to normalize the ECST values of each measurement.

### Sedimentation of proto-pyrenoid

Cross-linked samples were prepared by vacuum infiltrating intact leaves 1% formaldehyde prior to sedimentation 200 mg of leaf tissue was flash frozen in liquid nitrogen and ground by bead beating twice at 30 Hz for 30 s. Four times volume of extraction buffer (50 mM HEPES-KOH pH 7.5, 17.4% (v/v) glycerol, 2% (v/v) Triton X- 100, cOmplete protease inhibitor tab) was added and sample mixed by bead beating again. Extract was filtered through one layer of miracloth. A small aliquot of filtered extract was saved as the input. Extract was then centrifuged at 500 x g for 3 min at 4℃ and the pellet discarded. The supernatant was centrifuged at 500 x g for 12min, 4℃. The pellet was washed once with extraction buffer and then centrifuged again. The pellet was resuspended in 100 µl extraction buffer then centrifuged for a further 5 min. Pellet was finally resuspended in 25 µl extraction buffer. Fractions were made up in 1x LDS loading buffer and 200 mM DTT. Ten microlitres was subjected to SDS-PAGE (NuPAGE™ 4-12% Bis-Tris Mini Gel, Thermo Fisher, Catalogue number: NP0322BOX) at 150 V for 60 min.

### Chloroplast fractionation

In order to biochemically localize transgenically expressed BST4, chloroplasts were fractionated as described in Herdean et al. (2016) using 100 g of leaf tissue from four-five week-old BST4 transgenic plants. The stromal fraction was concentrated using a 10,000 MWCO centrifugal concentrator (Sartorius Stedim Biotech GmbH, product number: VS1502).

### Electron microscopy of Chlamydomonas

High (3%) and low (0.04%) CO2 acclimated cells were harvested by centrifugation (×1000g, 4 minutes, 20°C). Primary fixation was performed in 1.25% glutaraldehyde in 50 mM sodium cacodylate (pH 7.15) in TP medium for 15 minutes followed by 2.5% Glutaraldehyde in 50 mM cacodylate for 2 hours. Fixed samples were washed three times with 50 mM sodium cacodylate by centrifugation. Samples were then osmicated with 1% OsO4 in 50 mM sodium 25 cacodylate for 1 hour on ice and washed with de-ionised water. Samples were block stained in 1% uranyl acetate in the dark for 1 hour. Samples were washed twice with dH2O and twice with 50 mM sodium cacodylate. Fixed samples were dehydrated in an acetone series (25%, 50%, 75%, 90% and 100%) ∼20 minutes each step. Dehydrated samples were infiltrated with Spurr’s resin by incubating in 25% then 50% Spurr resin in acetone for 30 minutes and transferred to 75% for 45 minutes at room temperature. They were then incubated in 100% Spurr resin overnight before polymerising at 70 °C for 24 hours. Sections ∼70 nm thick were collected on Copper grids and stained with saturated uranyl acetate and lead citrate. Images were collected with a FEI Tecnai 12 BT at 120kV using a Ceta camera.

### Electron microscopy of Arabidopsis

Leaves were cut into small 5 mm strips and fixed in 4% (v/v) Paraformaldehyde/0.5% glutaraldehyde in 0.05M sodium cacodylate (pH = 7.2) by vacuum infiltration three times for 15 min and incubation at 4 °C overnight with gentle agitation followed by dehydration in increasing amounts of ethanol 50/70/80/90/100% 1 hr each then overnight rotation. 100% ethanol was repeated 3 times, 1 hr each and a final over night at 4 °C. Samples were then fixed in LR resin by infiltrating with increasing concentration (50/70/100%) with a repeat of 100% and then polymerized overnight at 50 °C. Ultrathin sections were cut and mounted onto plastic-coated copper grids. Grids were stained with 2% uranyl acetate and visualized by transmission electron microscopy (TEM).

### Protease protection assay

Investigation of the orientation of BST4 in isolated thylakoid membranes was conducted as described in Stepien and Johnson (2018). Briefly, trypsin made up in 50 mM acetic acid according to the manufacturer’s instructions (Thermo Scientific, ref number: 90057). To disrupt the thylakoid membrane and allow degradation to lumen-facing peptides, 1% (v/v) Triton X-100 was added, and tubes gently agitated prior to addition of trypsin.

### Xenopus oocyte electrophysiology

Destination clones containing BST4 for Xenopus expression were linearized with EcoRI, before proceeding to the *in vitro* transcription (mMessage mMachine T7 transcription kit, Thermofisher Scientific, ref number AM1344). Stage IV oocytes were injected with 20 ng of RNA per oocyte. Measurements of ion transport was done by voltage clamp using an Axoclamp 2B amplifier (Axon instruments, Foster City, CA) (Lefoulon et al., 2014; Leyman et al., 1999). They were performed under perfusion of either 100 mM KCl, K-HCO3, Na-HCO3, K-PEP, or K-Gluconate, with 1 mM CaCl2, 1.5 mM MgCl2, and 10 mM HEPES-NaOH, pH 7.3. Recordings were obtained and analysed using Henry IV software (Y-Science, Glasgow, UK). BST4 expression in oocytes was validated by western blotting (Fig S10B), as described above.

## ACKNOWLEDGEMENTS

The authors thank the University of York Horticulture team who helped with setting up the Chlamydomonas phototaxis assays. TEM for Arabidopsis was carried out with the support of Stephen Mitchell and the Wellcome Trust Multi User Equipment Grant (WT104915MA) For the purpose of open access, a Creative Commons Attribution (CC BY) licence is applied to any Author Accepted Manuscript version arising from this submission.

## AUTHOR CONTRIBUTIONS

LA, CW, CSL, AB, CS, AM and LM conceived and designed the study with input from all authors. LA performed all Arabidopsis experiments, BN-PAGE and immunoblots. CW performed Chlamydomonas physiology experiments and immunoblots. CSL performed Chlamydomonas microscopy imaging and immunoblots. LA, AB, KMS, KAvM, and ED performed Chlamydomonas and Arabidopsis Chl fluorescence and ECS experiments and generated figures. APD and CW performed molecular tracking experiments with input from ML. CL and MB performed xenopus oocyte work. PG and JB did *in silico* structure predictions. TEM contributed to the complementation of Chlamydomonas *bst4* mutant. NA contributed to Arabidopsis transgenic line generation. GP assisted with experimental design and data interpretation. LA and CW generated figures. LA led the writing of the manuscript with input from CW. All other authors provided manuscript feedback.

## Supplemental Figures

**Supplemental Table 1.**
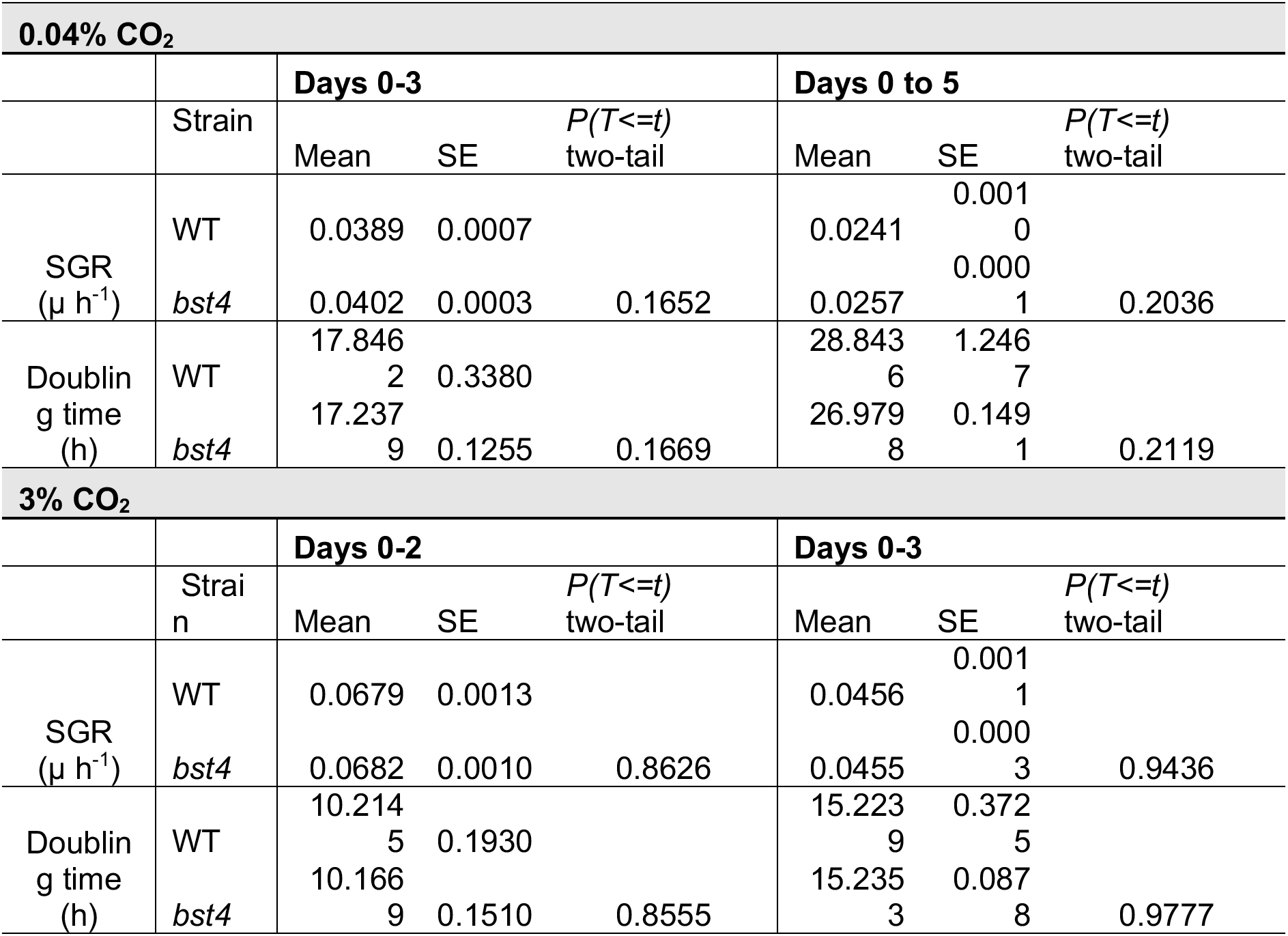
compares specific growth rates (μ h^-1^) and cell doubling times of WT and *bst4* strains (n=3) during liquid growth assays at 0.04 and 3 % CO2 (+/- 2 ppm).

**Supplemental Figure 1.**
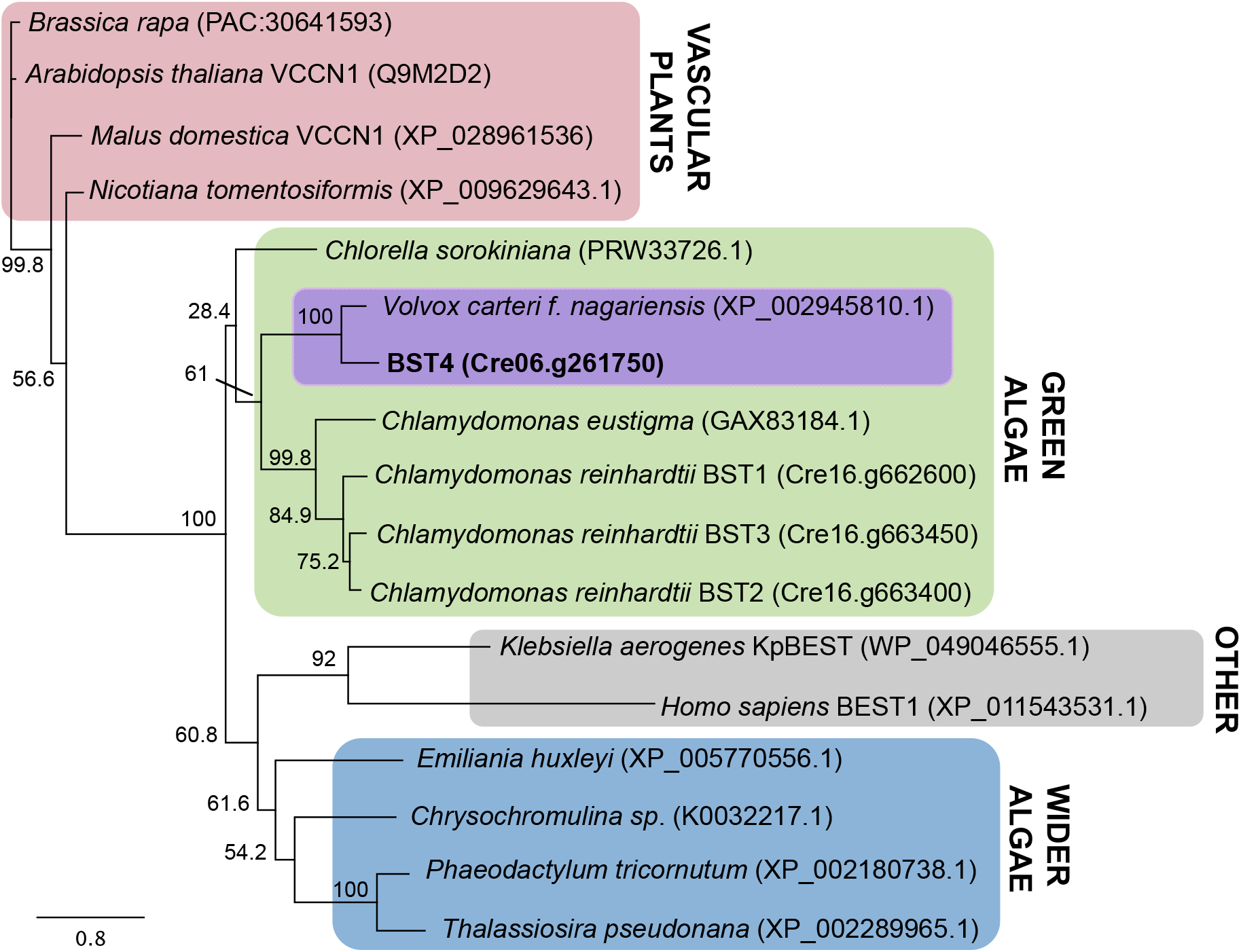
BST4 structure and sequence compared to other bestrophins. Phylogenetic analysis of the full length Chlamydomonas BST4 amino acid sequence (bold). The evolutionary history of BST4 was inferred by using the maximum likelihood method based on the Le and Gascuel substitution model with discrete Gamma distribution (5 categories) and 500 bootstrap replicates. The tree is drawn to scale, with branch lengths measured in the number of substitutions per site.

**Supplemental Figure 2.**
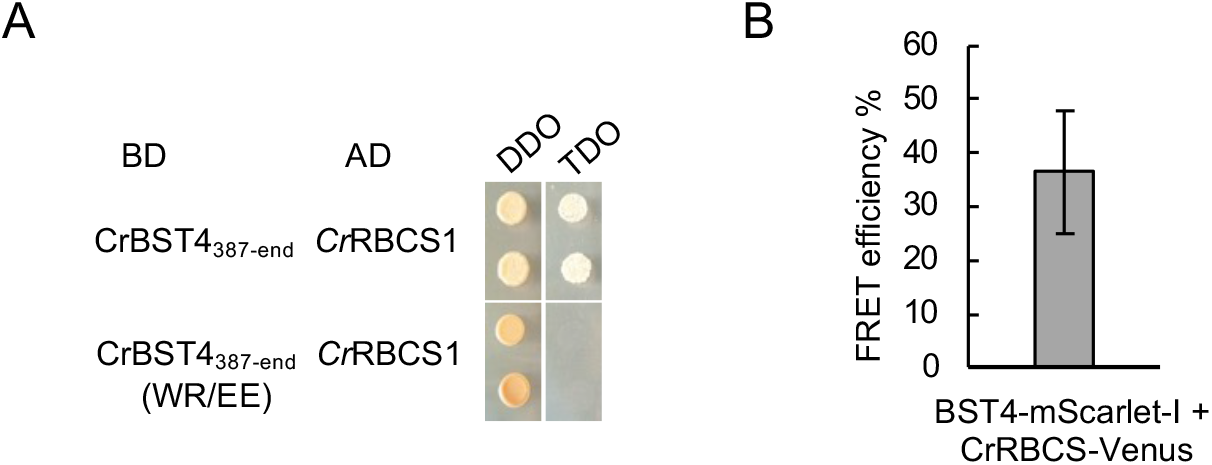
Interaction between BST4 and RBCS. **A.**Yeast-2-hybrid experiment. The disordered region of BST4 C-terminus (amino acid 387-end) and with residues WR from RBMs are changed to EE. Abbreviations: AD, activation domain; BD, binding domain; DDO, double drop out media; TDO, triple drop out media. Growth on TDO indicates an interaction. **B**. FRET efficiency between BST4-mScarlet-I and CrRBCS-Venus. Sensitized-FRET measurement on dual-tagged cells (BST4-Venus and RBCS1-mCherry) showed a moderate FRET efficiency of 35% (n=10).

**Supplemental Figure 3.**
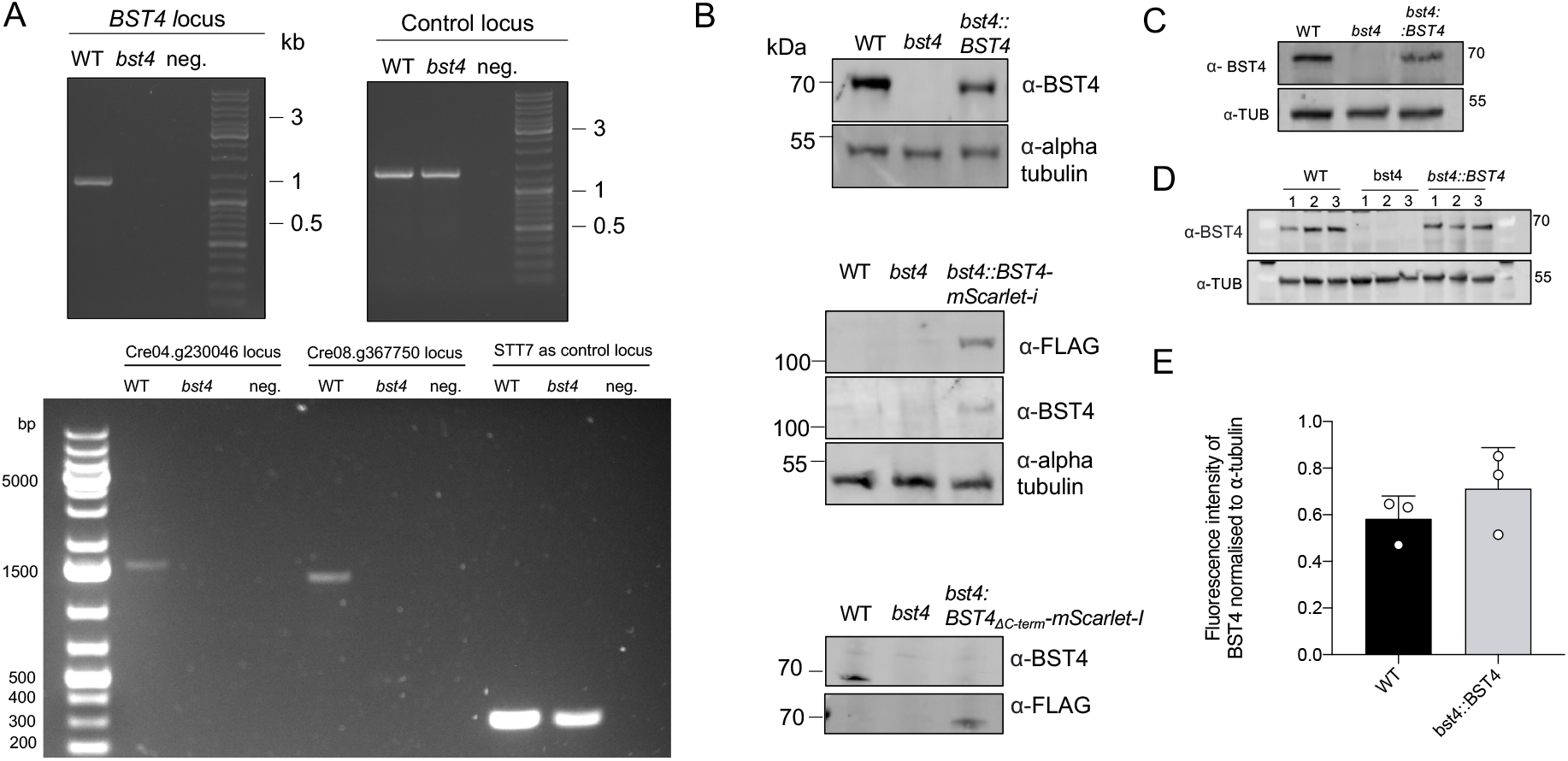
Validation of Chlamydomonas BST4 lines. **A.**PCR amplification of the mapped CIB1 insertion sites and control loci of *bst4* (LMJ.RY0402.159478) and WT control strain (CMJ030 (CC-4533; cw15, mt -) gDNA to confirm the insertion of the CIB1 cassette in *bst4.* **B.** Immunoblots confirming the production of BST4 in WT, *bst4, bst4::BST4* and *bst4::BST-truncated* Chlamydomonas lines. **C.** Immunoblot showing the absence of BST4 in the *bst4* knock-out line and presence in the WT and *bst4::BST4* complemented lines. **D.** Western blot used to quantify the amount of BST4 protein detected in WT and *bst4::BST4.* **E.** Immunoblot was used for the quantification of fluorescent intensity as a measure of BST4 protein abundance. All fluorescent intensity measurements were normalized to respective alpha-tubulin loading controls and conducted in triplicate. No statistical difference was observed between WT and *bst4::BST4* BST4 protein abundance (Paired two-tail t-test *p =* 0.22, n=3)

**Supplemental Figure 4.**
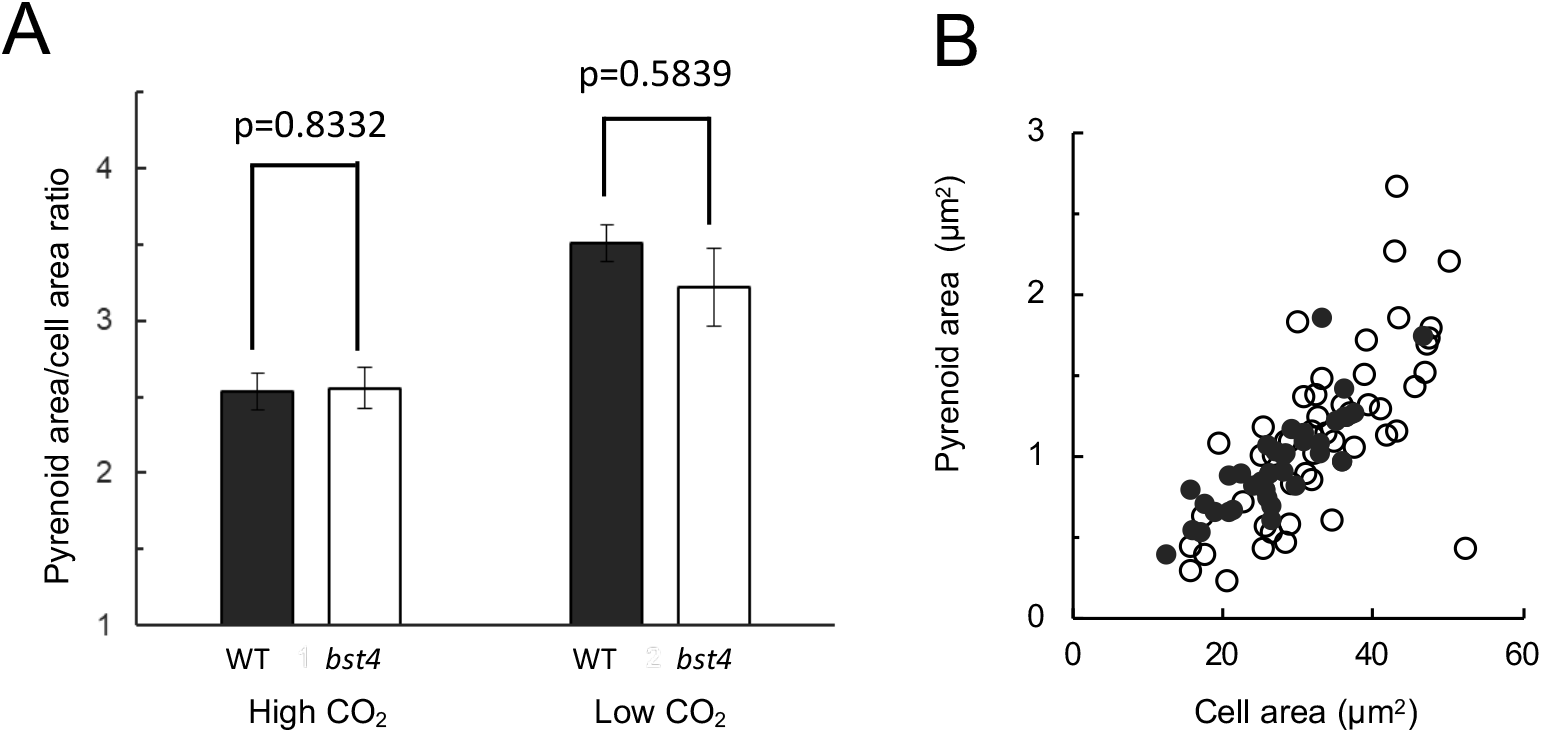
Pyrenoid morphology in bst4 vs WT. **A.**The pyrenoid area to cell area ratio of WT control strain (black) and *bst4* (white) strains at high (3%) and low (0.04%) CO2 concentrations. Each bar represents the mean value ± SEM (n=40–50). The data for wild type and *bst4* strains were compared using a two sample T-test. The p-values (showing no significant difference) are displayed above the bars**. B.** Pyrenoid area vs cell area of WT and *bst4* at 0.04% [CO2].

**Supplemental Figure 5.**
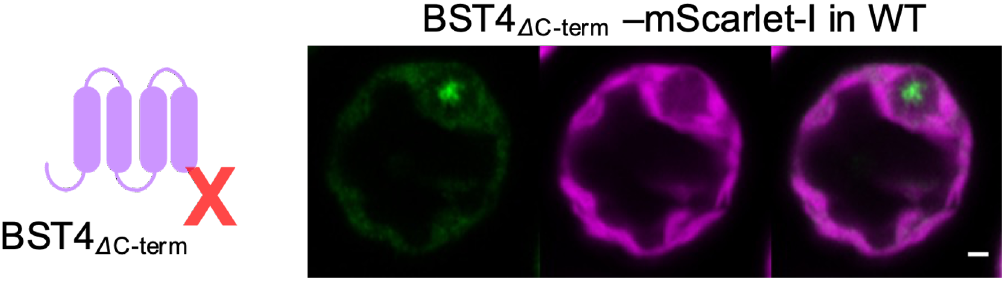
Localization of C-terminally truncated BST4 in WT background. BST4ΔCterm-mScarlet-I fluorescence shown in green and chlorophyll autofluorescence in magenta. Scale is bar 1 µm.

**Supplemental Figure 6.**
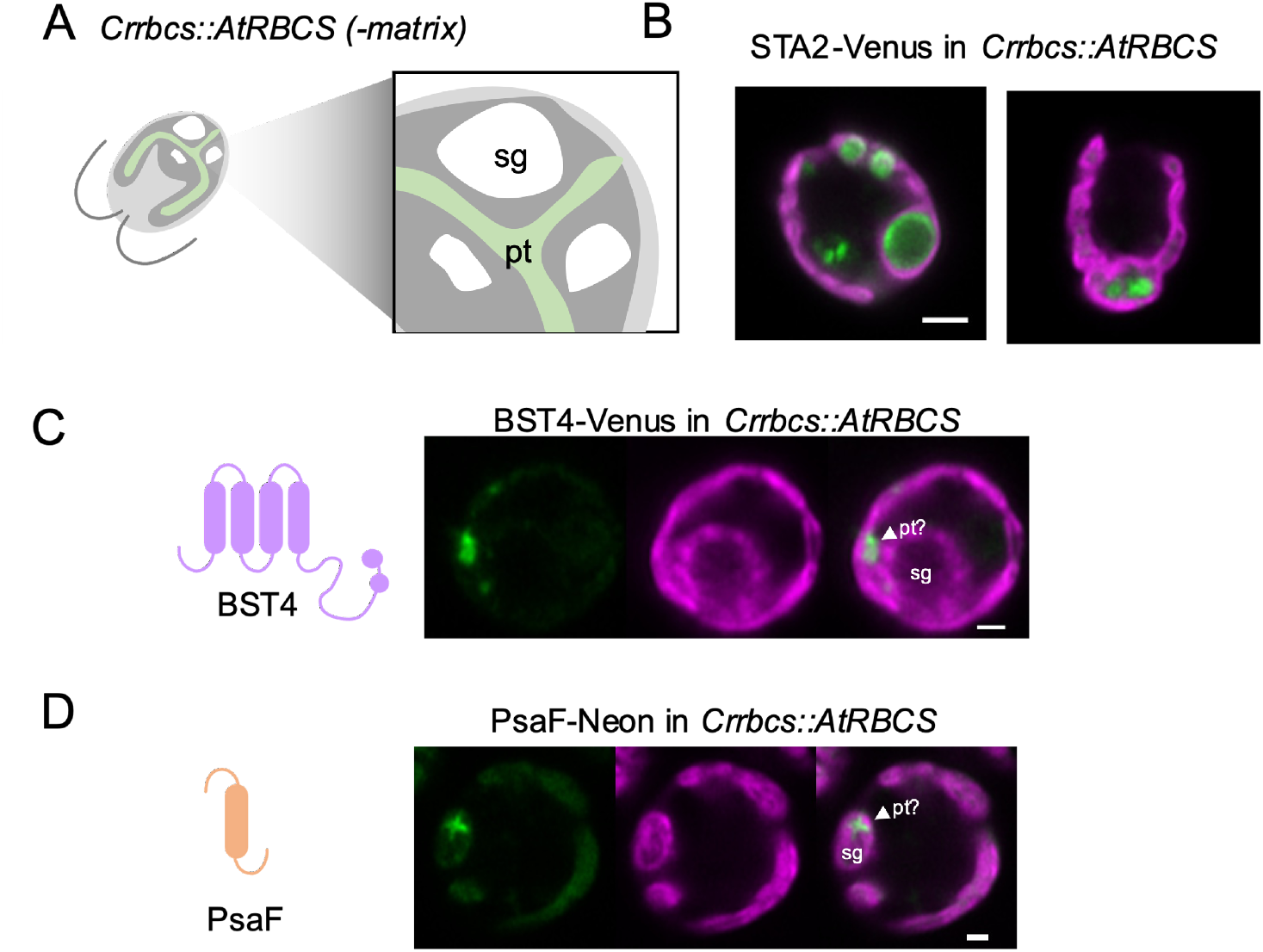
Localization of proteins in a pyrenoid matrix-less background. **A.**Diagram showing a section of the *Crrbcs::AtRBCS* matrix-less mutant (*-matrix*). The nascent pyrenoid tubules (pt) are shown in dark green, starch granules (sg) in white. **B.** Confocal image of starch binding protein STA2 fused to Venus in *Crrbcs::AtRBCS.* Scale bar is 2 µm. **C.** BST4-Venus in a WT Chlamydomonas cell. Scale bar is 1 µm. **D.** PsaF- mNeon in *Crrbcs::AtRBCS*. Venus and mNeonGreen fluorescence are shown in green and chlorophyll autofluorescence in magenta. Scale bar is 1 µm.

**Supplemental Figure 7.**
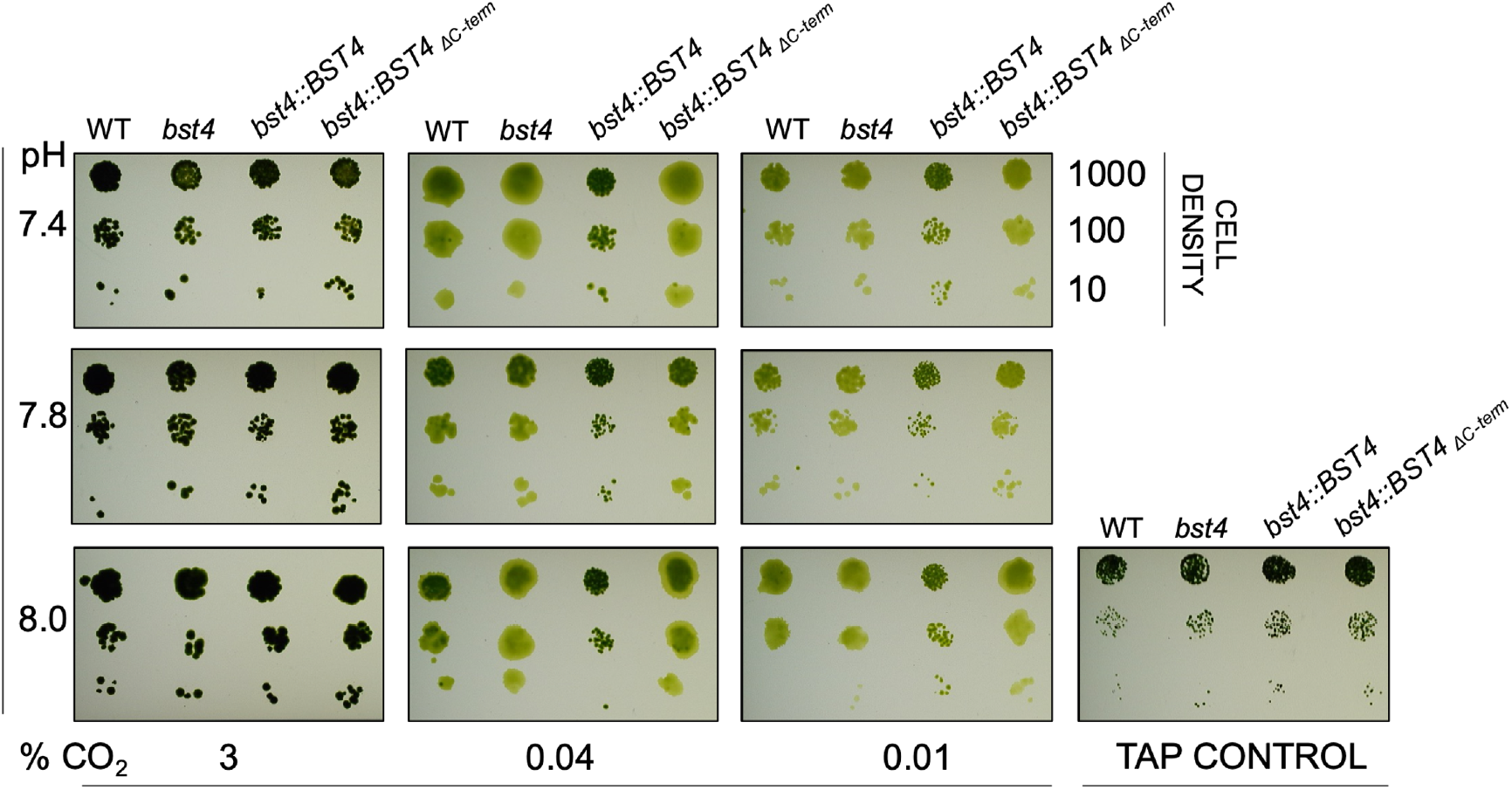
Spot test of WT, *bst4* and BST4 complemented lines under CCM induced conditions. *Chlamydomonas* strains were grown in serial dilution on agar plates in saturating light (400 μmol m^−2^ s^−1^) under a range of CO2 (+/- 2 ppm) and pH conditions (specified) to induce the CCM. All lines grew comparably to WT across the conditions used and on the low light TAP control plate.

**Supplemental Figure 8.**
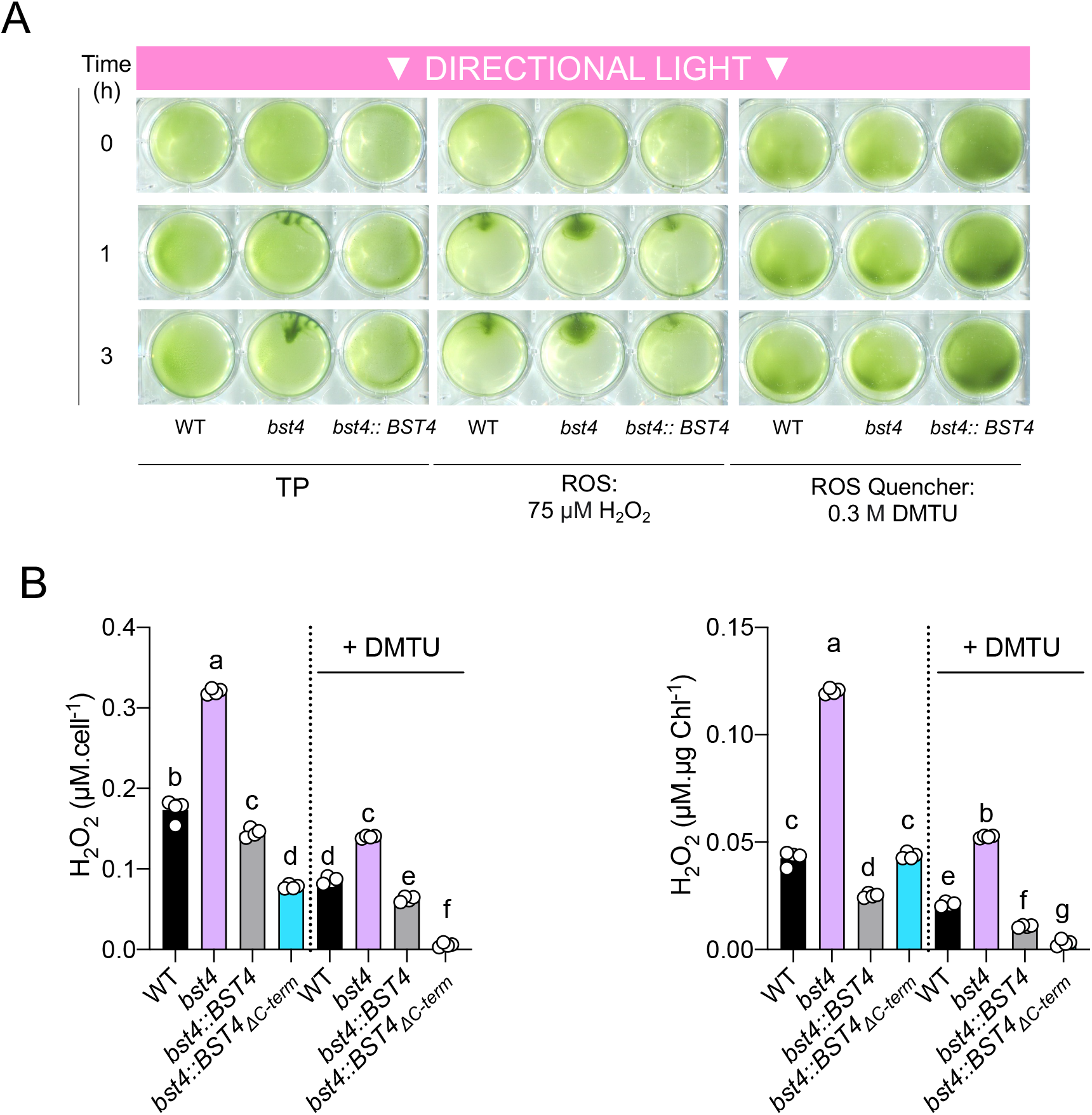
Phototaxis and ROS assay. **A.**TP liquid cultures of Chlamydomonas cells were uniformly distributed on 0.8% (w/v) agar and subjected to directional light (150 μmol m^−2^ s^−1^). Cell phototaxis was monitored at 0, 1, and 3 h. The assay was performed in the presence of 75 µM ROS hydrogen peroxide (H2O2) or 0.3 M ROS quencher N,N’-dimethylthiourea (DMTU). **B.** H2O2 assay. Chlamydomonas cells were grown in minimal TP liquid media and exposed to 150 µmol photons m^-2^s^-1^. A subset of cells were treated with the quencher DMTU. The concentration of H2O2 was subsequently quantified using Amplex Red (n=4), and is presented both proportionately to cell density and chlorophyll content. Different letters indicate significance (p<0.05) as determined by a one-way ANOVA and Tukey’s post-hoc test.

**Supplemental Figure 9.**
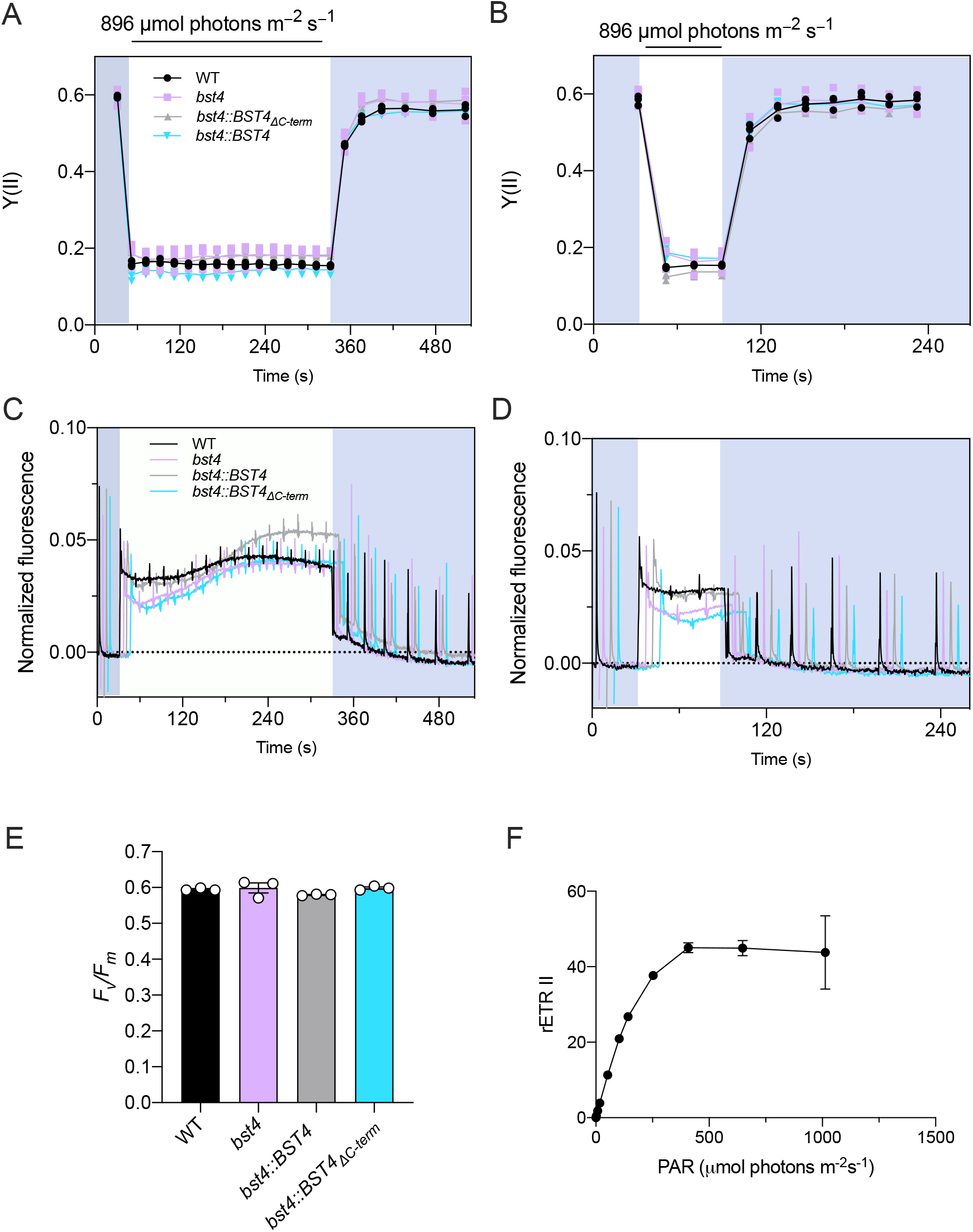
Chlorophyll fluorescence measurements. **A.**Y(II) during 5 min illumination and **B.** One minute illumination. **C.** and **D.** Raw fluorescence curves for A. and B., respectively. Curves are normalized to *Fo*. Genotypes are artificially spaced by 5 s for clarity. **E.** *Fv/Fm* measurements **F.** Light curve for WT in the presence of bicarbonate. Points represent the mean of three technical replicates ±SEM.

**Supplemental Figure 10.**
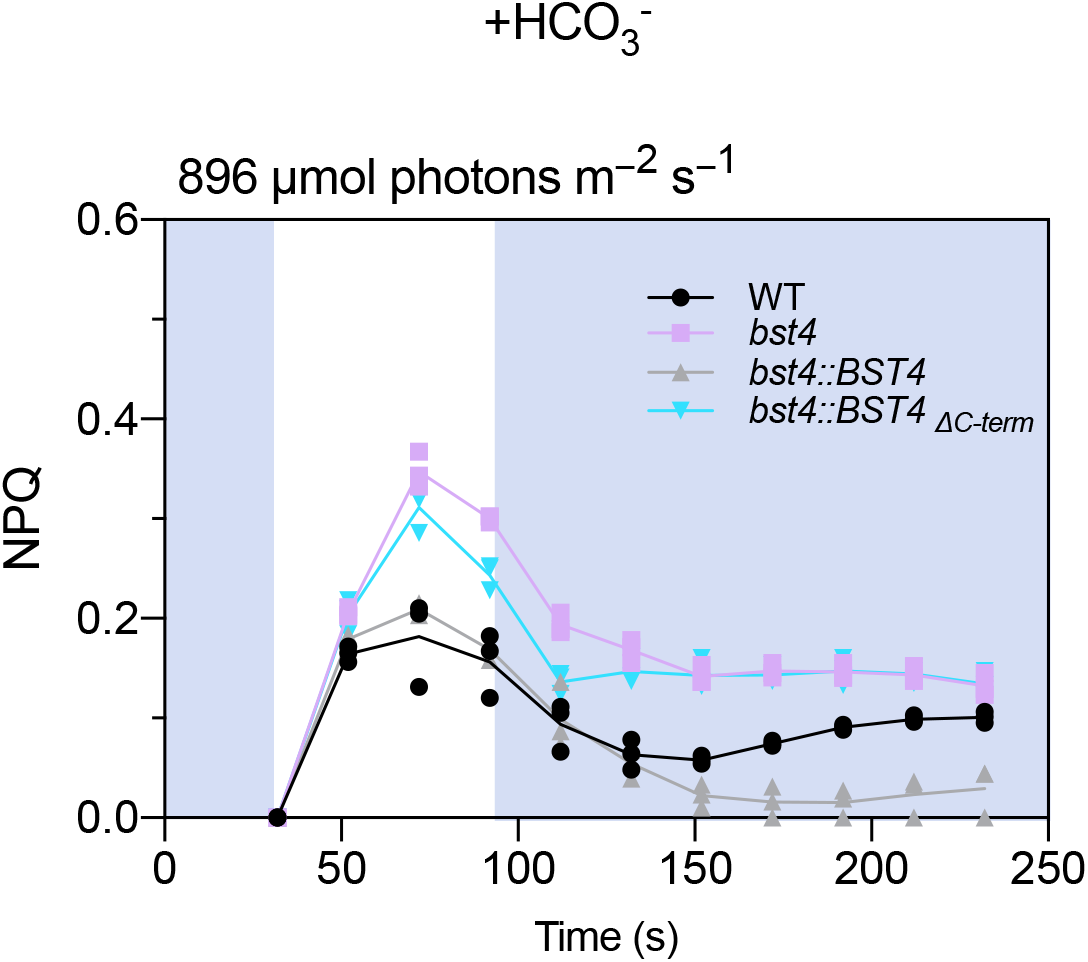
NPQ with supplemented bicarbonate. Wild type (WT) and mutants were grown in HS medium at 80 µmol photons m^−2^ s^−1^ and measured at 10 µg Chl ml^-1^. Cells were supplemented with 500 µM HCO3^-^ and then dark adapted for 5 min before the measurements. Dynamics of Non-photochemical photochemical quenching (NPQ) on transition from dark to high light. Kinetics for induction of chlorophyll fluorescence were recorded during 1 min of illumination at 896 µmol photons m^−2^ s^−1^ followed by 5 min in darkness.

**Supplemental Figure 11.**
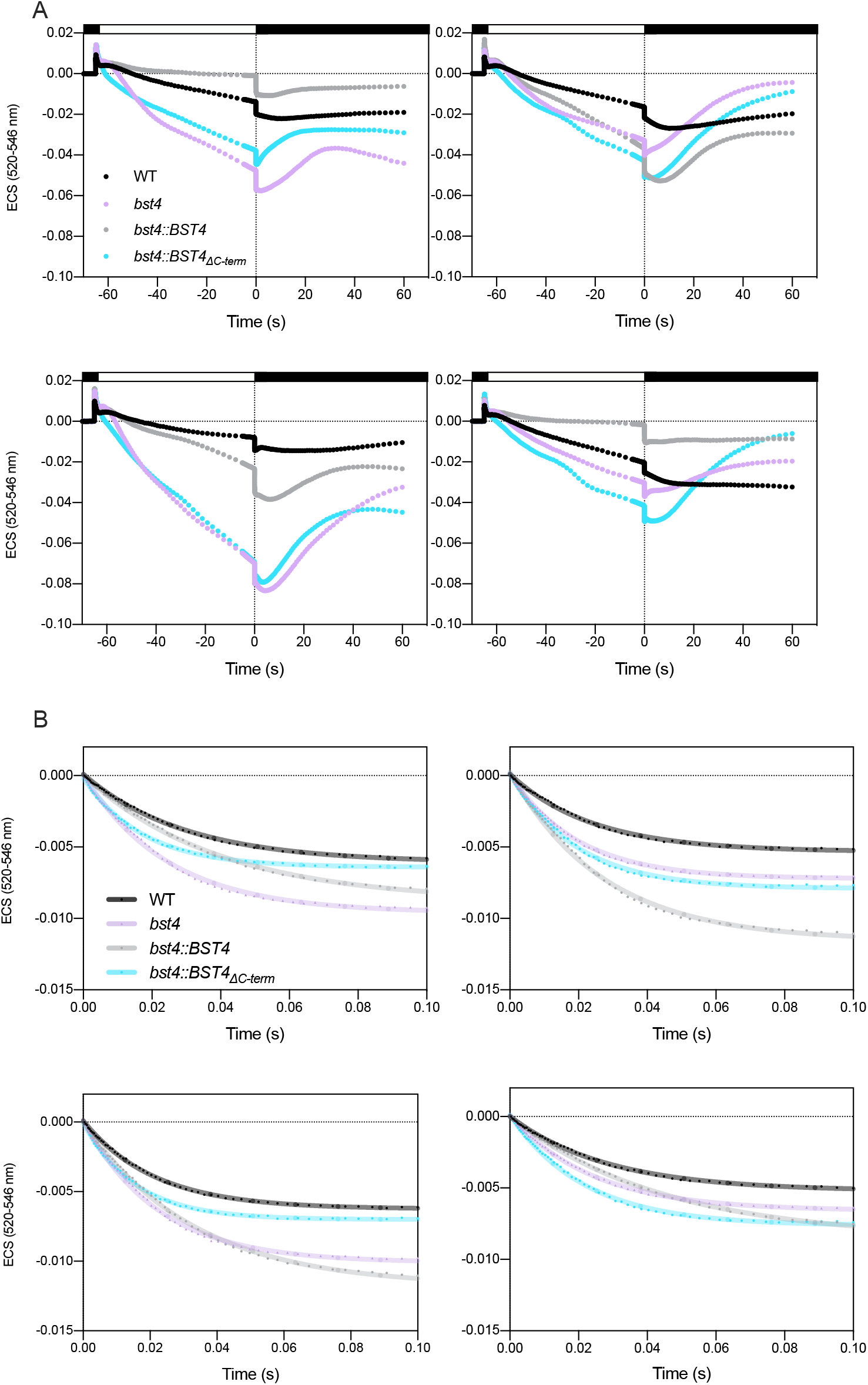
Electrochromic shift (ECS) traces across four biological repeats. **A.** Cells resuspended in HS at 150 µg Chl ml^-1^ were dark-adapted for 1 min and then illuminated for 1 min with 890 µmol photons m^−2^ s^−1^ after which the light was switched off to record ECS in darkness. We were unable to calculate the partitioning of the PMF due to non-canonical ECS slow kinetics. **B.** Normalized ECS decay curves. ECS decay of the first 100 ms was fitted to calculate gH^+^ (s^-1^) =1/time constant for decay. Data are the means of n=3 technical replicates.

**Supplemental Figure 12.**
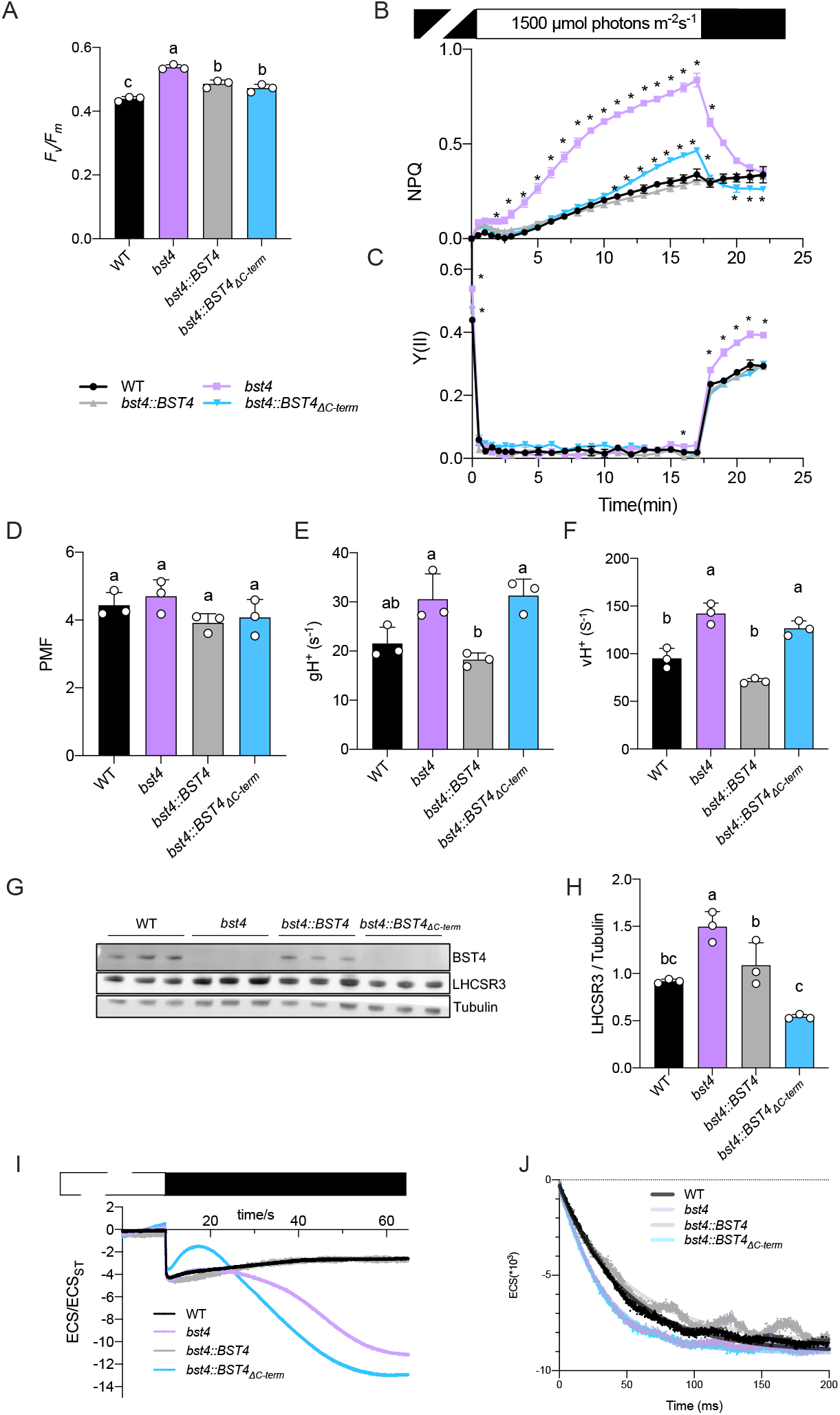
Chlamydomonas bst4 mutant has an enhanced NPQ and proton conductance under high light and limiting Ci conditions. Wild type (WT) and mutants were grown on TAP medium at 20 µmol photons m^−2^ s^−1^, resuspended in TP at 30 µg Chl ml^-1^ and exposed for 3 h to light at 150 µmol photons m^−2^ s^−1^. The cells were dark adapted for 1 h before the measurements. **A.** Maximum quantum yield of photosystem II. **B.** Dynamics of photosynthesis on transition from dark to high light. Kinetics for induction of chlorophyll fluorescence were recorded during 17 min of illumination at 1500 µmol photons m^−2^ s^−1^ followed by 5 min in darkness. Non-photochemical quenching (NPQ) and **C.** Photosystem II quantum yield (Y(II)). **D to F.** ECS decay kinetics were performed on cells pre-exposed for 10 min to high light and the **D.** total PMF, **E.** gH^+^, and **F.** total H^+^ flux (vH^+^) were determined as described in Methods. Data are the means ± SEM (n=3 replicates). **G.** Immunoblot of NPQ protein LHCSR3 in each genotype compared to tubulin after exposure of cells to 3 h 150 µmol photons m^−2^ s^−1^ in TP. **H.** Fluorescence of LHCSR3 protein band normalized to α-tubulin. Data are the means ± SEM (n=3 replicates). Different letters indicate statistically significant difference among the genotypes (one-way ANOVA test, followed by Tukey’s post hoc test, P < 0.05). **I.** Representative Electrochromic shift (ECS) curves used to determine PMF values in D. **J.** Representative ECS decay curves. To determine the gH+ parameter in E., ECS kinetics were recorded during 600 ms dark intervals. The ECS decay of the first 100 ms was fitted to calculate gH+ (s^-1^) =1/time

**Supplemental Figure 13.**
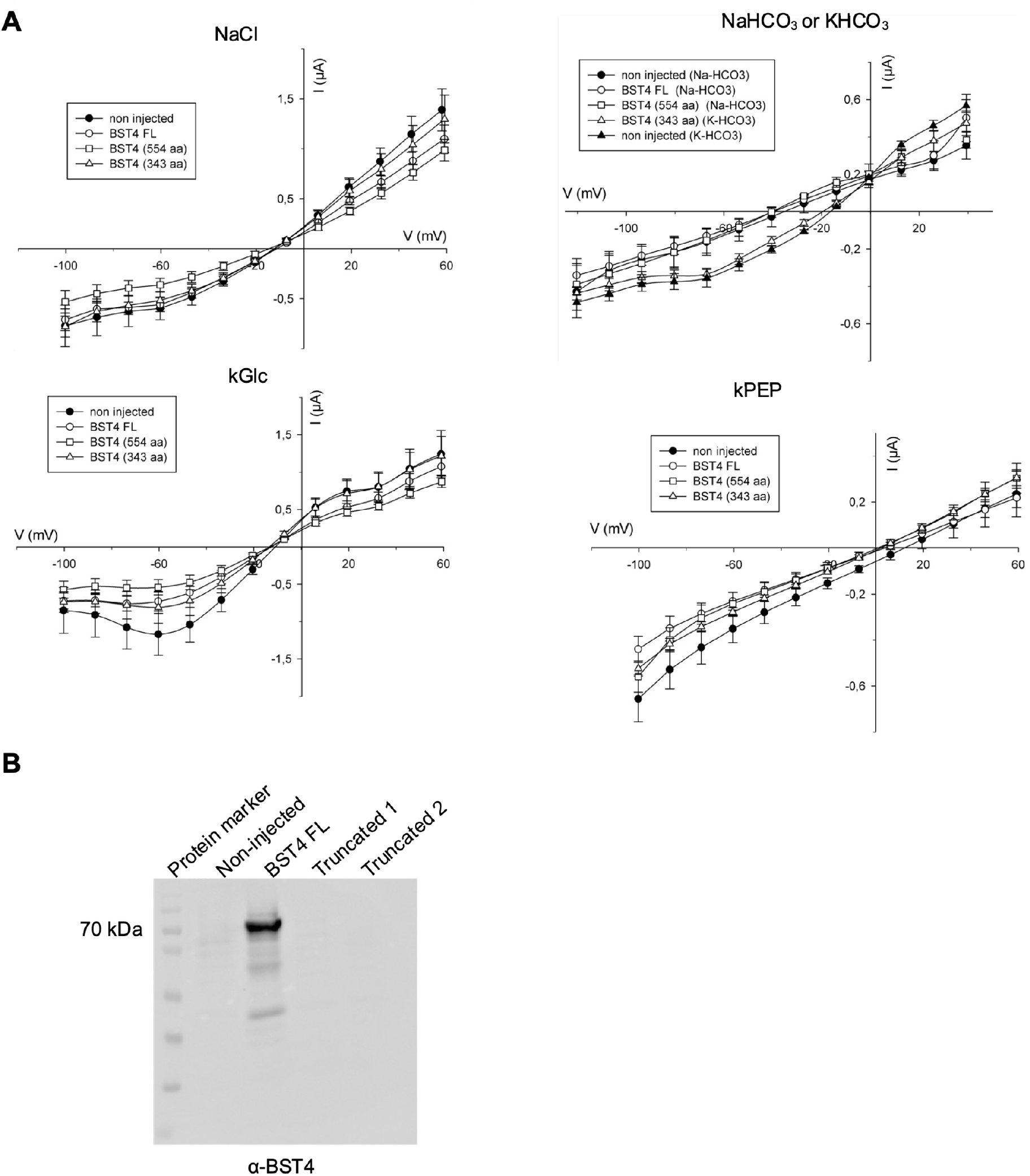
No currents were detected for BST4 with any anions tested in Xenopus oocytes. **A.** Steady state currents analysis of oocytes injected with BST4, full length or truncated, compared to non-injected oocytes. The voltage steps start from 60 mV to -100 mV (Cl^-^, PEP^-^, and Gluconate conditions) 40 mV to -120 mV (HCO3^-^ conditions, right panel), holding voltage is -20 mV. Recordings were performed on n>4 oocytes. There is no differences between the currents recorded in non-injected oocytes and expressing the protein. **B.** Western Blot analysis of oocytes. Lane 1 is the marker, lane 2 are non-injected oocytes, lane 3 is the full-length protein (about 66 kDa), lanes 3 and 4 are the truncated proteins. We can only detect full length BST4 as the antibody is directed against the C-terminal part of the protein that is removed in the two truncated versions of the channel.

**Supplemental Figure 14.**
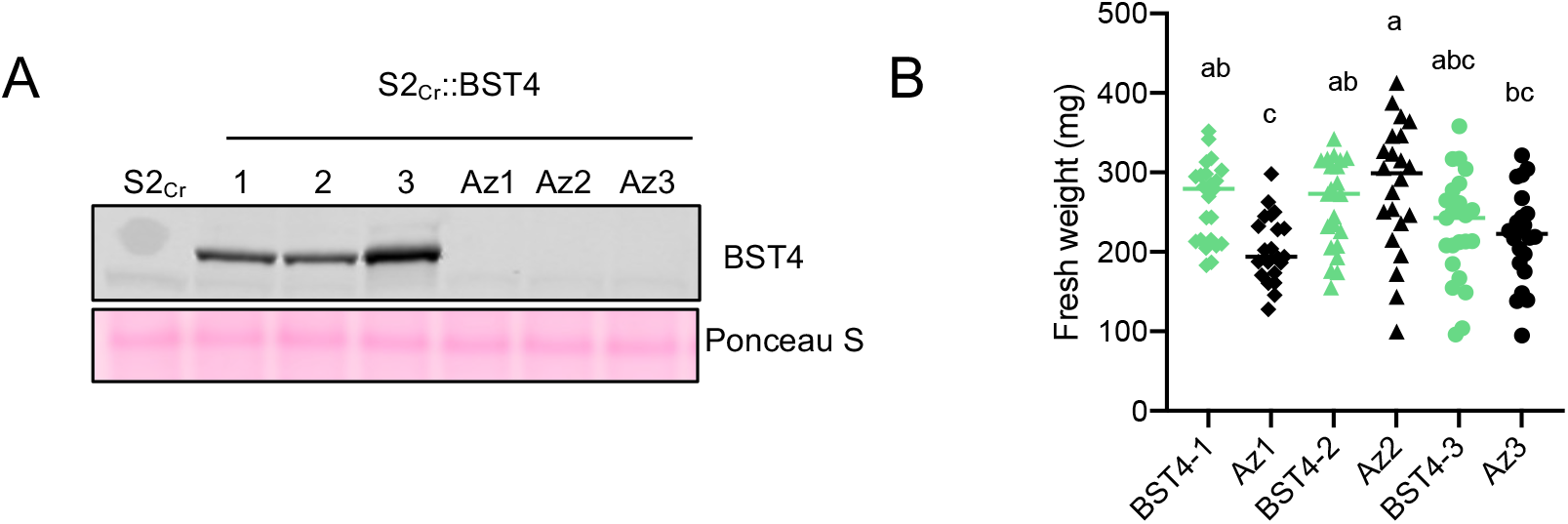
Phenotype of BST4 transgenic Arabidopsis line. **A.** Immunoblot against BST4 of proteins extracted from BST4 no tag lines and Azygous segregants (Az). S2Cr is the parent line. **B.** Fresh weight of 28-day old rosettes, bars represent mean weight for each genotype n=18-21. Different letters indicate statistically significant difference among the genotypes (one-way ANOVA test, followed by Turkey’s post hoc test, P < 0.05).

**Supplemental Figure 15.**
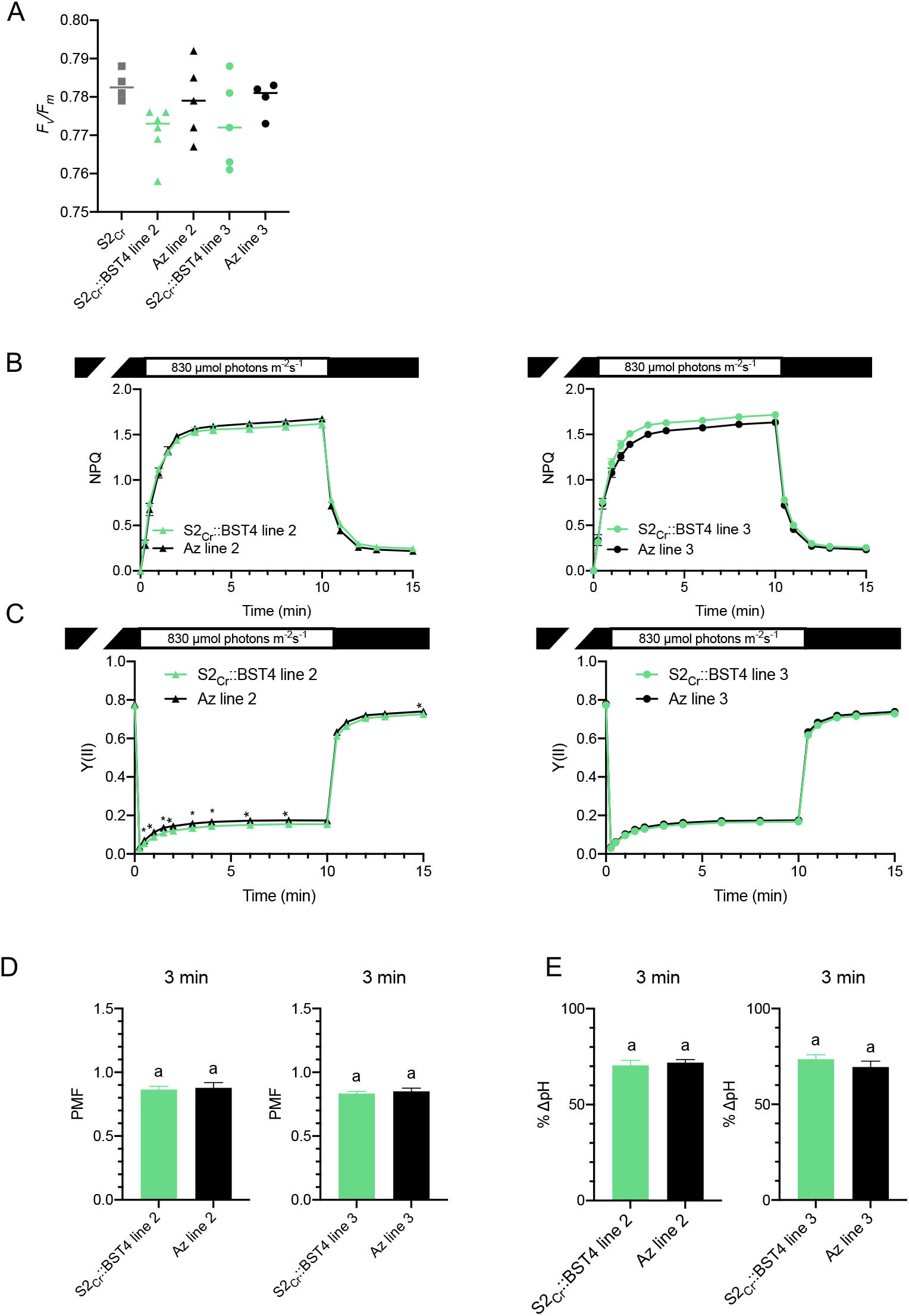
Photosynthetic measurements of BST4 transgenic Arabidopsis lines 2 and 3. **A.** *F*v/*F*m values measured on attached, 30 min dark adapted leaves of 8-week-old plants (n=15-21). The letters in indicate nonsignificant differences between plants expressing BST4 and their azygous segregants using Tukey post hoc test (P > 0.05). **B.** Non-photochemical quenching (NPQ) as a measure of photoprotection and **C.** photosystem II quantum yield (Y(II)) were recorded during 10 min of illumination at 830 µmol photons m^−2^ s^−1^ followed by a 5-minute dark period. Data are presented as means ± SEM (n=4-6). Asterisks indicate statistical difference between plants expressing BST4 and their Azygous (Az) segregants according to unpaired t-test (P ≤ 0.05). **D.** Proton motive force (PMF) size and **E.** partitioning to pH gradient (ΔpH) after 3 min illumination at 830 µmol photons m^−2^ s^−1^. Data are means ± SEM (n=5-6). The letters in E and F indicate non-significant differences between plants expressing BST4 and their azygous segregants according to unpaired t-test (P > 0.05).

**Supplemental Figure 16.**
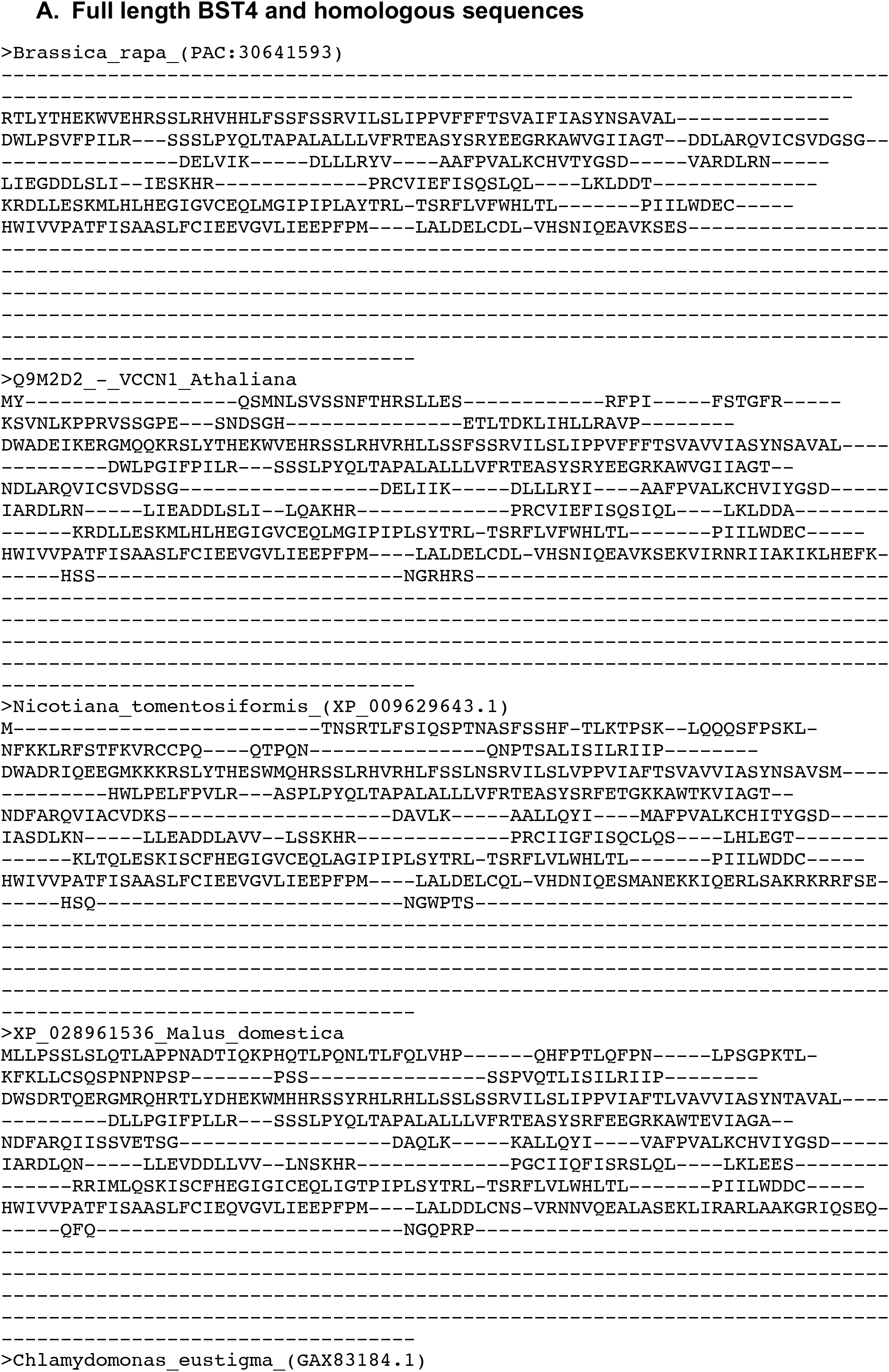
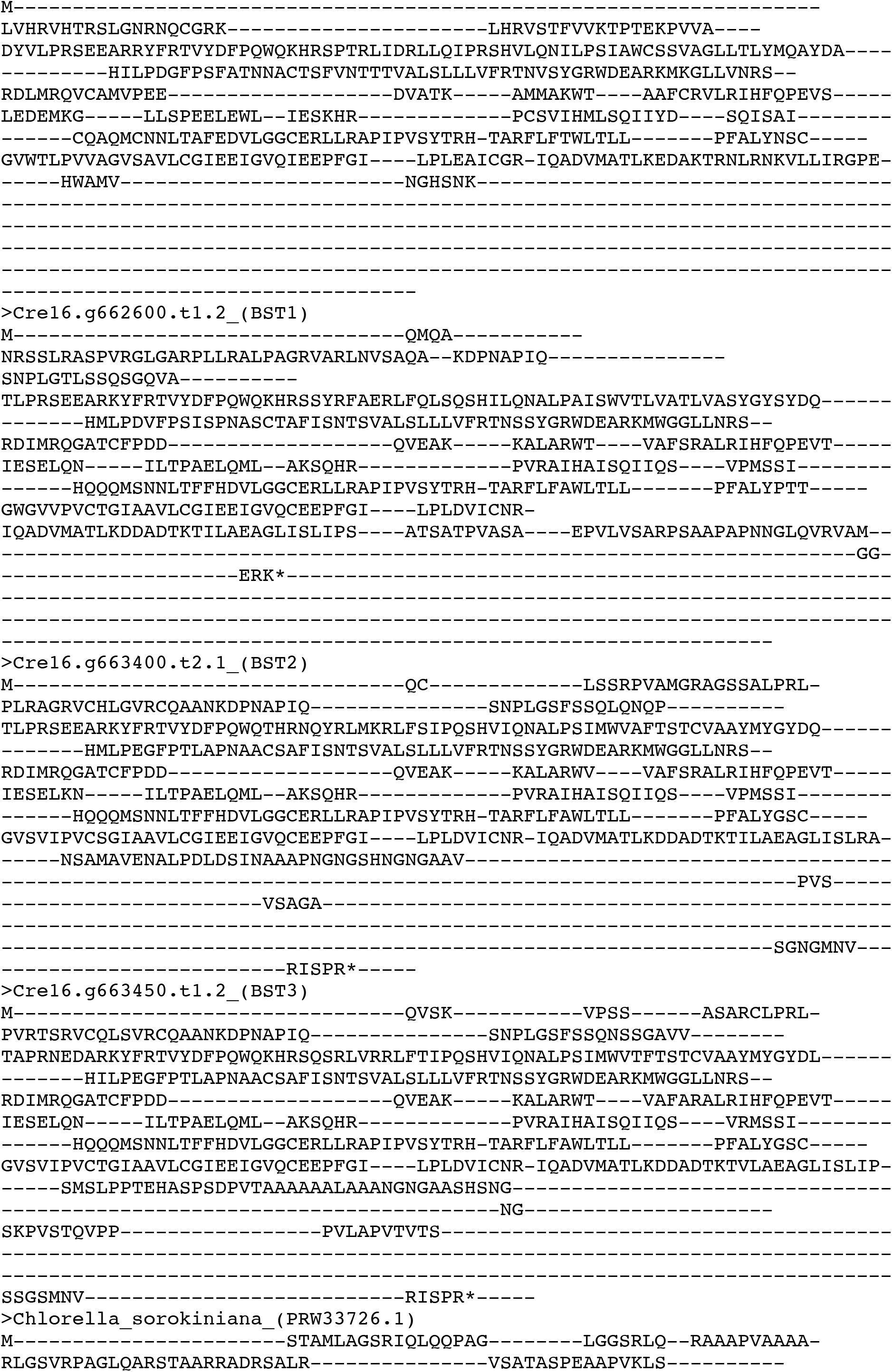
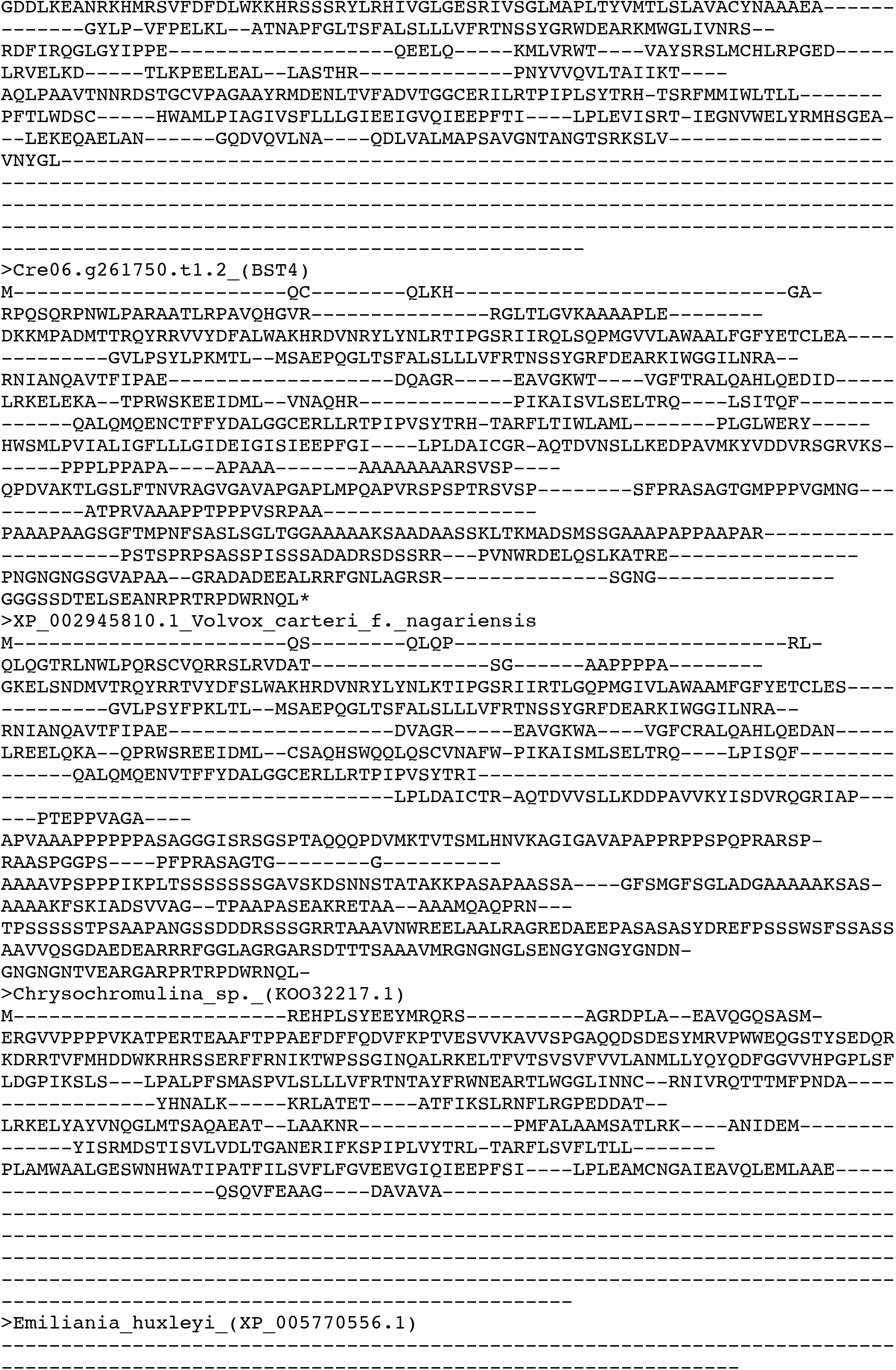
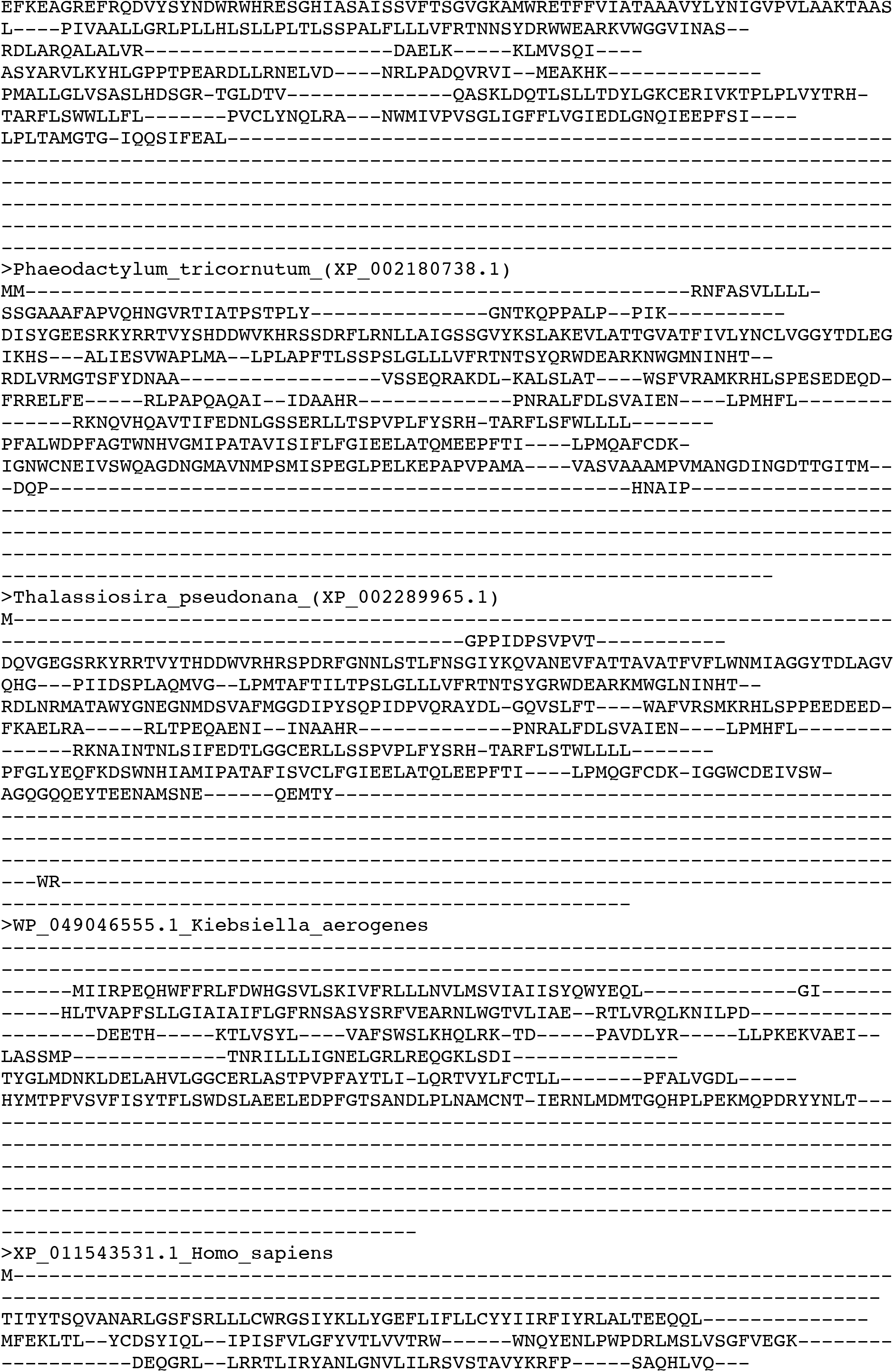
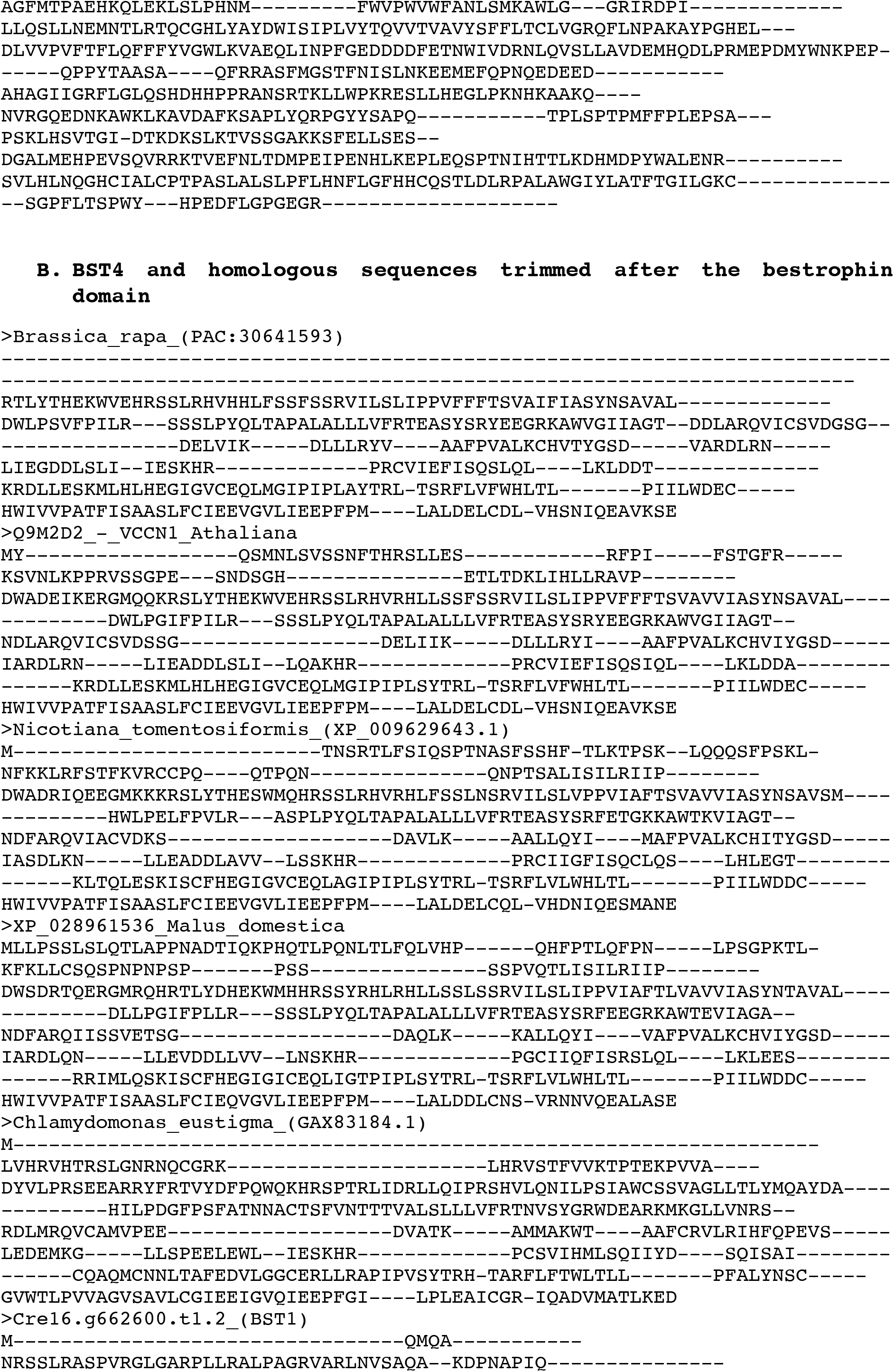
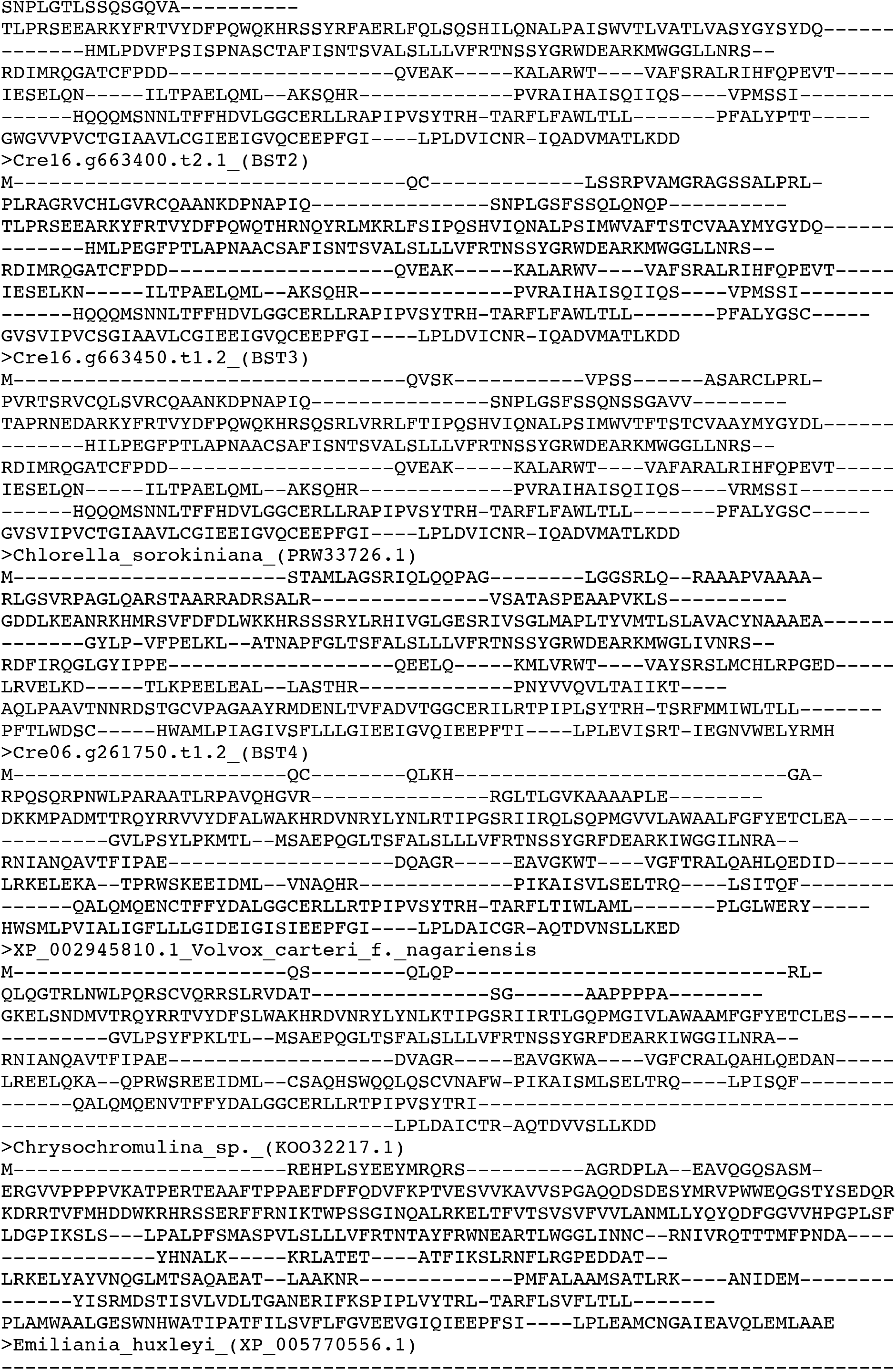
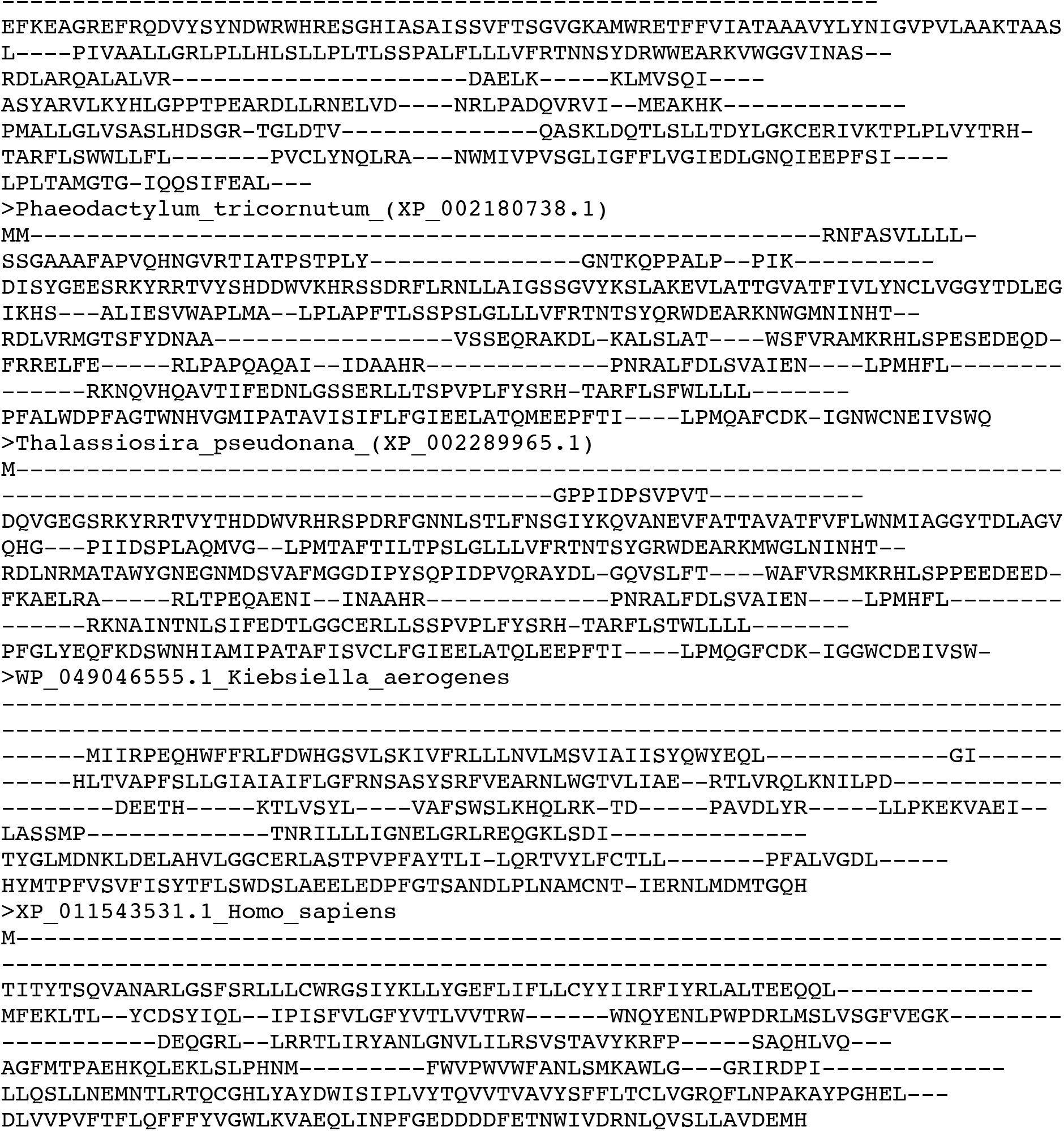
Sequences used to generate MAFFT alignments used to generate the phylogenetic trees included in this manuscript. FASTA files of MAFFT alignments of **A.** Full length and **B.** trimmed after the bestrophin domain BST4 and homologous amino acid sequences.

