## Supplemental Table and Figures for "The role of BST4 in the pyrenoid of *Chlamydomonas reinhardtii*"

### Supplemental Figures

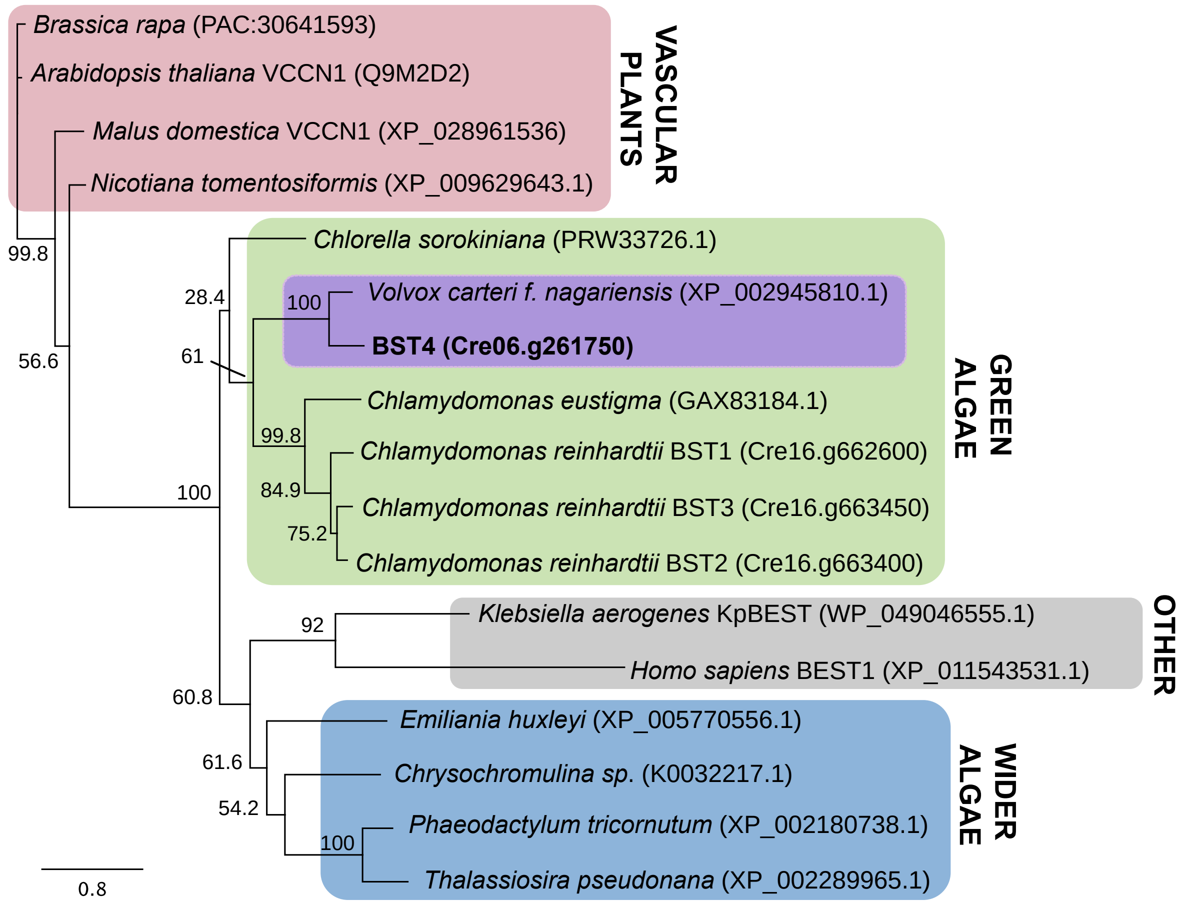


**Supplemental Figure 1. BST4 structure and sequence compared to other bestrophins.** Phylogenetic analysis of the full length Chlamydomonas BST4 amino acid sequence (bold). The evolutionary history of BST4 was inferred by using the maximum likelihood method based on the Le and Gascuel substitution model with discrete Gamma distribution (5 categories) and 500 bootstrap replicates. The tree is drawn to scale, with branch lengths measured in the number of substitutions per site.

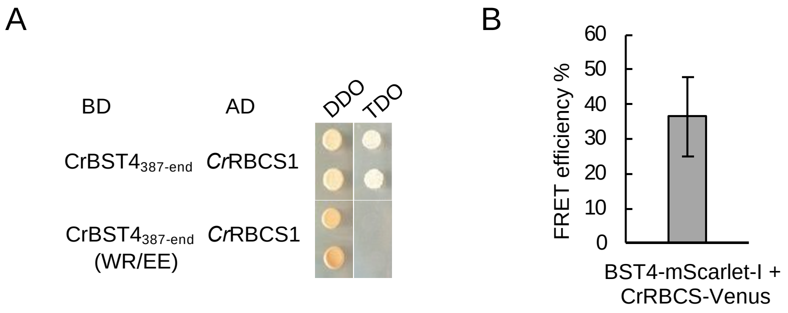


**Supplemental Figure 2. Interaction between BST4 and RBCS. A.** Yeast-2-hybrid experiment. The disordered region of BST4 C-terminus (amino acid 387-end) and with residues WR from RBMs are changed to EE. Abbreviations: AD, activation domain; BD, binding domain; DDO, double drop out media; TDO, triple drop out media. Growth on TDO indicates an interaction. **B**. FRET efficiency between BST4-mScarlet-I and CrRBCS-Venus. Sensitized-FRET measurement on dual-tagged cells (BST4-Venus and RBCS1-mCherry) showed a moderate FRET efficiency of 35% (n=10).

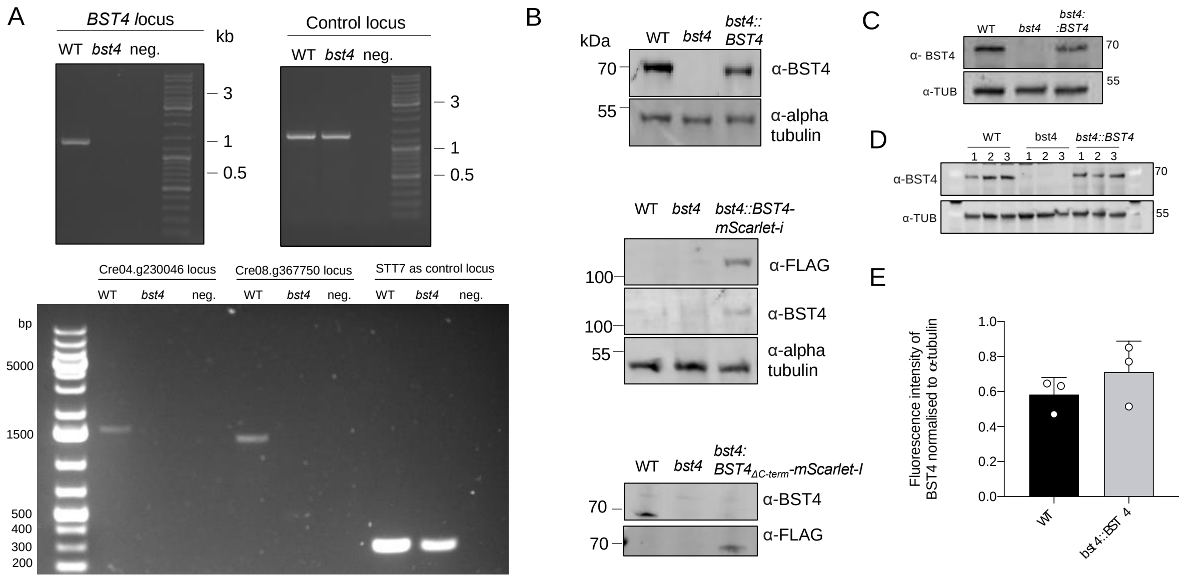


**Supplemental Figure 3. Validation of Chlamydomonas BST4 lines**. **A.** PCR amplification of the mapped CIB1 insertion sites and control loci of *bst4* (LMJ.RY0402.159478) and WT control strain (CMJ030 (CC-4533; cw15, mt -) gDNA to confirm the insertion of the CIB1 cassette in *bst4.* **B.** Immunoblots confirming the production of BST4 in WT, *bst4, bst4::BST4* and *bst4::BST-truncated* Chlamydomonas lines. **C.** Immunoblot showing the absence of BST4 in the *bst4* knock-out line and presence in the WT and *bst4::BST4* complemented lines. **D.** Western blot used to quantify the amount of BST4 protein detected in WT and *bst4::BST4.* **E.** Immunoblot was used for the quantification of fluorescent intensity as a measure of BST4 protein abundance. All fluorescent intensity measurements were normalized to respective alpha-tubulin loading controls and conducted in triplicate. No statistical difference was observed between WT and *bst4::BST4* BST4 protein abundance (Paired two-tail t-test $p$ = 0.22, n=3)

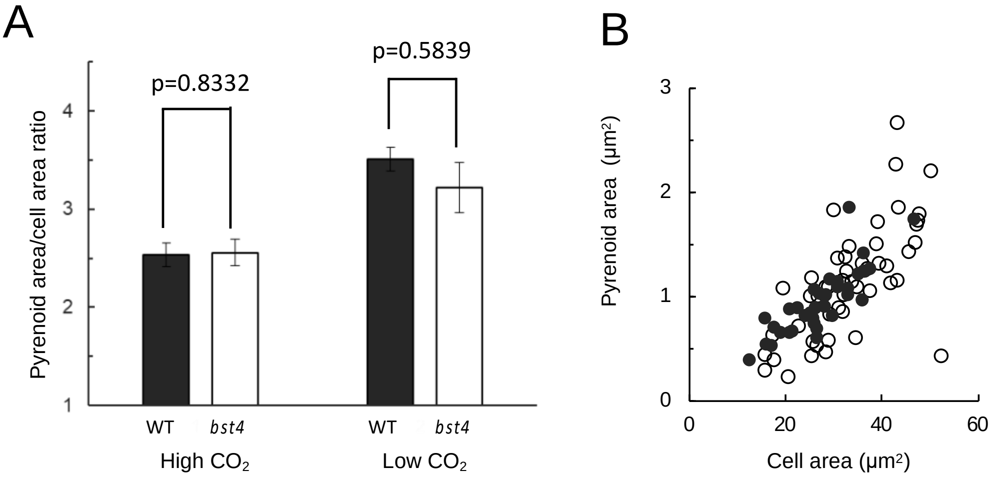


**Supplemental Figure 4. Pyrenoid morphology in bst4 vs WT. A.** The pyrenoid area to cell area ratio of WT control strain (black) and *bst4* (white) strains at high (3%) and low (0.04%) $CO_2$ concentrations. Each bar represents the mean value ± SEM (n=40–50). The data for wild type and *bst4* strains were compared using a two sample T-test. The p-values (showing no significant difference) are displayed above the bars**. B.** Pyrenoid area vs cell area of WT and *bst4* at 0.04% [$CO_2$].

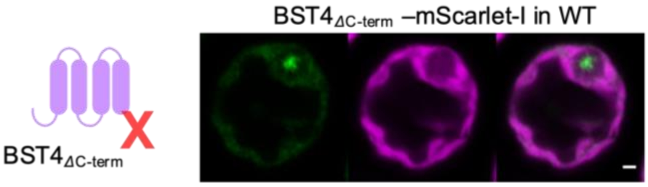


**Supplemental Figure 5. Localization of C-terminally truncated BST4 in WT background.** BST4$_{\Delta Cterm}$-mScarlet-I fluorescence shown in green and chlorophyll autofluorescence in magenta. Scale is bar 1 µm.

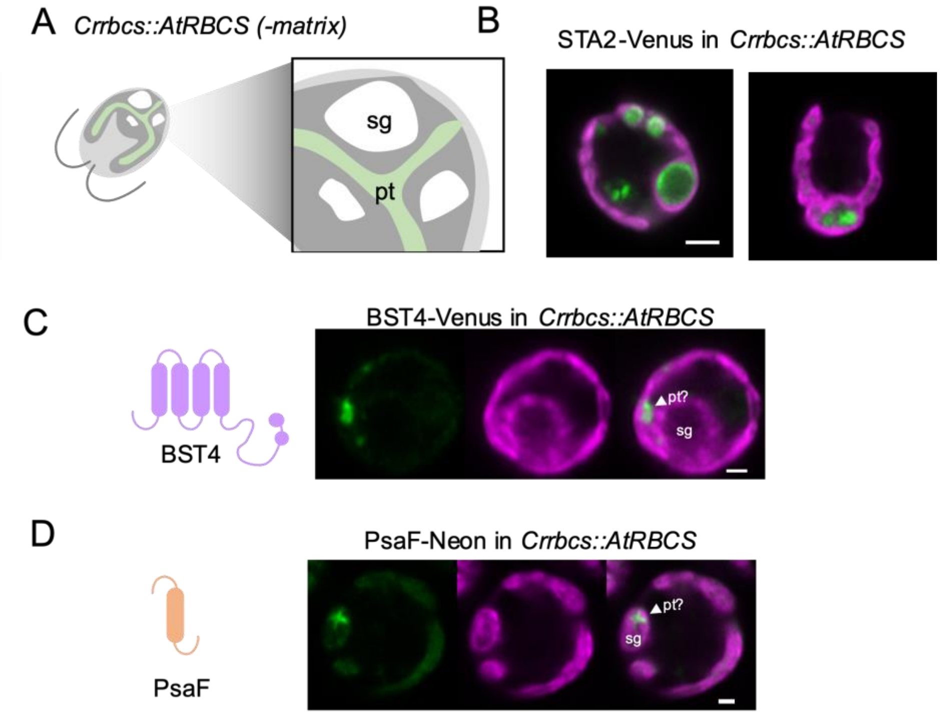


**Supplemental Figure 6. Localization of proteins in a pyrenoid matrix-less background. A.** Diagram showing a section of the *Crrbcs::AtRBCS* matrix-less mutant (-*matrix*). The nascent pyrenoid tubules (pt) are shown in dark green, starch granules (sg) in white. **B.** Confocal image of starch binding protein STA2 fused to Venus in *Crrbcs::AtRBCS.* Scale bar is 2 µm. **C.** BST4-Venus in a WT Chlamydomonas cell. Scale bar is 1 µm. **D.** PsaF-mNeon in *Crrbcs::AtRBCS*. Venus and mNeonGreen fluorescence are shown in green and chlorophyll autofluorescence in magenta. Scale bar is 1 µm.

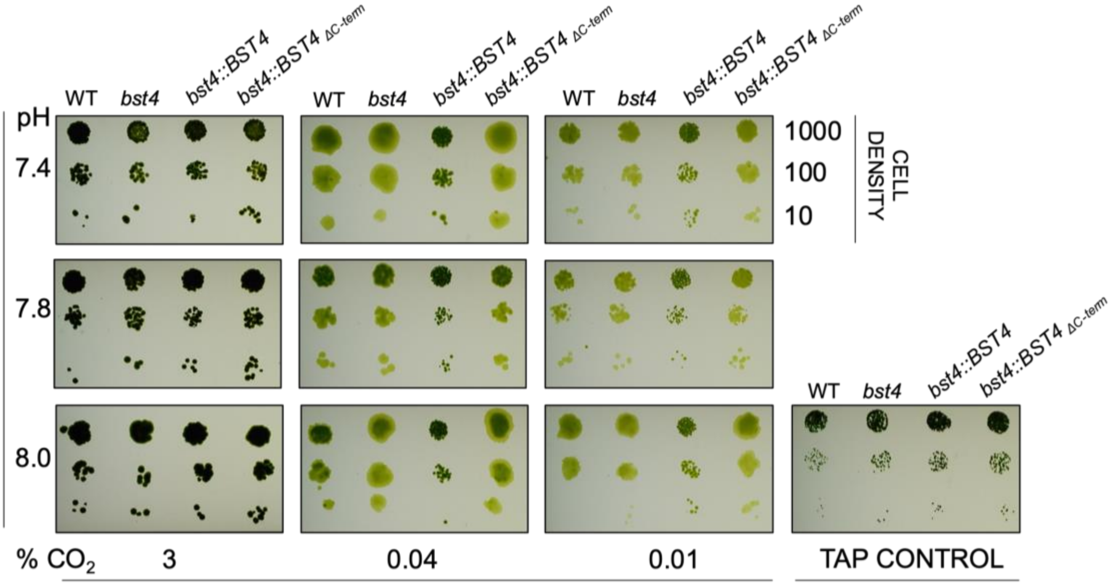


**Supplemental Figure 7. Spot test of WT, *bst4* and BST4 complemented lines under CCM induced conditions*.*** *Chlamydomonas* strains were grown in serial dilution on agar plates in saturating light (400 µmol $m^{-2}$ $s^{-1}$) under a range of $CO_2$ (+/- 2 ppm) and pH conditions (specified) to induce the CCM. All lines grew comparably to WT across the conditions used and on the low light TAP control plate.

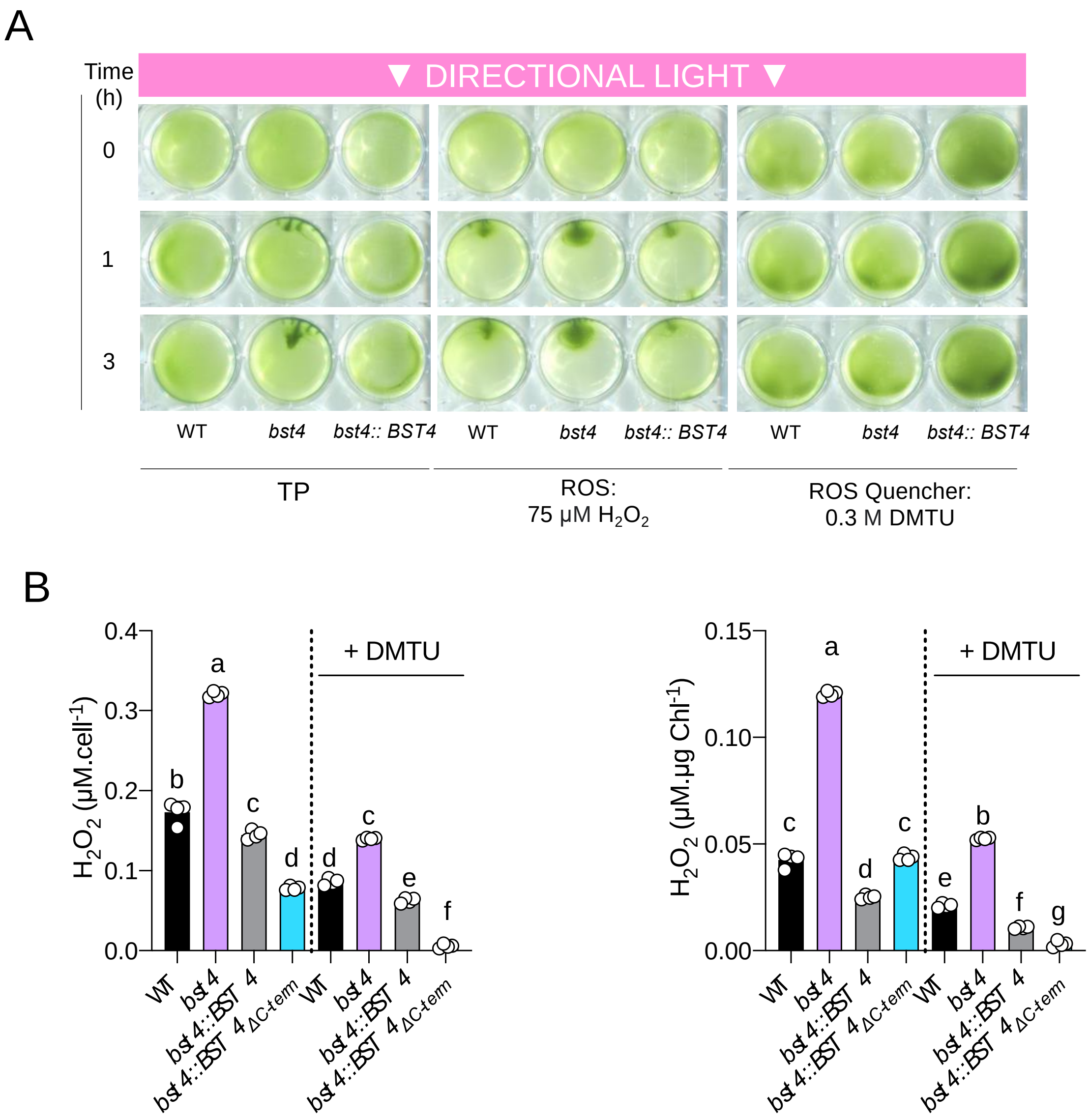


**Supplemental Figure 8. Phototaxis and ROS assay. A.** TP liquid cultures of Chlamydomonas cells were uniformly distributed on 0.8% (w/v) agar and subjected to directional light (150 µmol $m^{-2}$ $s^{-1}$). Cell phototaxis was monitored at 0, 1, and 3 h. The assay was performed in the presence of 75 µM ROS hydrogen peroxide ($H_2O_2$) or 0.3 M ROS quencher N,N'-dimethylthiourea (DMTU). **B.** $H_2O_2$ assay. Chlamydomonas cells were grown in minimal TP liquid media and exposed to 150 µmol photons $m^{-2}s^{-1}$. A subset of cells were treated with the quencher DMTU. The concentration of $H_2O_2$ was subsequently quantified using Amplex Red (n=4), and is presented both proportionately to cell density and chlorophyll content. Different letters indicate significance ($p<0.05$) as determined by a one-way ANOVA and Tukey's post-hoc test.

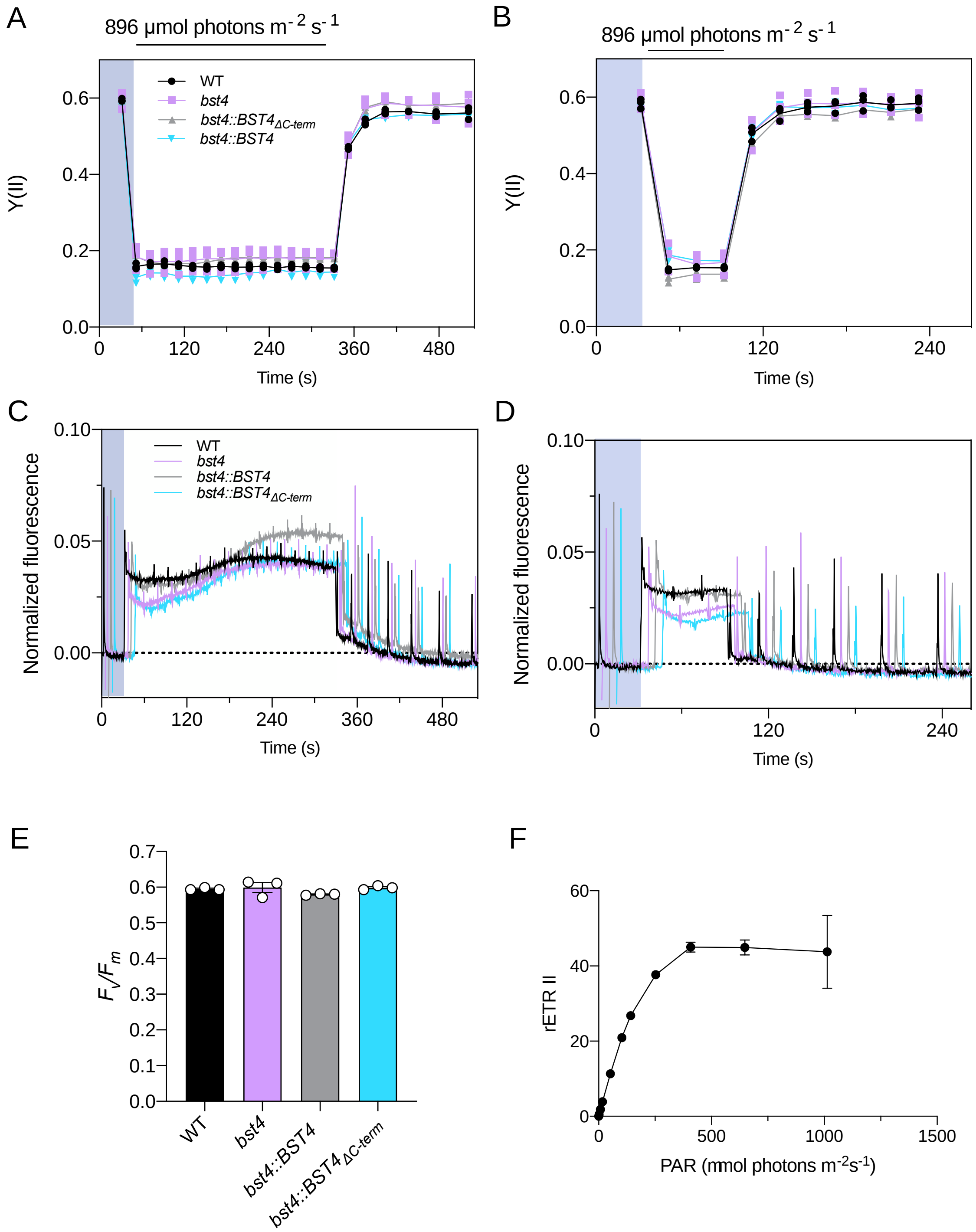


**Supplemental Figure 9. Chlorophyll fluorescence measurements. A.** Y(II) during 5 min illumination and **B.** One minute illumination. **C.** and **D.** Raw fluorescence curves for A. and B., respectively. Curves are normalized to $F_o$. Genotypes are artificially spaced by 5 s for clarity. **E.** $F_v/F_m$ measurements **F.** Light curve for WT in the presence of bicarbonate. Points represent the mean of three technical replicates ±SEM.

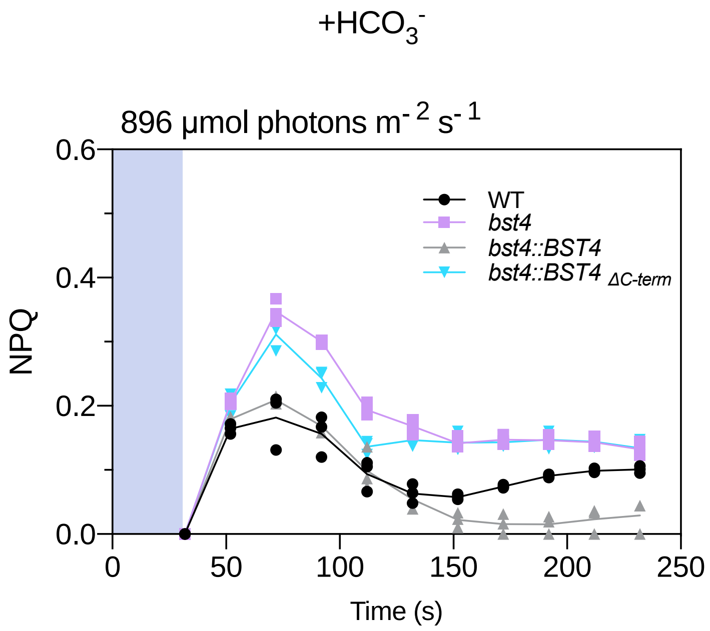


**Supplemental Figure 10. NPQ with supplemented bicarbonate.** Wild type (WT) and mutants were grown in HS medium at 80 µmol photons $m^{-2}$ $s^{-1}$ and measured at 10 µg Chl $ml^{-1}$. Cells were supplemented with 500 µM $HCO_3^-$ and then dark adapted for 5 min before the measurements. Dynamics of Non-photochemical photochemical quenching (NPQ) on transition from dark to high light. Kinetics for induction of chlorophyll fluorescence were recorded during 1 min of illumination at 896 µmol photons $m^{-2}$ $s^{-1}$ followed by 5 min in darkness.

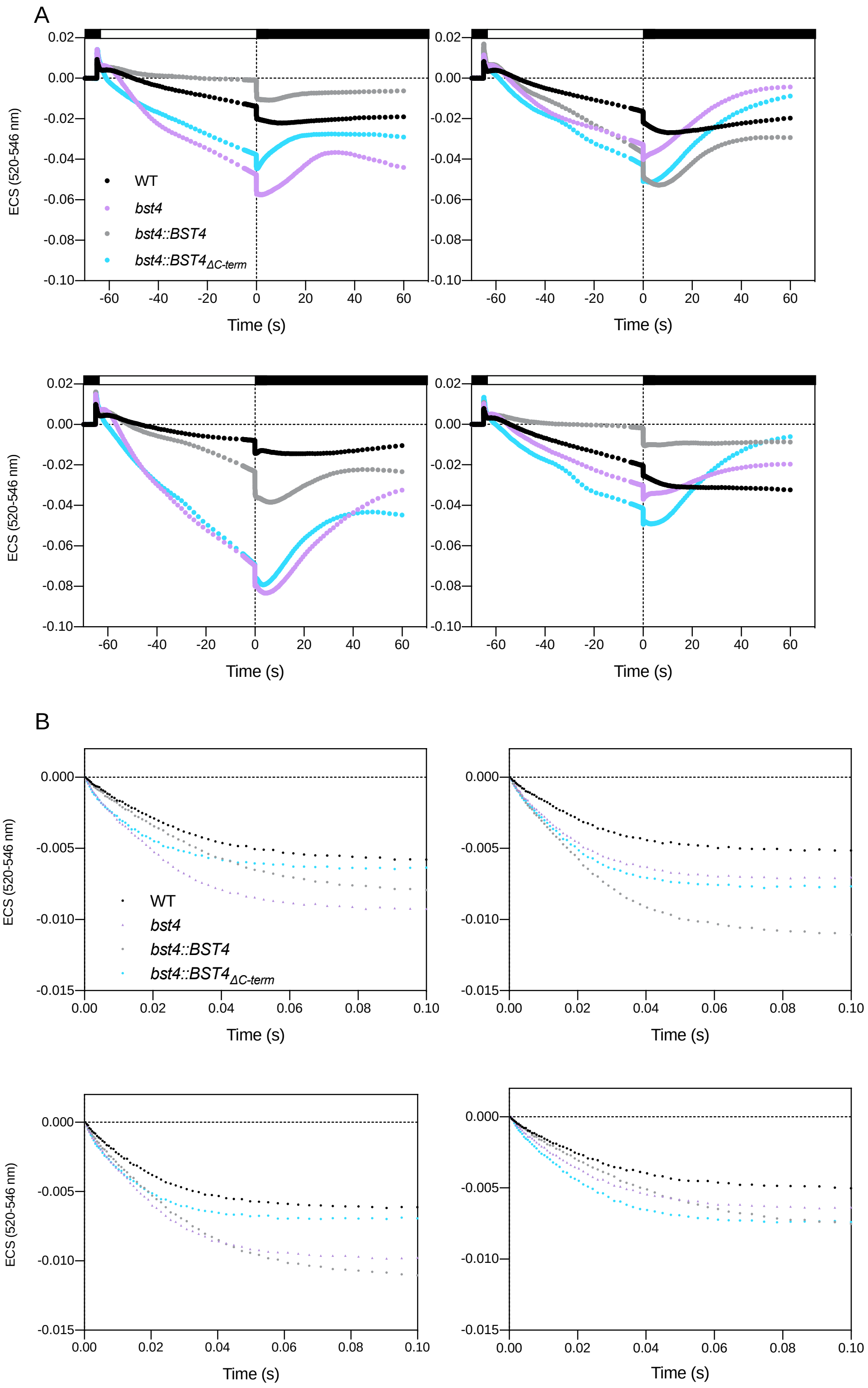


**Supplemental Figure 11.** Electrochromic shift (**ECS) traces across four biological repeats. A.** Cells resuspended in HS at 150 µg Chl ml$^{-1}$ were dark-adapted for 1 min and then illuminated for 1 min with 890 µmol photons m$^{-2}$ s$^{-1}$ after which the light was switched off to record ECS in darkness. We were unable to calculate the partitioning of the PMF due to non-canonical ECS slow kinetics. **B.** Normalized ECS decay curves. ECS decay of the first 100 ms was fitted to calculate $g_{H}^{+}$ (s$^{-1}$) =1/time constant for decay. Data are the means of n=3 technical replicates.

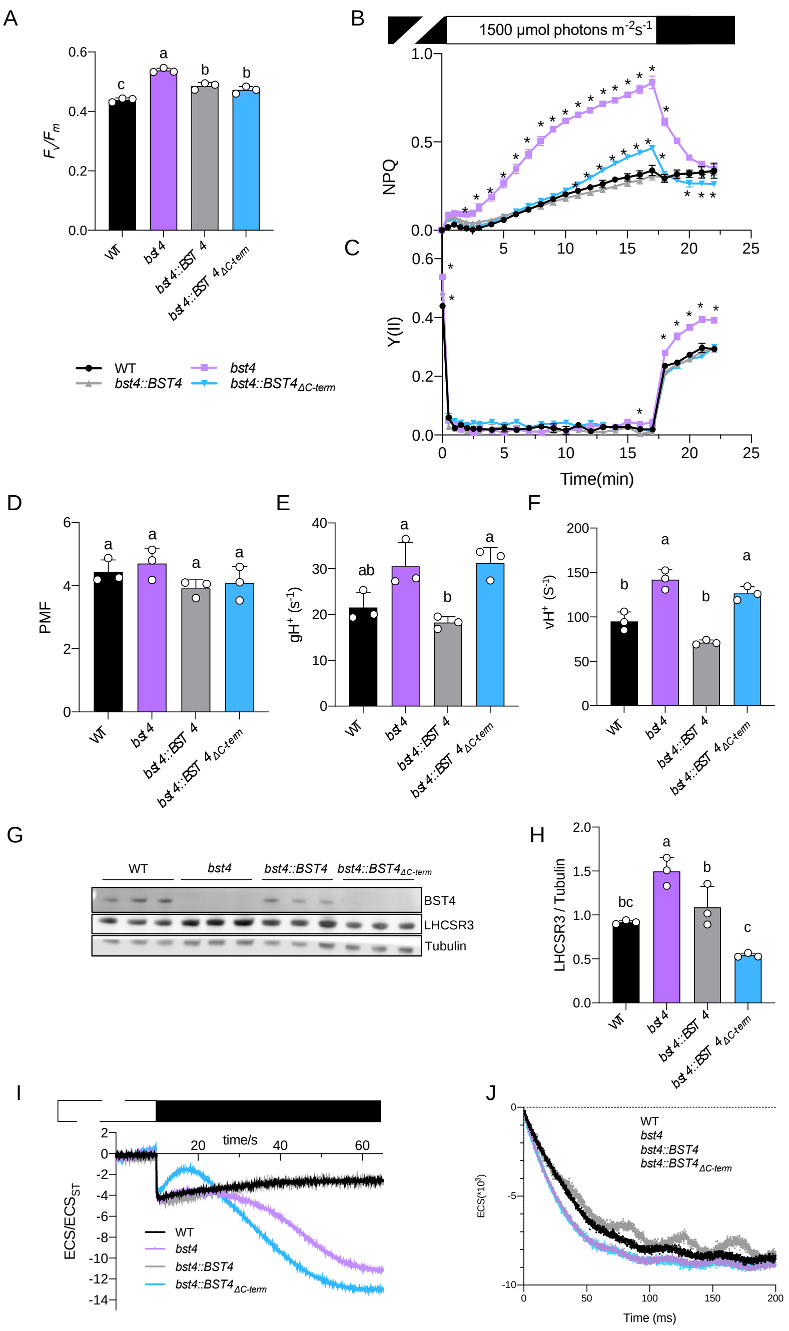


**Supplemental Figure 12. Chlamydomonas bst4 mutant has an enhanced NPQ and proton conductance under high light and limiting Ci conditions.** Wild type (WT) and mutants were grown on TAP medium at 20 µmol photons $m^{-2}$ $s^{-1}$, resuspended in TP at 30 µg Chl $ml^{-1}$ and exposed for 3 h to light at 150 µmol photons $m^{-2}$ $s^{-1}$. The cells were dark adapted for 1 h before the measurements. **A.** Maximum quantum yield of photosystem II. **B.** Dynamics of photosynthesis on transition from dark to high light. Kinetics for induction of chlorophyll fluorescence were recorded during 17 min of illumination at 1500 µmol photons $m^{-2}$ $s^{-1}$ followed by 5 min in darkness. Non-photochemical quenching (NPQ) and **C.** Photosystem II quantum yield (Y(II)). **D to F.** ECS decay kinetics were performed on cells pre-exposed for 10 min to high light and the **D.** total PMF, **E.** $g_{H^+}$, and **F.** total $H^+$ flux ($v_{H^+}$) were determined as described in Methods. Data are the means ± SEM (n=3 replicates). **G.** Immunoblot of NPQ protein LHCSR3 in each genotype compared to tubulin after exposure of cells to 3 h 150 µmol photons $m^{-2}$ $s^{-1}$ in TP. **H.** Fluorescence of LHCSR3 protein band normalized to α-tubulin. Data are the means ± SEM (n=3 replicates). Different letters indicate statistically significant difference among the genotypes (one-way ANOVA test, followed by Tukey's post hoc test, P < 0.05). **I.** Representative Electrochromic shift (ECS) curves used to determine PMF values in D. **J.** Representative ECS decay curves. To determine the gH+ parameter in E., ECS kinetics were recorded during 600 ms dark intervals. The ECS decay of the first 100 ms was fitted to calculate gH+ ($s^{-1}$) =1/time

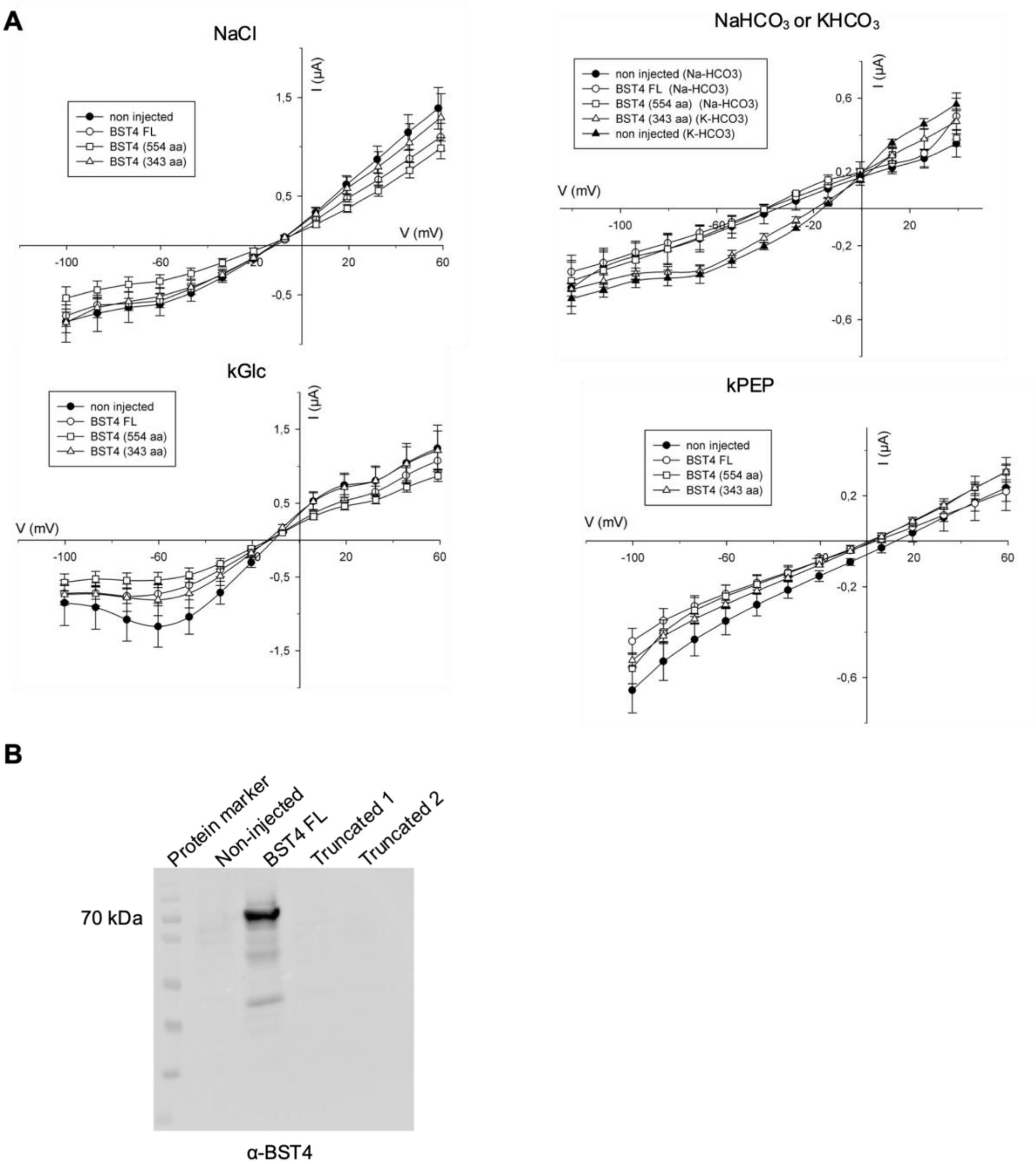


**Supplemental Figure 13. No currents were detected for BST4 with any anions tested in Xenopus oocytes** **A.** Steady state currents analysis of oocytes injected with BST4, full length or truncated, compared to non-injected oocytes. The voltage steps start from 60 mV to -100 mV ($Cl^-$, $PEP^-$, and Gluconate conditions) 40 mV to -120 mV ($HCO3^-$ conditions, right panel), holding voltage is -20 mV. Recordings were performed on n>4 oocytes. There is no differences between the currents recorded in non-injected oocytes and expressing the protein. **B.** Western Blot analysis of oocytes. Lane 1 is the marker, lane 2 are non-injected oocytes, lane 3 is the full-length protein (about 66 kDa), lanes 3 and 4 are the truncated proteins. We can only detect full length BST4 as the antibody is directed against the C-terminal part of the protein that is removed in the two truncated versions of the channel.

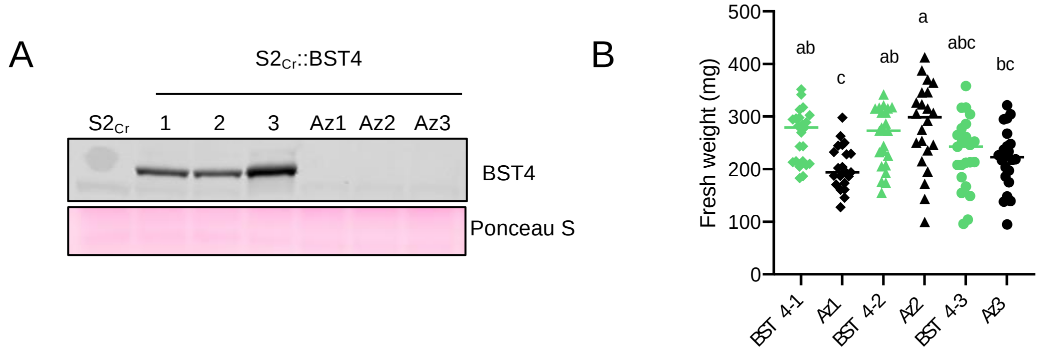


**Supplemental Figure 14. Phenotype of BST4 transgenic Arabidopsis line. A.** Immunoblot against BST4 of proteins extracted from BST4 no tag lines and Azygous segregants (Az). $S2_{Cr}$ is the parent line. **B.** Fresh weight of 28-day old rosettes, bars represent mean weight for each genotype n=18-21. Different letters indicate statistically significant difference among the genotypes (one-way ANOVA test, followed by Turkey's post hoc test, $P < 0.05$).

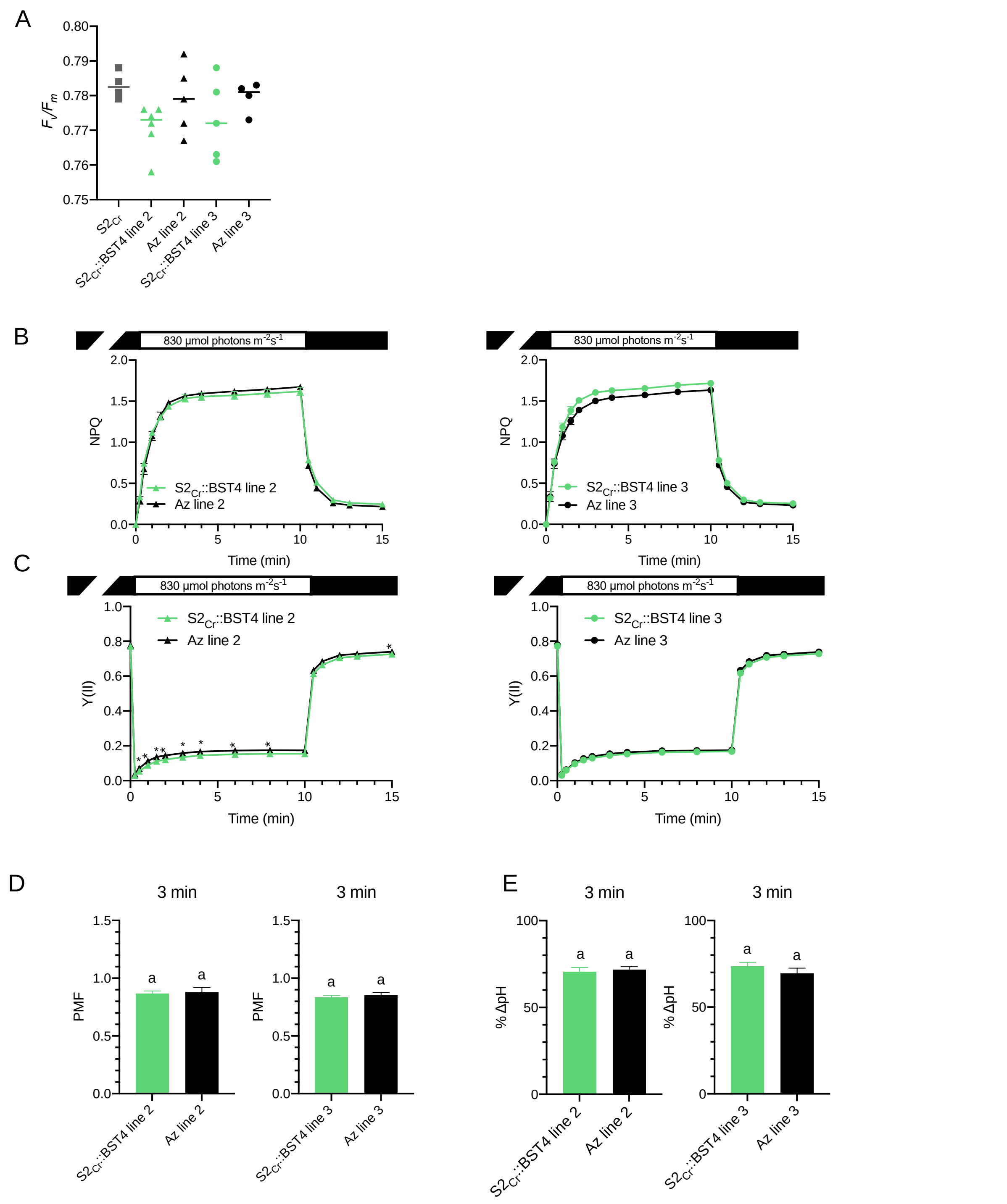


**Supplemental Figure 15. Photosynthetic measurements of BST4 transgenic Arabidopsis lines 2 and 3. A.** $F_v/F_m$ values measured on attached, 30 min dark adapted leaves of 8-week-old plants (n=15-21). The letters in indicate nonsignificant differences between plants expressing BST4 and their azygous segregants using Tukey post hoc test (P > 0.05). **B.** Non-photochemical quenching (NPQ) as a measure of photoprotection and **C.** photosystem II quantum yield (Y(II)) were recorded during 10 min of illumination at 830 µmol photons $m^{-2}$ $s^{-1}$ followed by a 5-minute dark period. Data are presented as means ± SEM (n=4-6). Asterisks indicate statistical difference between plants expressing BST4 and their Azygous (Az) segregants according to unpaired t-test (P ≤ 0.05). **D.** Proton motive force (PMF) size and **E.** partitioning to pH gradient (ΔpH) after 3 min illumination at 830 µmol photons $m^{-2}$ $s^{-1}$. Data are means ± SEM (n=5-6). The letters in E and F indicate non-significant differences between plants expressing BST4 and their azygous segregants according to unpaired t-test (P > 0.05).

##### A. Full length BST4 and homologous sequences

```
>Brassica_rapa_(PAC:30641593)
---------------------------------------------------------------------------
------------------------------------------------------------------------
RTLYTHEKWVEHRSSLRHVHHLFSSFSSRVILSLIPPVFFFTSVAIFIASYNSAVAL-------------
DWLPSVFPILR---SSSLPYQLTAPALALLLVFRTEASYSRYEEGRKAWVGIIAGT--DDLARQVICSVDGSG--
---------------DELVIK-----DLLLRYV----AAFPVALKCHVTYGSD-----VARDLRN-----
LIEGDDLSLI--IESKHR-------------PRCVIEFISQSLQL----LKLDDT--------------
KRDLLESKMLHLHEGIGVCEQLMGIPIPLAYTRL-TSRFLVFWHLTL-------PIILWDEC-----
HWIVVPATFISAASLFCIEEVGVLIEEPFPM----LALDELCDL-VHSNIQEAVKSES-----------------
---------------------------------------------------------------------------
---------------------------------------------------------------------------
---------------------------------------------------------------------------
---------------------------------------------------------------------------
---------------------------------------------------------------------------
-----------------------------------
>Q9M2D2_-_VCCN1_Athaliana
MY------------------QSMNLSVSSNFTHRSLLES------------RFPI-----FSTGFR-----
KSVNLKPPRVSSGPE---SNDSGH---------------ETLTDKLIHLLRAVP--------
DWADEIKERGMQQKRSLYTHEKWVEHRSSLRHVRHLLSSFSSRVILSLIPPVFFFTSVAVVIASYNSAVAL----
---------DWLPGIFPILR---SSSLPYQLTAPALALLLVFRTEASYSRYEEGRKAWVGIIAGT--
NDLARQVICSVDSSG-----------------DELIIK-----DLLLRYI----AAFPVALKCHVIYGSD-----
IARDLRN-----LIEADDLSLI--LQAKHR-------------PRCVIEFISQSIQL----LKLDDA--------
------KRDLLESKMLHLHEGIGVCEQLMGIPIPLSYTRL-TSRFLVFWHLTL-------PIILWDEC-----
HWIVVPATFISAASLFCIEEVGVLIEEPFPM----LALDELCDL-VHSNIQEAVKSEKVIRNRIIAKIKLHEFK-
-----HSS--------------------------NGRHRS-----------------------------------
---------------------------------------------------------------------------
---------------------------------------------------------------------------
---------------------------------------------------------------------------
---------------------------------------------------------------------------
-----------------------------------
>Nicotiana_tomentosiformis_(XP_009629643.1)
M--------------------------TNSRTLFSIQSPTNASFSSHF-TLKTPSK--LQQQSFPSKL-
NFKKLRFSTFKVRCCPQ----QTPQN---------------QNPTSALISILRIIP--------
DWADRIQEEGMKKKRSLYTHESWMQHRSSLRHVRHLFSSLNSRVILSLVPPVIAFTSVAVVIASYNSAVSM----
---------HWLPELFPVLR---ASPLPYQLTAPALALLLVFRTEASYSRFETGKKAWTKVIAGT--
NDFARQVIACVDKS-------------------DAVLK-----AALLQYI----MAFPVALKCHITYGSD-----
IASDLKN-----LLEADDLAVV--LSSKHR-------------PRCIIGFISQCLQS----LHLEGT--------
------KLTQLESKISCFHEGIGVCEQLAGIPIPLSYTRL-TSRFLVLWHLTL-------PIILWDDC-----
HWIVVPATFISAASLFCIEEVGVLIEEPFPM----LALDELCQL-VHDNIQESMANEKKIQERLSAKRKRRFSE-
-----HSQ--------------------------NGWPTS-----------------------------------
---------------------------------------------------------------------------
---------------------------------------------------------------------------
---------------------------------------------------------------------------
---------------------------------------------------------------------------
-----------------------------------
>XP_028961536_Malus_domestica
MLLPSSLSLQTLAPPNADTIQKPHQTLPQNLTLFQLVHP------QHFPTLQFPN-----LPSGPKTL-
KFKLLCSQSPNPNPSP-------PSS---------------SSPVQTLISILRIIP--------
DWSDRTQERGMRQHRTLYDHEKWMHHRSSYRHLRHLLSSLSSRVILSLIPPVIAFTLVAVVIASYNTAVAL----
---------DLLPGIFPLLR---SSSLPYQLTAPALALLLVFRTEASYSRFEEGRKAWTEVIAGA--
NDFARQIISSVETSG------------------DAQLK-----KALLQYI----VAFPVALKCHVIYGSD-----
IARDLQN-----LLEVDDLLVV--LNSKHR-------------PGCIIQFISRSLQL----LKLEES--------
------RRIMLQSKISCFHEGIGICEQLIGTPIPLSYTRL-TSRFLVLWHLTL-------PIILWDDC-----
HWIVVPATFISAASLFCIEQVGVLIEEPFPM----LALDDLCNS-VRNNVQEALASEKLIRARLAAKGRIQSEQ-
-----QFQ--------------------------NGQPRP-----------------------------------
---------------------------------------------------------------------------
---------------------------------------------------------------------------
---------------------------------------------------------------------------
---------------------------------------------------------------------------
```

```
-----------------------------------
>Chlamydomonas_eustigma_(GAX83184.1)
M--------------------------------------------------------------------
LVHRVHTRSLGNRNQCGRK----------------------LHRVSTFVVKTPTEKPVVA----
DYVLPRSEEARRYFRTVYDFPQWQKHRSPTRLIDRLLQIPRSHVLQNILPSIAWCSSVAGLLTLYMQAYDA----
---------HILPDGFPSFATNNACTSFVNTTTVALSLLLVFRTNVSYGRWDEARKMKGLLVNRS--
RDLMRQVCAMVPEE-------------------DVATK-----AMMAKWT----AAFCRVLRIHFQPEVS-----
LEDEMKG-----LLSPEELEWL--IESKHR-------------PCSVIHMLSQIIYD----SQISAI--------
------CQAQMCNNLTAFEDVLGGCERLLRAPIPVSYTRH-TARFLFTWLTLL-------PFALYNSC-----
GVWTLPVVAGVSAVLCGIEEIGVQIEEPFGI----LPLEAICGR-IQADVMATLKEDAKTRNLRNKVLLIRGPE-
-----HWAMV------------------------NGHSNK-----------------------------------
---------------------------------------------------------------------------
---------------------------------------------------------------------------
---------------------------------------------------------------------------
---------------------------------------------------------------------------
-----------------------------------
>Cre16.g662600.t1.2_(BST1)
M---------------------------------QMQA-----------
NRSSLRASPVRGLGARPLLRALPAGRVARLNVSAQA--KDPNAPIQ---------------
SNPLGTLSSQSGQVA----------
TLPRSEEARKYFRTVYDFPQWQKHRSSYRFAERLFQLSQSHILQNALPAISWVTLVATLVASYGYSYDQ------
-------HMLPDVFPSISPNASCTAFISNTSVALSLLLVFRTNSSYGRWDEARKMWGGLLNRS--
RDIMRQGATCFPDD-------------------QVEAK-----KALARWT----VAFSRALRIHFQPEVT-----
IESELQN-----ILTPAELQML--AKSQHR-------------PVRAIHAISQIIQS----VPMSSI--------
------HQQQMSNNLTFFHDVLGGCERLLRAPIPVSYTRH-TARFLFAWLTLL-------PFALYPTT-----
GWGVVPVCTGIAAVLCGIEEIGVQCEEPFGI----LPLDVICNR-
IQADVMATLKDDADTKTILAEAGLISLIPS----ATSATPVASA----EPVLVSARPSAAPAPNNGLQVRVAM--
------------------------------------------------------------------------GG-
--------------------ERK*---------------------------------------------------
---------------------------------------------------------------------------
---------------------------------------------------------------------------
-----------------------------------------------------------------
>Cre16.g663400.t2.1_(BST2)
M---------------------------------QC-------------LSSRPVAMGRAGSSALPRL-
PLRAGRVCHLGVRCQAANKDPNAPIQ---------------SNPLGSFSSQLQNQP----------
TLPRSEEARKYFRTVYDFPQWQTHRNQYRLMKRLFSIPQSHVIQNALPSIMWVAFTSTCVAAYMYGYDQ------
-------HMLPEGFPTLAPNAACSAFISNTSVALSLLLVFRTNSSYGRWDEARKMWGGLLNRS--
RDIMRQGATCFPDD-------------------QVEAK-----KALARWV----VAFSRALRIHFQPEVT-----
IESELKN-----ILTPAELQML--AKSQHR-------------PVRAIHAISQIIQS----VPMSSI--------
------HQQQMSNNLTFFHDVLGGCERLLRAPIPVSYTRH-TARFLFAWLTLL-------PFALYGSC-----
GVSVIPVCSGIAAVLCGIEEIGVQCEEPFGI----LPLDVICNR-IQADVMATLKDDADTKTILAEAGLISLRA-
-----NSAMAVENALPDLDSINAAAPNGNGSHNGNGAAV------------------------------------
-------------------------------------------------------------------PVS-----
----------------------VSAGA------------------------------------------------
---------------------------------------------------------------------------
-----------------------------------------------------------------SGNGMNV---
------------------------RISPR*-----
>Cre16.g663450.t1.2_(BST3)
M---------------------------------QVSK-----------VPSS------ASARCLPRL-
PVRTSRVCQLSVRCQAANKDPNAPIQ---------------SNPLGSFSSQNSSGAVV--------
TAPRNEDARKYFRTVYDFPQWQKHRSQSRLVRRLFTIPQSHVIQNALPSIMWVTFTSTCVAAYMYGYDL------
-------HILPEGFPTLAPNAACSAFISNTSVALSLLLVFRTNSSYGRWDEARKMWGGLLNRS--
RDIMRQGATCFPDD-------------------QVEAK-----KALARWT----VAFARALRIHFQPEVT-----
IESELQN-----ILTPAELQML--AKSQHR-------------PVRAIHAISQIIQS----VRMSSI--------
------HQQQMSNNLTFFHDVLGGCERLLRAPIPVSYTRH-TARFLFAWLTLL-------PFALYGSC-----
GVSVIPVCTGIAAVLCGIEEIGVQCEEPFGI----LPLDVICNR-IQADVMATLKDDADTKTVLAEAGLISLIP-
-----SMSLPPTEHASPSDPVTAAAAAALAAANGNGAASHSNG--------------------------------
------------------------------------------NG---------------------
SKPVSTQVPP-----------------PVLAPVTVTS--------------------------------------
---------------------------------------------------------------------------
---------------------------------------------------------------------------
```

SSGSMNV---------------------------RISPR*-----
>Chlorella_sorokiniana_(PRW33726.1)
M-----------------------STAMLAGSRIQLQQPAG--------LGGSRLQ--RAAAPVAAAA-
RLGSVRPAGLQARSTAARRADRSALR---------------VSATASPEAAPVKLS----------
GDDLKEANRKHMRSVFDFDLWKKHRSSSRYLRHIVGLGESRIVSGLMAPLTYVMTLSLAVACYNAAAEA------
-------GYLP-VFPELKL--ATNAPFGLTSFALSLLLVFRTNSSYGRWDEARKMWGLIVNRS--
RDFIRQGLGYIPPE-------------------QEELQ-----KMLVRWT----VAYSRSLMCHLRPGED-----
LRVELKD-----TLKPEELEAL--LASTHR-------------PNYVVQVLTAIIKT----
AQLPAAVTNNRDSTGCVPAGAAYRMDENLTVFADVTGGCERILRTPIPLSYTRH-TSRFMMIWLTLL-------
PFTLWDSC-----HWAMLPIAGIVSFLLLGIEEIGVQIEEPFTI----LPLEVISRT-IEGNVWELYRMHSGEA-
--LEKEQAELAN------GQDVQVLNA----QDLVALMAPSAVGNTANGTSRKSLV------------------
VNYGL----------------------------------------------------------------------
---------------------------------------------------------------------------
---------------------------------------------------------------------------
---------------------------------------------------------------------------
---------------------------------------------------------------------------
-------------------------------------------------
>Cre06.g261750.t1.2_(BST4)
M-----------------------QC--------QLKH----------------------------GA-
RPQSQRPNWLPARAATLRPAVQHGVR---------------RGLTLGVKAAAAPLE--------
DKKMPADMTTRQYRRVVYDFALWAKHRDVNRYLYNLRTIPGSRIIRQLSQPMGVVLAWAALFGFYETCLEA----
---------GVLPSYLPKMTL--MSAEPQGLTSFALSLLLVFRTNSSYGRFDEARKIWGGILNRA--
RNIANQAVTFIPAE-------------------DQAGR-----EAVGKWT----VGFTRALQAHLQEDID-----
LRKELEKA--TPRWSKEEIDML--VNAQHR-------------PIKAISVLSELTRQ----LSITQF--------
------QALQMQENCTFFYDALGGCERLLRTPIPVSYTRH-TARFLTIWLAML-------PLGLWERY-----
HWSMLPVIALIGFLLLGIDEIGISIEEPFGI----LPLDAICGR-AQTDVNSLLKEDPAVMKYVDDVRSGRVKS-
-----PPPLPPAPA----APAAA-------AAAAAAAARSVSP----
QPDVAKTLGSLFTNVRAGVGAVAPGAPLMPQAPVRSPSPTRSVSP--------SFPRASAGTGMPPPVGMNG---
-------ATPRVAAAPPTPPPVSRPAA------------------
PAAAPAAGSGFTMPNFSASLSGLTGGAAAAAKSAADAASSKLTKMADSMSSGAAAPAPPAAPAR-----------
----------PSTSPRPSASSPISSSADADRSDSSRR---PVNWRDELQSLKATRE----------------
PNGNGNGSGVAPAA--GRADADEEALRRFGNLAGRSR--------------SGNG---------------
GGGSSDTELSEANRPRTRPDWRNQL*
>XP_002945810.1_Volvox_carteri_f._nagariensis
M-----------------------QS--------QLQP----------------------------RL-
QLQGTRLNWLPQRSCVQRRSLRVDAT---------------SG------AAPPPPA--------
GKELSNDMVTRQYRRTVYDFSLWAKHRDVNRYLYNLKTIPGSRIIRTLGQPMGIVLAWAAMFGFYETCLES----
---------GVLPSYFPKLTL--MSAEPQGLTSFALSLLLVFRTNSSYGRFDEARKIWGGILNRA--
RNIANQAVTFIPAE-------------------DVAGR-----EAVGKWA----VGFCRALQAHLQEDAN-----
LREELQKA--QPRWSREEIDML--CSAQHSWQQLQSCVNAFW-PIKAISMLSELTRQ----LPISQF--------
------QALQMQENVTFFYDALGGCERLLRTPIPVSYTRI-----------------------------------
---------------------------------LPLDAICTR-AQTDVVSLLKDDPAVVKYISDVRQGRIAP---
---PTEPPVAGA----
APVAAAPPPPPPASAGGGISRSGSPTAQQQPDVMKTVTSMLHNVKAGIGAVAPAPPRPPSPQPRARSP-
RAASPGGPS----PFPRASAGTG--------G----------
AAAAVPSPPPIKPLTSSSSSSSGAVSKDSNNSTATAKKPASAPAASSA----GFSMGFSGLADGAAAAAKSAS-
AAAAKFSKIADSVVAG--TPAAPASEAKRETAA--AAAMQAQPRN---
TPSSSSSTPSAAPANGSSDDDRSSSGRRTAAAVNWREELAALRAGREDAEEPASASASYDREFPSSSWSFSSASS
AAVVQSGDAEDEARRRFGGLAGRGARSDTTTSAAAVMRGNGNGLSENGYGNGYGNDN-
GNGNGNTVEARGARPRTRPDWRNQL-
>Chrysochromulina_sp._(KOO32217.1)
M-----------------------REHPLSYEEYMRQRS----------AGRDPLA--EAVQGQSASM-
ERGVVPPPPVKATPERTEAAFTPPAEFDFFQDVFKPTVESVVKAVVSPGAQQDSDESYMRVPWWEQGSTYSEDQR
KDRRTVFMHDDWKRHRSSERFFRNIKTWPSSGINQALRKELTFVTSVSVFVVLANMLLYQYQDFGGVVHPGPLSF
LDGPIKSLS---LPALPFSMASPVLSLLLVFRTNTAYFRWNEARTLWGGLINNC--RNIVRQTTTMFPNDA----
-------------YHNALK-----KRLATET----ATFIKSLRNFLRGPEDDAT--
LRKELYAYVNQGLMTSAQAEAT--LAAKNR-------------PMFALAAMSATLRK----ANIDEM--------
------YISRMDSTISVLVDLTGANERIFKSPIPLVYTRL-TARFLSVFLTLL-------
PLAMWAALGESWNHWATIPATFILSVFLFGVEEVGIQIEEPFSI----LPLEAMCNGAIEAVQLEMLAAE-----
------------------QSQVFEAAG----DAVAVA--------------------------------------
---------------------------------------------------------------------------
---------------------------------------------------------------------------

```
---------------------------------------------------------------------------
---------------------------------------------------------------------------
------------------------------------------------
>Emiliania_huxleyi_(XP_005770556.1)
---------------------------------------------------------------------------
--------------------------------------------------------------
EFKEAGREFRQDVYSYNDWRWHRESGHIASAISSVFTSGVGKAMWRETFFVIATAAAVYLYNIGVPVLAAKTAAS
L----PIVAALLGRLPLLHLSLLPLTLSSPALFLLLVFRTNNSYDRWWEARKVWGGVINAS--
RDLARQALALVR---------------------DAELK-----KLMVSQI----
ASYARVLKYHLGPPTPEARDLLRNELVD----NRLPADQVRVI--MEAKHK-------------
PMALLGLVSASLHDSGR-TGLDTV--------------QASKLDQTLSLLTDYLGKCERIVKTPLPLVYTRH-
TARFLSWWLLFL-------PVCLYNQLRA---NWMIVPVSGLIGFFLVGIEDLGNQIEEPFSI----
LPLTAMGTG-IQQSIFEAL--------------------------------------------------------
---------------------------------------------------------------------------
---------------------------------------------------------------------------
---------------------------------------------------------------------------
---------------------------------------------------------------------------
---------------------------------------------------------------------------
>Phaeodactylum_tricornutum_(XP_002180738.1)
MM--------------------------------------------------------RNFASVLLLL-
SSGAAAFAPVQHNGVRTIATPSTPLY---------------GNTKQPPALP--PIK----------
DISYGEESRKYRRTVYSHDDWVKHRSSDRFLRNLLAIGSSGVYKSLAKEVLATTGVATFIVLYNCLVGGYTDLEG
IKHS---ALIESVWAPLMA--LPLAPFTLSSPSLGLLLVFRTNTSYQRWDEARKNWGMNINHT--
RDLVRMGTSFYDNAA-----------------VSSEQRAKDL-KALSLAT----WSFVRAMKRHLSPESEDEQD-
FRRELFE-----RLPAPQAQAI--IDAAHR-------------PNRALFDLSVAIEN----LPMHFL--------
------RKNQVHQAVTIFEDNLGSSERLLTSPVPLFYSRH-TARFLSFWLLLL-------
PFALWDPFAGTWNHVGMIPATAVISIFLFGIEELATQMEEPFTI----LPMQAFCDK-
IGNWCNEIVSWQAGDNGMAVNMPSMISPEGLPELKEPAPVPAMA----VASVAAAMPVMANGDINGDTTGITM--
-DQP-------------------------------------------------HNAIP-----------------
---------------------------------------------------------------------------
---------------------------------------------------------------------------
---------------------------------------------------------------------------
-----------------------------------------------------------------
>Thalassiosira_pseudonana_(XP_002289965.1)
M--------------------------------------------------------------------------
---------------------------------------GPPIDPSVPVT-----------
DQVGEGSRKYRRTVYTHDDWVRHRSPDRFGNNLSTLFNSGIYKQVANEVFATTAVATFVFLWNMIAGGYTDLAGV
QHG---PIIDSPLAQMVG--LPMTAFTILTPSLGLLLVFRTNTSYGRWDEARKMWGLNINHT--
RDLNRMATAWYGNEGNMDSVAFMGGDIPYSQPIDPVQRAYDL-GQVSLFT----WAFVRSMKRHLSPPEEDEED-
FKAELRA-----RLTPEQAENI--INAAHR-------------PNRALFDLSVAIEN----LPMHFL--------
------RKNAINTNLSIFEDTLGGCERLLSSPVPLFYSRH-TARFLSTWLLLL-------
PFGLYEQFKDSWNHIAMIPATAFISVCLFGIEELATQLEEPFTI----LPMQGFCDK-IGGWCDEIVSW-
AGQGQQEYTEENAMSNE------QEMTY-----------------------------------------------
---------------------------------------------------------------------------
---------------------------------------------------------------------------
---------------------------------------------------------------------------
---------------------------------------------------------------------------
---WR----------------------------------------------------------------------
-----------------------------------------------------
>WP_049046555.1_Kiebsiella_aerogenes
---------------------------------------------------------------------------
---------------------------------------------------------------------------
------MIIRPEQHWFFRLFDWHGSVLSKIVFRLLLNVLMSVIAIISYQWYEQL-------------GI------
-----HLTVAPFSLLGIAIAIFLGFRNSASYSRFVEARNLWGTVLIAE--RTLVRQLKNILPD------------
--------DEETH-----KTLVSYL----VAFSWSLKHQLRK-TD-----PAVDLYR-----LLPKEKVAEI--
LASSMP-------------TNRILLLIGNELGRLREQGKLSDI--------------
TYGLMDNKLDELAHVLGGCERLASTPVPFAYTLI-LQRTVYLFCTLL-------PFALVGDL-----
HYMTPFVSVFISYTFLSWDSLAEELEDPFGTSANDLPLNAMCNT-IERNLMDMTGQHPLPEKMQPDRYYNLT---
---------------------------------------------------------------------------
---------------------------------------------------------------------------
---------------------------------------------------------------------------
---------------------------------------------------------------------------
```

---------------------------------------------------------------------------
-----------------------------------
>XP_011543531.1_Homo_sapiens
M--------------------------------------------------------------------------
--------------------------------------------------------------------------
TITYTSQVANARLGSFSRLLLCWRGSIYKLLYGEFLIFLLCYYIIRFIYRLALTEEQQL--------------
MFEKLTL--YCDSYIQL--IPISFVLGFYVTLVVTRW------WNQYENLPWPDRLMSLVSGFVEGK--------
-----------DEQGRL--LRRTLIRYANLGNVLILRSVSTAVYKRFP-----SAQHLVQ---
AGFMTPAEHKQLEKLSLPHNM---------FWVPWVWFANLSMKAWLG---GRIRDPI-------------
LLQSLLNEMNTLRTQCGHLYAYDWISIPLVYTQVVTVAVYSFFLTCLVGRQFLNPAKAYPGHEL---
DLVVPVFTFLQFFFYVGWLKVAEQLINPFGEDDDDFETNWIVDRNLQVSLLAVDEMHQDLPRMEPDMYWNKPEP-
-----QPPYTAASA----QFRRASFMGSTFNISLNKEEMEFQPNQEDEED-----------
AHAGIIGRFLGLQSHDHHPPRANSRTKLLWPKRESLLHEGLPKNHKAAKQ----
NVRGQEDNKAWKLKAVDAFKSAPLYQRPGYYSAPQ-----------TPLSPTPMFFPLEPSA---
PSKLHSVTGI-DTKDKSLKTVSSGAKKSFELLSES--
DGALMEHPEVSQVRRKTVEFNLTDMPEIPENHLKEPLEQSPTNIHTTLKDHMDPYWALENR----------
SVLHLNQGHCIALCPTPASLALSLPFLHNFLGFHHCQSTLDLRPALAWGIYLATFTGILGKC-------------
--SGPFLTSPWY---HPEDFLGPGEGR--------------------

**B. BST4 and homologous sequences trimmed after the bestrophin domain**

>Brassica_rapa_(PAC:30641593)
---------------------------------------------------------------------------
------------------------------------------------------------------------
RTLYTHEKWVEHRSSLRHVHHLFSSFSSRVILSLIPPVFFFTSVAIFIASYNSAVAL-------------
DWLPSVFPILR---SSSLPYQLTAPALALLLVFRTEASYSRYEEGRKAWVGIIAGT--DDLARQVICSVDGSG--
---------------DELVIK-----DLLLRYV----AAFPVALKCHVTYGSD-----VARDLRN-----
LIEGDDLSLI--IESKHR-------------PRCVIEFISQSLQ----LKLDDT---------------
KRDLLESKMLHLHEGIGVCEQLMGIPIPLAYTRL-TSRFLVFWHLTL-------PIILWDEC-----
HWIVVPATFISAASLFCIEEVGVLIEEPFPM----LALDELCDL-VHSNIQEAVKSE
>Q9M2D2_-_VCCN1_Athaliana
MY-----------------QSMNLSVSSNFTHRSLLES-------------RFPI-----FSTGFR-----
KSVNLKPPRVSSGPE---SNDSGH---------------ETLTDKLIHLLRAVP--------
DWADEIKERGMQQKRSLYTHEKWVEHRSSLRHVRHLLSSFSSRVILSLIPPVFFFTSVAVVIASYNSAVAL----
---------DWLPGIFPILR---SSSLPYQLTAPALALLLVFRTEASYSRYEEGRKAWVGIIAGT--
NDLARQVICSVDSSG-----------------DELIIK-----DLLLRYI----AAFPVALKCHVIYGSD-----
IARDLRN-----LIEADDLSLI--LQAKHR-------------PRCVIEFISQSIQ----LKLDDA---------
------KRDLLESKMLHLHEGIGVCEQLMGIPIPLSYTRL-TSRFLVFWHLTL-------PIILWDEC-----
HWIVVPATFISAASLFCIEEVGVLIEEPFPM----LALDELCDL-VHSNIQEAVKSE
>Nicotiana_tomentosiformis_(XP_009629643.1)
M--------------------------TNSRTLFSIQSPTNASFSSHF-TLKTPSK--LQQQSFPSKL-
NFKKLRFSTFKVRCCPQ----QTPQN---------------QNPTSALISILRIIP--------
DWADRIQEEGMKKKRSLYTHESWMQHRSSLRHVRHLFSSLNSRVILSLVPPVIAFTSVAVVIASYNSAVSM----
---------HWLPELFPVLR---ASPLPYQLTAPALALLLVFRTEASYSRFETGKKAWTKVIAGT--
NDFARQVIACVDKS-------------------DAVLK-----AALLQYI----MAFPVALKCHITYGSD-----
IASDLKN-----LLEADDLAVV--LSSKHR-------------PRCIIGFISQCLQS----LHLEGT--------
------KLTQLESKISCFHEGIGVCEQLAGIPIPLSYTRL-TSRFLVLWHLTL-------PIILWDDC-----
HWIVVPATFISAASLFCIEEVGVLIEEPFPM----LALDELCQL-VHDNIQESMANE
>XP_028961536_Malus_domestica
MLLPSSLSLQTLAPPNADTIQKPHQTLPQNLTLFQLVHP------QHFPTLQFPN-----LPSGPKTL-
KFKLLCSQSPNPNPSP-------PSS---------------SSPVQTLISILRIIP--------
DWSDRTQERGMRQHRTLYDHEKWMHHRSSYRHLRHLLSSLSSRVILSLIPPVIAFTLVAVVIASYNTAVAL----
---------DLLPGIFPLLR---SSSLPYQLTAPALALLLVFRTEASYSRFEEGRKAWTEVIAGA--
NDFARQIISSVETSG------------------DAQLK-----KALLQYI----VAFPVALKCHVIYGSD-----
IARDLQN-----LLEVDDLLVV--LNSKHR-------------PGCIIQFISRSLQL----LKLEES--------
------RRIMLQSKISCFHEGIGICEQLIGTPIPLSYTRL-TSRFLVLWHLTL-------PIILWDDC-----
HWIVVPATFISAASLFCIEQVGVLIEEPFPM----LALDDLCNS-VRNNVQEALASE
>Chlamydomonas_eustigma_(GAX83184.1)
M--------------------------------------------------------------------

```
LVHRVHTRSLGNRNQCGRK----------------------LHRVSTFVVKTPTEKPVVA----
DYVLPRSEEARRYFRTVYDFPQWQKHRSPTRLIDRLLQIPRSHVLQNILPSIAWCSSVAGLLTLYMQAYDA----
---------HILPDGFPSFATNNACTSFVNTTTVALSLLLVFRTNVSYGRWDEARKMKGLLVNRS--
RDLMRQVCAMVPEE-------------------DVATK-----AMMAKWT----AAFCRVLRIHFQPEVS-----
LEDEMKG-----LLSPEELEWL--IESKHR-------------PCSVIHMLSQIIYD----SQISAI--------
------CQAQMCNNLTAFEDVLGGCERLLRAPIPVSYTRH-TARFLFTWLTLL-------PFALYNSC-----
GVWTLPVVAGVSAVLCGIEEIGVQIEEPFGI----LPLEAICGR-IQADVMATLKED
>Cre16.g662600.t1.2_(BST1)
M---------------------------------QMQA-----------
NRSSLRASPVRGLGARPLLRALPAGRVARLNVSAQA--KDPNAPIQ---------------
SNPLGTLSSQSGQVA----------
TLPRSEEARKYFRTVYDFPQWQKHRSSYRFAERLFQLSQSHILQNALPAISWVTLVATLVASYGYSYDQ------
-------HMLPDVFPSISPNASCTAFISNTSVALSLLLVFRTNSSYGRWDEARKMWGGLLNRS--
RDIMRQGATCFPDD-------------------QVEAK-----KALARWT----VAFSRALRIHFQPEVT-----
IESELQN-----ILTPAELQML--AKSQHR-------------PVRAIHAISQIIQS----VPMSSI--------
------HQQQMSNNLTFFHDVLGGCERLLRAPIPVSYTRH-TARFLFAWLTLL-------PFALYPTT-----
GWGVVPVCTGIAAVLCGIEEIGVQCEEPFGI----LPLDVICNR-IQADVMATLKDD
>Cre16.g663400.t2.1_(BST2)
M---------------------------------QC-------------LSSRPVAMGRAGSSALPRL-
PLRAGRVCHLGVRCQAANKDPNAPIQ---------------SNPLGSFSSQLQNQP----------
TLPRSEEARKYFRTVYDFPQWQTHRNQYRLMKRLFSIPQSHVIQNALPSIMWVAFTSTCVAAYMYGYDQ------
-------HMLPEGFPTLAPNAACSAFISNTSVALSLLLVFRTNSSYGRWDEARKMWGGLLNRS--
RDIMRQGATCFPDD-------------------QVEAK-----KALARWV----VAFSRALRIHFQPEVT-----
IESELKN-----ILTPAELQML--AKSQHR-------------PVRAIHAISQIIQS----VPMSSI--------
------HQQQMSNNLTFFHDVLGGCERLLRAPIPVSYTRH-TARFLFAWLTLL-------PFALYGSC-----
GVSVIPVCSGIAAVLCGIEEIGVQCEEPFGI----LPLDVICNR-IQADVMATLKDD
>Cre16.g663450.t1.2_(BST3)
M---------------------------------QVSK-----------VPSS------ASARCLPRL-
PVRTSRVCQLSVRCQAANKDPNAPIQ---------------SNPLGSFSSQNSSGAVV--------
TAPRNEDARKYFRTVYDFPQWQKHRSQSRLVRRLFTIPQSHVIQNALPSIMWVTFTSTCVAAYMYGYDL------
-------HILPEGFPTLAPNAACSAFISNTSVALSLLLVFRTNSSYGRWDEARKMWGGLLNRS--
RDIMRQGATCFPDD-------------------QVEAK-----KALARWT----VAFARALRIHFQPEVT-----
IESELQN-----ILTPAELQML--AKSQHR-------------PVRAIHAISQIIQS----VRMSSI--------
------HQQQMSNNLTFFHDVLGGCERLLRAPIPVSYTRH-TARFLFAWLTLL-------PFALYGSC-----
GVSVIPVCTGIAAVLCGIEEIGVQCEEPFGI----LPLDVICNR-IQADVMATLKDD
>Chlorella_sorokiniana_(PRW33726.1)
M-----------------------STAMLAGSRIQLQQPAG--------LGGSRLQ--RAAAPVAAAA-
RLGSVRPAGLQARSTAARRADRSALR---------------VSATASPEAAPVKLS----------
GDDLKEANRKHMRSVFDFDLWKKHRSSSRYLRHIVGLGESRIVSGLMAPLTYVMTLSLAVACYNAAAEA------
-------GYLP-VFPELKL--ATNAPFGLTSFALSLLLVFRTNSSYGRWDEARKMWGLIVNRS--
RDFIRQGLGYIPPE-------------------QEELQ-----KMLVRWT----VAYSRSLMCHLRPGED-----
LRVELKD-----TLKPEELEAL--LASTHR-------------PNYVVQVLTAIIKT----
AQLPAAVTNNRDSTGCVPAGAAYRMDENLTVFADVTGGCERILRTPIPLSYTRH-TSRFMMIWLTLL-------
PFTLWDSC-----HWAMLPIAGIVSFLLLGIEEIGVQIEEPFTI----LPLEVISRT-IEGNVWELYRMH
>Cre06.g261750.t1.2_(BST4)
M-----------------------QC--------QLKH----------------------------GA-
RPQSQRPNWLPARAATLRPAVQHGVR---------------RGLTLGVKAAAAPLE--------
DKKMPADMTTRQYRRVVYDFALWAKHRDVNRYLYNLRTIPGSRIIRQLSQPMGVVLAWAALFGFYETCLEA----
---------GVLPSYLPKMTL--MSAEPQGLTSFALSLLLVFRTNSSYGRFDEARKIWGGILNRA--
RNIANQAVTFIPAE-------------------DQAGR-----EAVGKWT----VGFTRALQAHLQEDID-----
LRKELEKA--TPRWSKEEIDML--VNAQHR-------------PIKAISVLSELTRQ----LSITQF--------
------QALQMQENCTFFYDALGGCERLLRTPIPVSYTRH-TARFLTIWLAML-------PLGLWERY-----
HWSMLPVIALIGFLLLGIDEIGISIEEPFGI----LPLDAICGR-AQTDVNSLLKED
>XP_002945810.1_Volvox_carteri_f._nagariensis
M-----------------------QS--------QLQP----------------------------RL-
QLQGTRLNWLPQRSCVQRRSLRVDAT---------------SG------AAPPPPA--------
GKELSNDMVTRQYRRTVYDFSLWAKHRDVNRYLYNLKTIPGSRIIRTLGQPMGIVLAWAAMFGFYETCLES----
---------GVLPSYFPKLTL--MSAEPQGLTSFALSLLLVFRTNSSYGRFDEARKIWGGILNRA--
RNIANQAVTFIPAE-------------------DVAGR-----EAVGKWA----VGFCRALQAHLQEDAN-----
LREELQKA--QPRWSREEIDML--CSAQHSWQQLQSCVNAFW-PIKAISMLSELTRQ----LPISQF--------
------QALQMQENVTFFYDALGGCERLLRTPIPVSYTRI-----------------------------------
```

```
---------------------------------LPLDAICTR-AQTDVVSLLKDD
>Chrysochromulina_sp._(KOO32217.1)
M-----------------------REHPLSYEEYMRQRS----------AGRDPLA--EAVQGQSASM-
ERGVVPPPPVKATPERTEAAFTPPAEFDFFQDVFKPTVESVVKAVVSPGAQQDSDESYMRVPWWEQGSTYSEDQR
KDRRTVFMHDDWKRHRSSERFFRNIKTWPSSGINQALRKELTFVTSVSVFVVLANMLLYQYQDFGGVVHPGPLSF
LDGPIKSLS---LPALPFSMASPVLSLLLVFRTNTAYFRWNEARTLWGGLINNC--RNIVRQTTTMFPNDA----
-------------YHNALK-----KRLATET----ATFIKSLRNFLRGPEDDAT--
LRKELYAYVNQGLMTSAQAEAT--LAAKNR-------------PMFALAAMSATLRK----ANIDEM--------
------YISRMDSTISVLVDLTGANERIFKSPIPLVYTRL-TARFLSVFLTLL-------
PLAMWAALGESWNHWATIPATFILSVFLFGVEEVGIQIEEPFSI----LPLEAMCNGAIEAVQLEMLAAE
>Emiliania_huxleyi_(XP_005770556.1)
---------------------------------------------------------------------------
--------------------------------------------------------------
EFKEAGREFRQDVYSYNDWRWHRESGHIASAISSVFTSGVGKAMWRETFFVIATAAAVYLYNIGVPVLAAKTAAS
L----PIVAALLGRLPLLHLSLLPLTLSSPALFLLLVFRTNNSYDRWWEARKVWGGVINAS--
RDLARQALALVR---------------------DAELK-----KLMVSQI----
ASYARVLKYHLGPPTPEARDLLRNELVD----NRLPADQVRVI--MEAKHK-------------
PMALLGLVSASLHDSGR-TGLDTV--------------QASKLDQTLSLLTDYLGKCERIVKTPLPLVYTRH-
TARFLSWWLLFL-------PVCLYNQLRA---NWMIVPVSGLIGFFLVGIEDLGNQIEEPFSI----
LPLTAMGTG-IQQSIFEAL---
>Phaeodactylum_tricornutum_(XP_002180738.1)
MM--------------------------------------------------------RNFASVLLLL-
SSGAAAFAPVQHNGVRTIATPSTPLY---------------GNTKQPPALP--PIK----------
DISYGEESRKYRRTVYSHDDWVKHRSSDRFLRNLLAIGSSGVYKSLAKEVLATTGVATFIVLYNCLVGGYTDLEG
IKHS---ALIESVWAPLMA--LPLAPFTLSSPSLGLLLVFRTNTSYQRWDEARKNWGMNINHT--
RDLVRMGTSFYDNAA-----------------VSSEQRAKDL-KALSLAT----WSFVRAMKRHLSPESEDEQD-
FRRELFE-----RLPAPQAQAI--IDAAHR-------------PNRALFDLSVAIEN----LPMHFL--------
------RKNQVHQAVTIFEDNLGSSERLLTSPVPLFYSRH-TARFLSFWLLLL-------
PFALWDPFAGTWNHVGMIPATAVISIFLFGIEELATQMEEPFTI----LPMQAFCDK-IGNWCNEIVSWQ
>Thalassiosira_pseudonana_(XP_002289965.1)
M--------------------------------------------------------------------------
---------------------------------------GPPIDPSVPVT-----------
DQVGEGSRKYRRTVYTHDDWVRHRSPDRFGNNLSTLFNSGIYKQVANEVFATTAVATFVFLWNMIAGGYTDLAGV
QHG---PIIDSPLAQMVG--LPMTAFTILTPSLGLLLVFRTNTSYGRWDEARKMWGLNINHT--
RDLNRMATAWYGNEGNMDSVAFMGGDIPYSQPIDPVQRAYDL-GQVSLFT----WAFVRSMKRHLSPPEEDEED-
FKAELRA-----RLTPEQAENI--INAAHR-------------PNRALFDLSVAIEN----LPMHFL--------
------RKNAINTNLSIFEDTLGGCERLLSSPVPLFYSRH-TARFLSTWLLLL-------
PFGLYEQFKDSWNHIAMIPATAFISVCLFGIEELATQLEEPFTI----LPMQGFCDK-IGGWCDEIVSW-
>WP_049046555.1_Kiebsiella_aerogenes
---------------------------------------------------------------------------
---------------------------------------------------------------------------
------MIIRPEQHWFFRLFDWHGSVLSKIVFRLLLNVLMSVIAIISYQWYEQL-------------GI------
-----HLTVAPFSLLGIAIAIFLGFRNSASYSRFVEARNLWGTVLIAE--RTLVRQLKNILPD------------
--------DEETH-----KTLVSYL----VAFSWSLKHQLRK-TD-----PAVDLYR-----LLPKEKVAEI--
LASSMP-------------TNRILLLIGNELGRLREQGKLSDI--------------
TYGLMDNKLDELAHVLGGCERLASTPVPFAYTLI-LQRTVYLFCTLL-------PFALVGDL-----
HYMTPFVSVFISYTFLSWDSLAEELEDPFGTSANDLPLNAMCNT-IERNLMDMTGQH
>XP_011543531.1_Homo_sapiens
M--------------------------------------------------------------------------
--------------------------------------------------------------------------
TITYTSQVANARLGSFSRLLLCWRGSIYKLLYGEFLIFLLCYYIIRFIYRLALTEEQQL--------------
MFEKLTL--YCDSYIQL--IPISFVLGFYVTLVVTRW------WNQYENLPWPDRLMSLVSGFVEGK--------
-----------DEQGRL--LRRTLIRYANLGNVLILRSVSTAVYKRFP-----SAQHLVQ---
AGFMTPAEHKQLEKLSLPHNM---------FWVPWVWFANLSMKAWLG---GRIRDPI-------------
LLQSLLNEMNTLRTQCGHLYAYDWISIPLVYTQVVTVAVYSFFLTCLVGRQFLNPAKAYPGHEL---
DLVVPVFTFLQFFFYVGWLKVAEQLINPFGEDDDDFETNWIVDRNLQVSLLAVDEMH
```

**Supplemental Figure 16. Sequences used to generate MAFFT alignments used to generate the phylogenetic trees included in this manuscript.** FASTA files of MAFFT alignments of **A.** Full length and **B.** trimmed after the bestrophin domain BST4 and homologous amino acid sequences.
